# Endogenous PROM1 activation redistributes liver cancer cells along a preexisting differentiation axis

**DOI:** 10.64898/2026.11.01.763386

**Authors:** Tom Schneider, Kota Kaneko, Marion Vogl, Kirsten Utpatel, Alexander Immel, Daniela Hirsch, Chiara Raggi, Elena Spinola Lasso, Matthias Evert, Gen-Sheng Feng, Diego F. Calvisi, Alexander Scheiter

## Abstract

CD133 is widely used as a marker of stem-like cells in hepatocellular carcinoma, yet its functional contribution remains incompletely defined. In PLC/PRF/5 cells, acute CRISPR activation of endogenous PROM1 suppresses hepatic programs, whereas PROM1 knockdown elicits an opposing hepatocyte-identity response, together defining a PROM1-responsive transcriptional axis. Single-cell profiling shows that this axis is preexisting and tracks endogenous PROM1 heterogeneity, separating a hepatocyte-like, PROM1-low compartment from a PROM1-enriched, less hepatocyte-like compartment. Sustained PROM1 activation reduces the hepatocyte-like compartment and shifts the population toward intermediate positions along this axis. Related transcriptional patterns are detectable in an independent HuH-7 dataset and a human HCC case. Endogenous ALFA tagging identifies candidate CD133-associated proteins. Together, these findings support CD133 as an active contributor to tumor-cell state redistribution, extending its role beyond a mere marker of stem-like populations.

## Introduction

Hepatocellular carcinoma (HCC) contains malignant cells with different degrees of hepatocyte identity, proliferative activity and responsiveness to environmental signals. Single-cell studies have resolved this variation within tumors and across patients, while experimental studies increasingly identify regulatory mechanisms that maintain or alter these states. A central question is therefore how an observable state marker relates to the mechanisms that change the distribution of states within a tumor-cell population.^1–3^

CD133, encoded by PROM1, is a pentaspan membrane glycoprotein widely used to enrich for stem-like populations. In HCC, early isolation experiments demonstrated increased tumorigenic and self-renewal properties in CD133-positive cells, while it was acknowledged that marker positivity alone cannot specify a universal cancer stem-cell identity or establish how that identity arose.^4–6^ Indeed, a variety of effector roles for CD133 have since been established, including regulation of EGFR–AKT signaling^7^ and a TM4SF5-associated circuit in HCC,^8^ while more recently CD133 was linked to MYH9–β-catenin signaling and tumor growth in MASLD-related HCC.^9^

Zhou et al.^10^ in addition provided a particularly informative in vivo framework. Using Prom1-directed lineage tracing in fibrotic and steatotic mouse HCC models, the authors demonstrated clonal expansion, tumor propagation and phenotypic plasticity of Prom1-marked cells. Single-cell analysis linked progression within the traced population to declining hepatic differentiation programs, and targeted ablation of the marked cells impaired tumor growth. These results established the importance of a Prom1-marked lineage in vivo and connected its heterogeneity to successive dedifferentiation.

These observations connect CD133-associated tumor propagation with changes in hepatocellular differentiation. Recent studies show that hepatocyte identity is actively maintained and can be altered by perturbing its regulatory networks. PROX1 preserves hepatocyte identity by repressing alternative cell-fate programs, while the FOXM1–CEBPB circuit regulates differentiation-associated heterogeneity in HCC.^3,11^ These findings provide a basis for examining CD133 in relation to the balance between more hepatocyte-like and less differentiated cellular states. Although CD133-dependent signaling and tumor-promoting functions have been established, how changes in endogenous PROM1 expression affect the organization of these states within an existing tumor-cell population remains incompletely understood. In particular, whether PROM1 perturbation engages the transcriptional programs underlying baseline heterogeneity and shifts the relative representation of the corresponding states requires direct investigation.

PROM1 uses alternative promoters and splicing to generate multiple transcript and protein isoforms.^12,13^ CRISPR activation (CRISPRa) employs catalytically inactive Cas9 to recruit transcriptional activators to endogenous promoters, implemented here with the synergistic activation mediator (SAM)-based LentiSAMPHv2 construct.^14,15^ Unlike plasmid overexpression of a single cloned CD133 cDNA, this approach can increase multiple endogenous splice isoforms arising from the targeted promoter while retaining cellular RNA processing, making it particularly useful for investigating PROM1 biology.

Here, we combine CRISPR activation of endogenous PROM1 with RNA interference and bulk and single-cell transcriptomics in PLC/PRF/5 cells to examine this relationship. By mapping acute transcriptional responses onto baseline cellular heterogeneity and profiling populations after sustained activation, we identify a preexisting differentiation-associated axis whose state distribution can be experimentally altered. PROM1 activation reduces the relative representation of a hepatocyte-like, PROM1-low compartment, while knockdown opposes the activation-associated transcriptional response. We further examine these programs in an independent HuH-7 dataset and human HCC samples and use endogenous ALFA tagging to identify candidate CD133-associated proteins. Together, these approaches connect PROM1 perturbation to the redistribution of existing cellular states and support a contribution of CD133 to the regulation of differentiation-associated heterogeneity.

## Results

### Acute CRISPR activation increases endogenous PROM1 without detectable enrichment of predicted off-target activation

SAM CRISPRa increased endogenous PROM1 expression in PLC/PRF/5 cells using two co-delivered guide-bearing vectors and guide-free SAMph controls. RNA sequencing showed a 3.6-fold increase (adjusted p = 1.2 × 10^−1^^4^; three wells/group), confirmed by TaqMan qPCR (4.9-fold; six wells/condition across two independent transfections) and CD133 immunoblotting (5.7-fold, adjusted p = 0.010; three wells/condition in one experiment). Immunofluorescence illustrated enhanced membrane-associated staining (Figures 1, S1; Data S1 and S2).

**Figure 1.**
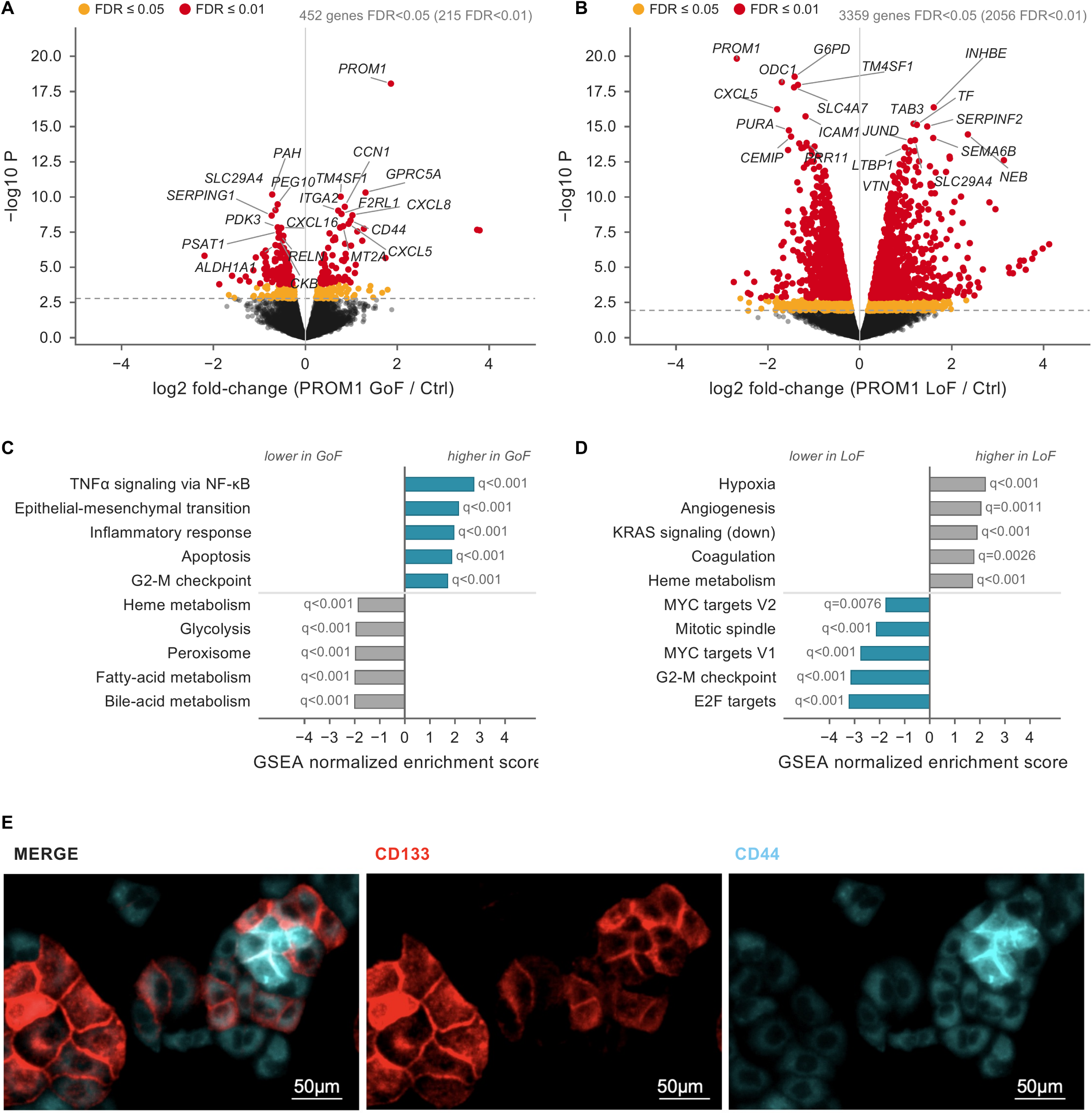
Acute PROM1 gain and loss exert opposing effects on hepatocyte identity and membrane-associated programmes. (A and B) Bulk RNA-seq 48 h after transient *PROM1* activation (CRISPRa versus spacer-less SAMPH control, A) or knockdown (siPOOL versus matched control pool, B). Log2 fold change versus −log10 p; amber, FDR < 0.05; red, FDR < 0.01; dashed line, p at FDR = 0.05. Labels identify the ten strongest genes per direction with |log2 fold change| > 0.5. (C and D) Hallmark enrichment for the activation (C) and knockdown (D) arm. Bars, normalized enrichment score (NES); five of 49 sets per direction with largest |NES| at FDR < 0.05; labels, q values. Teal, high-*PROM1* programs; gray, low-*PROM1*. (E) CD133 (red) and CD44 (cyan) immunofluorescence; merge at left, single channels at center and right. Qualitative. Scale bars, 50 µm. (A–D) n = 3 independently transfected wells per group and arm, one library each. Statistics: two-sided limma-voom moderated t test (A, B); preranked GSEA (C, D); Benjamini–Hochberg correction. See STAR Methods, Table S1 and Figure S1.

Consistent with amplification of existing endogenous promoter activity, transcription increased from the proximal first exon already used in controls, while alternative upstream first exons remained essentially inactive (Figures S1A and S1B). The local response was confined to PROM1: among PROM1 and seven expressed neighbouring genes within ±1 Mb, only PROM1 fell outside its expression-matched expected range (Figure S1C; Table S1).

Predicted off-target candidates with up to four guide mismatches showed no enrichment for activation (265 expressed genes; permutation p = 0.45). Of 249 evaluated for differential expression, eight (3.2%) were differentially expressed, matching the genome-wide background rate. All eight were associated exclusively with four-mismatch sites. A separate promoter-similarity analysis detected a very weak positive correlation with expression changes (Spearman ρ = 0.0534; empirical p = 0.0005; 15,704 genes). These results support selective endogenous PROM1 activation, although they do not establish direct dCas9 binding or exclude distal effects (Table S1).

### Acute PROM1 gain and loss exert opposing effects on hepatocyte identity and membrane-associated programmes

Acute PROM1 activation altered 452 genes (151 increased, 301 decreased; adjusted p < 0.05; Figure 1A; Table S1). Increased transcripts included membrane- and invasion-associated genes, whereas hepatic metabolic and secretory genes and the hepatocyte transcription factors HNF4A, ONECUT1 and PROX1 decreased. qPCR confirmed increases in CD44, KRT80, GPRC5A, PLAU, CXCR4 and SFRP1; immunoblotting supported increases in CD44, KRT80 and the combined upper CXCR4-immunoreactive bands (Figures S1D and S1F). CD44 immunofluorescence revealed marked cell-to-cell heterogeneity (Figure 1E). Hallmark analysis showed negative enrichment of hepatic metabolic programmes, including bile acid metabolism (NES −2.00), and positive enrichment of TNFα signalling via NF-κB (+2.76), EMT (+2.14), inflammatory response, apoptosis and G2M checkpoint (Figure 1C; Table S1).

siPOOL-mediated knockdown reduced PROM1 RNA by 84% in sequencing and 87% in qPCR, with protein near background (Figures 1B, S1E and S1G). It altered 3,359 genes (1,723 increased, 1,636 decreased), with negative enrichment of E2F, G2M and MYC programmes and positive enrichment of hypoxia, angiogenesis, coagulation and heme metabolism (Figure 1D; Table S1). SFRP1, GPRC5A, KRT80 and PLAU decreased, whereas CD44 and CXCR4 remained near control levels (Figure S1E). Thus, knockdown elicited a broader response rather than uniformly reversing activation.

Nevertheless, inferred HNF1A, HNF4A, ONECUT1 and FOXO1 regulon activities decreased after activation and increased after knockdown. HNF1A and HNF4A ranked among the top three regulons by absolute ULM enrichment score in both perturbations, highlighting their prominent reciprocal response (Table S1). Among 124 genes significant in both perturbations at adjusted p < 0.01, 117 (94.4%) changed in opposite directions (Table S1). Of these, 38 increased with activation and decreased with knockdown, including PROM1, membrane/invasion-associated genes, keratins, VIM and MAPK feedback regulators. The other 79 showed the inverse pattern and included hepatic genes, HNF4A, ONECUT1, PROX1 and the oncofetal/HCC-associated markers GPC3 and PEG10. This reciprocal component associated PROM1 activation with reduced hepatocyte identity and increased membrane/invasion-associated expression. Because bulk measurements could not resolve the cellular distribution of these programmes, we next used single-cell profiling to examine pre-existing heterogeneity and population changes under sustained PROM1 activation.

### PLC/PRF/5 control cultures are organised along a PROM1-associated transcriptional axis

Control (SAMph) and PROM1-CRISPRa cells, generated by lentiviral transduction, were each co-cultured with GFP-labelled bystander cells before single-cell profiling (two pools per condition). Bystanders were included to detect non-cell-autonomous transcriptional changes; they showed no clear response to PROM1-activated neighbours (not shown) and were excluded. We first examined 6,589 control cells.

Control cells contained a PROM1-low compartment, corresponding to cluster C0 (35.0% of cells): only 15.9% of these cells had detectable PROM1, compared with 70.0% of the remaining cells, and PROM1-positive cells within it expressed less PROM1 (median 0.63–0.66 versus 0.97 log-normalised expression; Figures 2A and 2B). The reciprocal signature from the acute experiment followed this endogenous distribution: its score (34 activation-upregulated and 73 activation-downregulated genes, excluding PROM1) formed a gradient from C0 towards the PROM1-enriched compartment (Figure S2H; Table S2).

**Figure 2.**
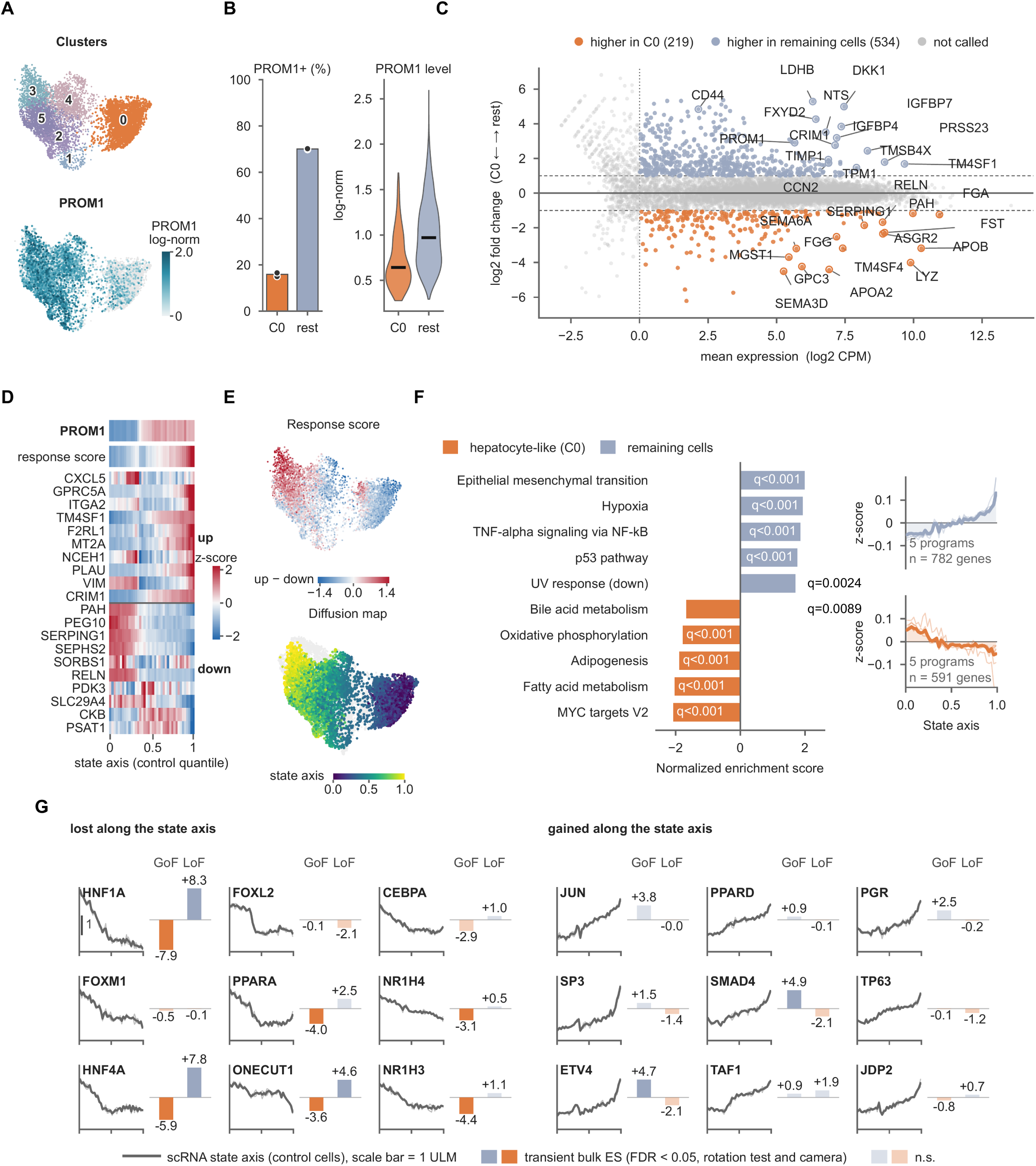
PLC/PRF/5 control cultures are organised along a PROM1-associated transcriptional axis. (A and B) UMAP of 6,589 control cells: Leiden clusters with the hepatocyte-like cluster C0 highlighted and log-normalized *PROM1* (A). *PROM1* in C0 versus remaining control cells (B): cells with ≥ 1 UMI (bars, all cells; points, pools) and log-normalized *PROM1* in *PROM1*-positive cells (line, median). (C) Pseudobulk differential expression, remaining control cells relative to C0; log2 fold change versus mean expression. Colored, FDR < 0.05, |log2 fold change| > 1, concordant in both pools, mean expression ≥ 0 log2 CPM; 15 labels per direction by |moderated t|. (D and E) Acute *PROM1*-response signature along the state axis (40 bins, pool mean, z-scored per gene): ten increased and ten decreased genes by |moderated t|, with *PROM1* and the response score (up − down) above (D); response score and axis on the UMAP (E). All signature genes, Figure S3. (F) Hallmark enrichment, remaining control cells relative to C0; the five camera-confirmed sets with the largest |NES| per direction, q values are shown. Right, mean z-score along the axis per set (thin) and for the union of genes (thick). (G) Regulons with the largest activity change between the lowest and highest decile of the axis (nine per direction, ≥ 25 targets). Lines, axis activity (scale bar, 1 ULM); bars, acute bulk enrichment (GoF, LoF); dark, FDR < 0.05 in rotation test and camera; light, n.s. Control cells only; n = 2 pools, each comprising two independently transduced cultures. Acute GoF/LoF data in (G) are from Figure 1. Statistics: limma-voom (pseudobulks in C and F; acute bulk in G); GSEA and camera (F); CollecTRI ULM with mroast/cameraPR for bulk bars (G); Benjamini–Hochberg correction. See STAR Methods, Table S2 and Figures S2 and S3.

To characterise this heterogeneity without predefined groups or response genes, we constructed a diffusion coordinate from control cells. No root cell, anchor genes or response score defined it; its direction was subsequently oriented towards increasing endogenous PROM1 (Figure 2E). This PROM1-associated axis was the leading diffusion component and matched the leading principal component of control cells (ρ = 0.93; Table S2). Cell-cycle variation occupied separate components, making cell-cycle regression optional: omitting it left the axis essentially unchanged (ρ = 0.95; Table S2), and cycling fractions were similar in C0 and remaining cells (Figure S2E). The axis was robust to technical covariates, signature-gene exclusion and alternative construction parameters (ρ ≥ 0.978; Table S2).

The axis recapitulated the endogenous PROM1 distribution, and the response score increased along it in both pools (ρ = 0.70 and 0.69), with opposing component-score trends (Figure 2D). This concordance was not limited to the aggregate score: most individual genes were concentrated at the poles of the axis rather than changing gradually (Figure S3A), and gene position along the axis correlated with acute activation effects across measurable activation-significant genes (ρ = 0.63) and activation-specific genes (ρ = 0.67), but not across the broader knockdown-significant set (ρ = −0.03; Figure S3B1–B4).

Pseudobulk analysis identified 219 genes higher in C0 and 534 elsewhere (FDR < 0.05, |log2 fold change| > 1). C0 expressed hepatic secretory/metabolic genes, including APOB, APOA2, FGA, FGG, ASGR2 and PAH, whereas the remaining cells were enriched for PROM1, CD44, DKK1, TM4SF1, TIMP1 and CCN2 (Figure 2C; Table S2). Of 50 Hallmark sets, 24 were significant by GSEA and 16 additionally supported by camera. Along the axis, hepatic/metabolic profiles declined, including fatty acid/bile acid metabolism and oxidative phosphorylation, while EMT, hypoxia and inflammatory/stress-associated profiles increased (Figure 2F; Table S2). Inferred HNF1A, HNF4A, ONECUT1, CEBPA and nuclear-receptor activities decreased, whereas JUN, ETV4 and SMAD4 increased; all 18 displayed regulons were concordant across pools (Figure 2G; Tables S1 and S2). HNF1A, HNF4A and ONECUT1 thus connected the axis to the opposing acute perturbation effects. HNF4A and ONECUT1 were also reciprocally regulated at the transcript level, whereas HNF1A changes reflected target expression only.

### Sustained PROM1 activation depletes the hepatocyte-like compartment and shifts cells along the control axis

We next examined 5,898 PROM1-CRISPRa cells. Detectable PROM1 increased from 51.1% to 93.3%; CD133 induction was confirmed by immunoblotting (Figures 3A and S2A). The hepatocyte-like C0 compartment decreased from 35.0% to 9.8%, consistently across pools (33.8%/35.9% versus 10.9%/8.8%; Figures 3B and S2B). Depletion persisted across preprocessing and clustering choices, without evidence of inferior C0 data quality, and four estimators independent of joint clustering recovered 2.6- to 3.7-fold depletion, including a signature excluding all bulk activation-responsive genes (Figures S2C, S2D and S2F). Cycling fractions were similar between conditions (Figure S2G).

**Figure 3.**
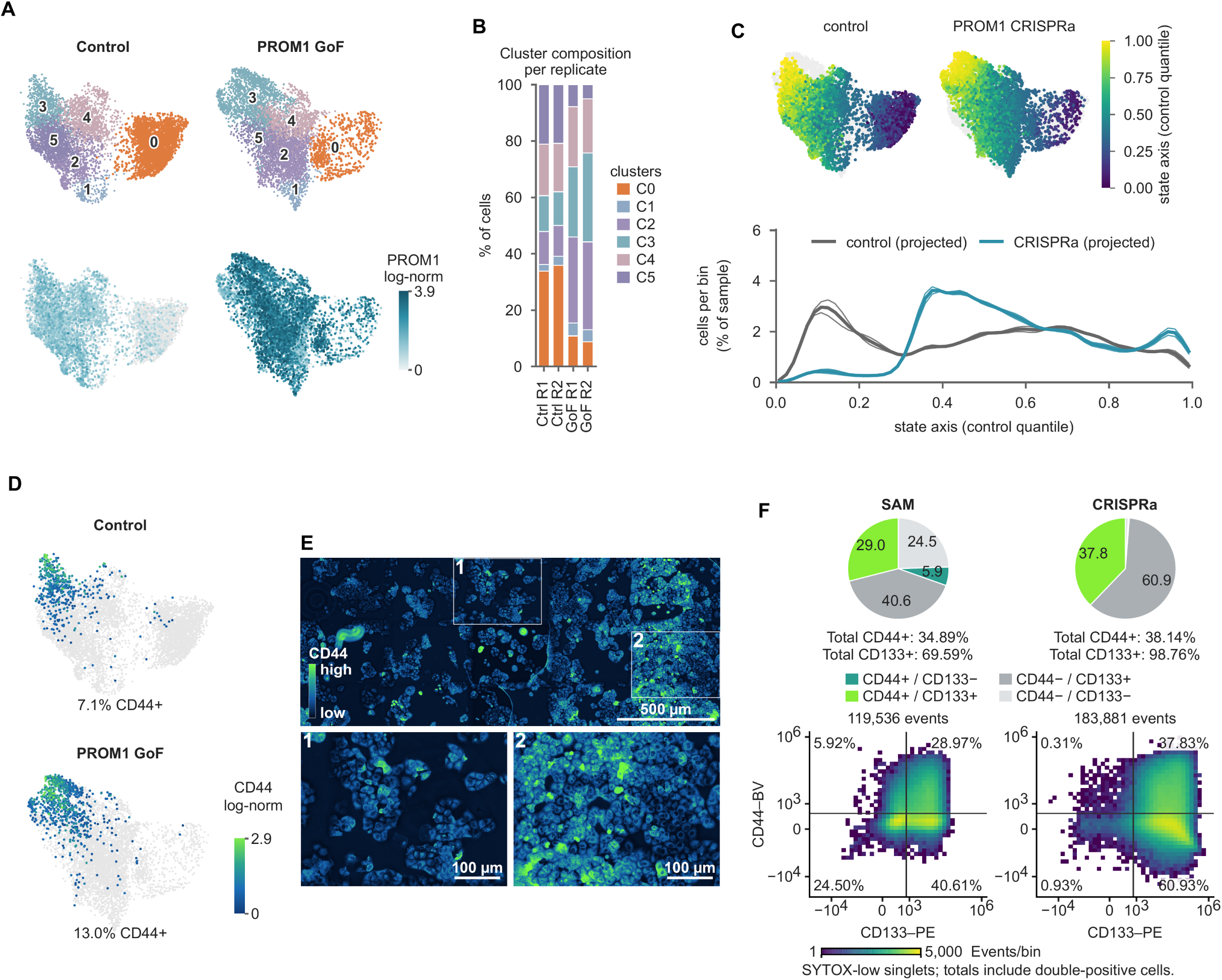
Sustained PROM1 activation depletes the hepatocyte-like compartment and shifts cells along the control axis. (A and B) UMAP of control (6,589) and PROM1-CRISPRa (5,898) cells: Leiden clusters with C0 highlighted and log-normalized *PROM1* (A); cluster composition per pool (B). (C) Top, projection onto the frozen control-derived axis. Bottom, cell distributions (60 bins, smoothed), with controls projected identically; thick lines, pool means; thin lines, pools. (D and E) Log-normalized CD44 per condition, percentages give cells with ≥ 1 UMI (D); CD44 immunofluorescence after activation with boxes 1 and 2 enlarged below (E). Scale bars, 500 µm (overview), 100 µm (enlargements). (F) Flow cytometry of CD133 and CD44 48 h after transient transfection, a separate experiment without bystander cells. Pie charts and density plots, quadrant percentages of SYTOX-low singlets; totals include double-positive cells. n = 2 pools per condition, each from two independently transduced cultures (A–D); one experiment (E, F). Descriptive; no inferential test of condition. See STAR Methods; values in Table S2 and Data S1. See also Figures S2, S4 and S5.

Projection onto the frozen control axis showed depletion of the early C0-associated peak and enrichment of adjoining intermediate positions (Figure 3C). A minority of activated cells (14.2%, versus 5% of control cells by construction) exceeded the control neighbourhood-distance threshold; these accounted for half of the cells at the distal PROM1-high end, suggesting states beyond the control range there, and limited how fully the projection represented them.

CD44 specifically marked the distal PROM1-high end of the axis and was rarely detected elsewhere (Figure S3A). CD44 detection increased from 7.1% to 13.0%, and immunofluorescence correspondingly showed isolated CD44-high cells against a CD44-negative background (Figures 3D and 3E; Table S2). In one separate experiment 48 h after transient transfection, without bystanders, flow cytometry showed increased CD133 (69.6% to 98.8%), CD44 (34.9% to 38.1%) and double positivity (29.0% to 37.8%; Figure 3F; Data S1). Together, sustained activation reduced the PROM1-low, hepatocyte-like state and enriched intermediate positions along the pre-existing axis.

### PROM1 activation has context-dependent effects in ancillary functional assays

At a later stage of passage PROM1 induction was reconfirmed (Figure S4A). Activated cells showed reduced colony and sphere readouts, while their conditioned medium elicited less monocyte migration and endothelial network formation (Figure S4B–G); mean HUVEC mesh counts were 1.5 versus 79 at 6 h and 2.5 versus 79 at 24 h. These assays were descriptive, comprising one experiment per assay except two sphere experiments with different formulations. Conversely, activated cells produced larger mean CAM graft volumes in both repeats (Figure S5A; Data S1): 12.62 versus 3.14 mm^3^ and 12.04 versus 5.45 mm^3^ in the two repeats. A repeat-adjusted exploratory model of all 46 grafts supported larger untreated CRISPRa grafts (geometric-mean ratio 2.57; 95% CI 1.41–4.67; Holm-adjusted p = 0.0114) and reduced volume under trametinib within CRISPRa (ratio 0.248; p = 0.00231), but did not establish differential treatment sensitivity (interaction p = 0.151; all p values Holm-adjusted). Qualitative histology showed no obvious Ki-67 or cleaved-caspase-3 difference and modestly more CD44-positive cells in the illustrated activated graft (Figure S5B).

### HuH-7 cells and a human HCC case show a related PROM1-associated axis

We asked whether this organisation extends beyond PLC/PRF/5. In a public HuH-7 dataset (GSE103867;^16^ 2,075 cells), the cells formed two disconnected subpopulations, so we constructed a separate axis for each (Figures 4A and 4B). Rather than separating PROM1-low and PROM1-high compartments, PROM1 increased gradually along both axes (ρ = 0.42 and 0.23; detection 11% to 69% and 44% to 77% from first to last decile); cell-cycle regression was again optional (ρ = 0.98 and 0.96 without it), and ordering was stable in G1 cells and without signature and C0-marker genes (ρ ≥ 0.93; Table S3). The acute response score followed the axis, with opposing trends of its up- and down-components (ρ = 0.49/0.41 and −0.36/−0.51; Figure 4B; Table S3).

**Figure 4.**
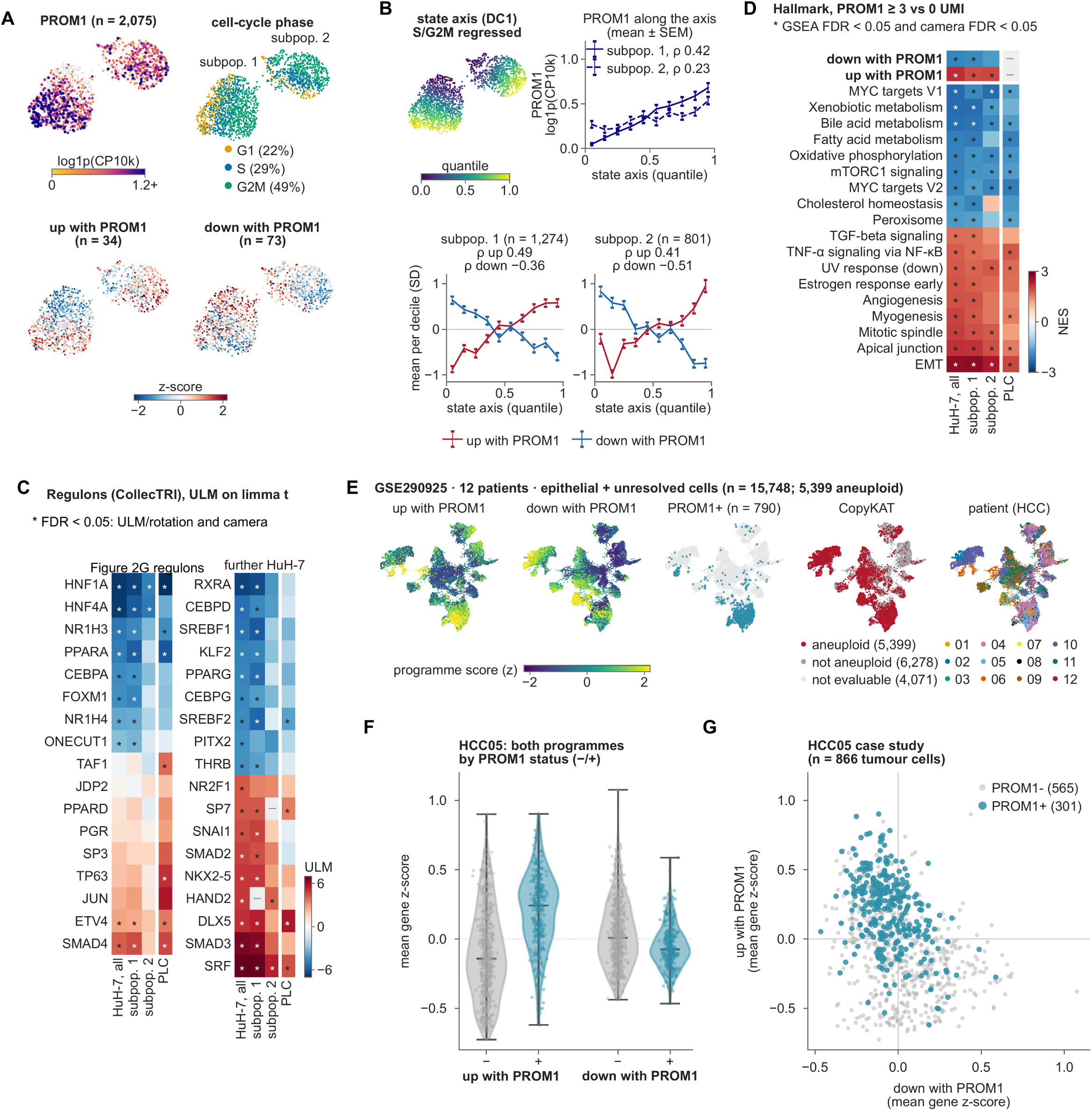
HuH-7 cells and a human HCC case show a related PROM1-associated axis. (A and B) HuH-7 (GSE103867; 2,075 cells): PROM1, cell cycle, subpopulations and response signatures (31/34 upregulated and 68/73 downregulated genes measurable; A). Diffusion axes after cell-cycle regression (B), with PROM1 (mean ± SEM) and signature scores (mean ± SD) per decile; Spearman correlations indicated. (C and D) Inferred regulon activity (CollecTRI, ULM on the limma t) in HuH-7 and PLC/PRF/5: the regulons of Figure 2G and further ones significant in HuH-7 (C). Hallmark enrichment in HuH-7 cells with ≥ 3 versus 0 *PROM1* UMI, beside PLC/PRF/5, signature sets on top (D). Asterisks: FDR < 0.05 in camera and in rotation-test/ULM (C) or GSEA (D). (E) 12 public HCC biopsies (GSE290925), epithelial and unresolved cells (n = 15,748): both score components, *PROM1* detection, CopyKAT call (aneuploid = putative tumour cells) and patient. (F and G) Both score components in the 866 HCC05 tumour cells by *PROM1* detection, as violins with median and range and points for individual cells (F) and as a scatter (G). One public HuH-7 library and one HCC05 patient; descriptive cell-level analyses. Statistics: limma-trend contrasts (C, D); ULM, rotation tests and cameraPR (C); GSEA and camera (D); Spearman correlations (B). See STAR Methods, Table S3 and Figure S6.

The two axes shared genes, pathways and regulons: 124 of 145 strongly PLC-axis-associated genes (85.5%) were concordant (Figure S6; Table S3); all 16 robust Hallmark programmes were concordant, with support in HuH-7 for 7 of 8 C0-associated and 5 of 8 PROM1-pole programmes (Figure 4D); and of 17 measurable PLC-axis regulons 16 were concordant, with all eight decreasing ones supported in HuH-7 but only 2 of 9 increasing ones (Figure 4C).

We then turned to 12 public HCC biopsies (GSE290925)^17^, examining the epithelial compartment (15,748 cells), of which CopyKAT classified 5,399 as aneuploid; we treated these as putative tumour cells (Figure 4E; Table S3). HCC05 was the only case with sufficient PROM1-positive tumour cells for a within-case comparison (301 of 866; Methods), and there PROM1-detected cells favoured the activation-upregulated score component and opposed the downregulated component (Figures 4F and 4G).

Comparing all three datasets on the 145 genes most strongly bound to the PLC/PRF/5 axis, PLC/PRF/5 and HuH-7 agreed in both directions along their independently constructed axes. HCC05 tumour cells were ordered using the 85 response-signature genes absent from the heatmap, separating a PROM1-low from a PROM1-high compartment (PROM1 detection 7.9% to 54.8% across score quartiles). Of the 145 displayed genes, 118 (81.4%) were concordant with PLC (53 of 69 and 65 of 76 at the two ends; Figure S6).

### Endogenous tagging identifies candidate CD133-associated proteins

We inserted the compact ALFA epitope^18^ into endogenous PROM1 at extracellular N-terminal or first cytoplasmic-loop sites, advancing one clone per site (1C4 and 4B4; Figures 5A and 5B). Immunofluorescence supported membrane-associated CD133/ALFA labelling (Figure 5C). After SAM activation, affinity-purification mass spectrometry compared four samples per tagged clone with four untagged, activated controls.

**Figure 5.**
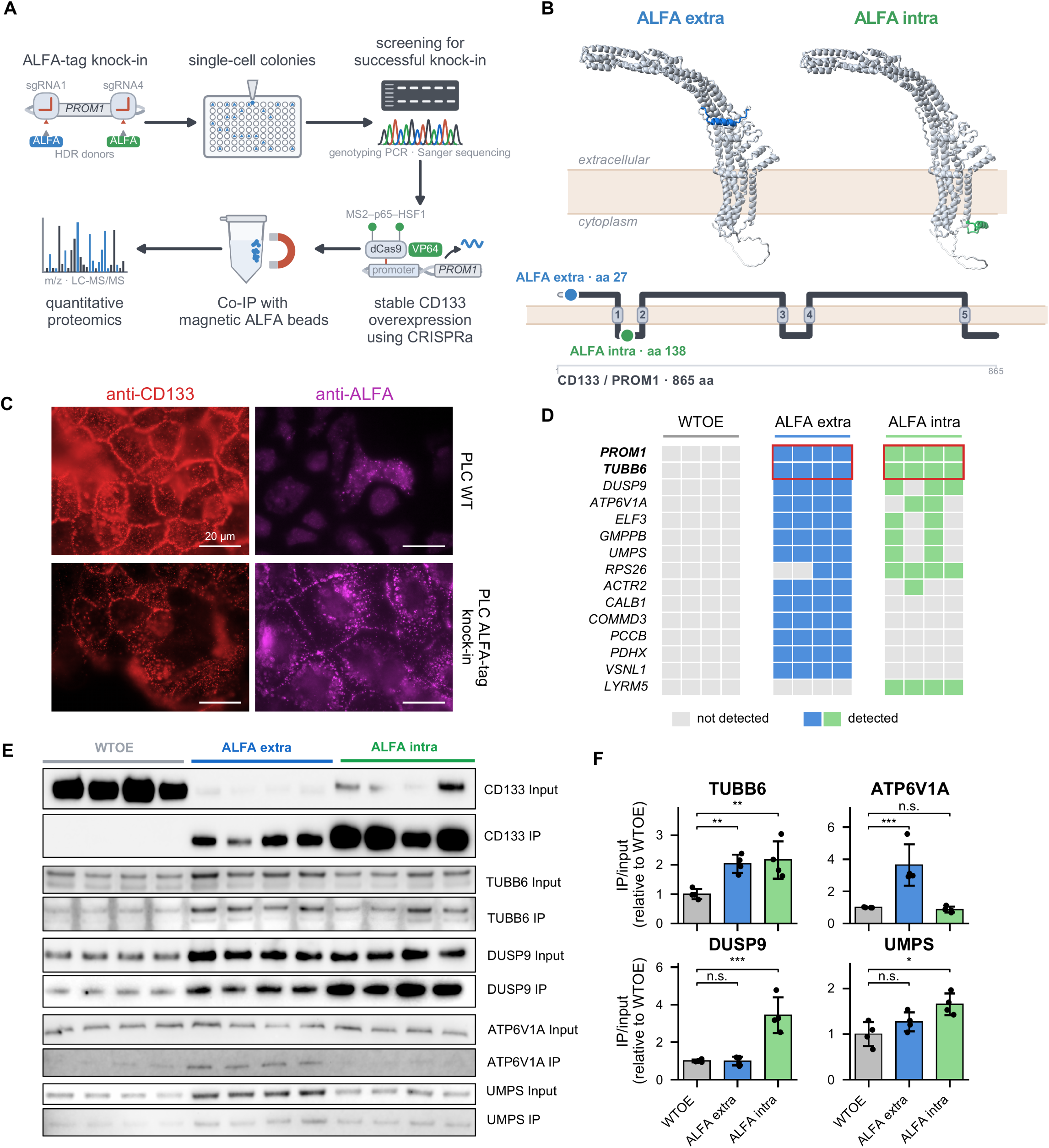
Endogenous tagging identifies candidate CD133-associated proteins. (A–C) Endogenous ALFA tagging and affinity-purification workflow (A); CD133 structure and topology showing insertion at residue 27 (extracellular; clone 1C4) or residue 138 (first cytoplasmic loop; clone 4B4) (B); CD133 (red) and ALFA (magenta) immunofluorescence in parental and tagged cells (C). Scale bars, 20 µm. (D) Detection screen: protein groups absent from every control and present in all four replicates of at least one tagged clone; one column per sample. PROM1 is the bait. (E and F) Input and immunoprecipitate immunoblots (four lanes/group; E) and IP/input densitometry relative to the untagged control mean (F). Bars, mean ± SD; points, replicates. CD133 is omitted because controls yield no pulldown. n = 4 samples per group, with one edited clone per tagging site, one experiment (D–F). Statistics (F): unpaired two-tailed Student’s t test on log2 values, Holm-Šidák across the two comparisons per protein; *p < 0.05, **p < 0.01, ***p < 0.001; n.s., not significant. See STAR Methods; all protein groups in Table S4.

Capture of the bait was highly specific: CD133 was identified by MS/MS in all eight tagged samples and in none of the controls, although these carried comparable CD133 after activation. Despite preclearing, however, the beads retained some background of unrelated proteins that was equally present in the controls.

A detection-based screen retained 15 of 1,519 protein groups absent from all controls and present in all four samples of at least one tagged clone (Figure 5D; Table S4). TUBB6 was directly identified by MS/MS in all eight tagged samples; positive ATP6V1A, DUSP9 and ELF3 entries also had direct MS/MS support, whereas two cytoplasmic-tag UMPS entries relied on matching between runs. Complementary LFQ analysis identified 19 groups enriched in both clones, including EHD4, SCYL2, ARFGAP2/3 and SMAP2. The nonoverlapping screens together prioritised 33 non-bait candidates.

Immunoblotting showed strong tagged CD133 recovery and supported TUBB6 co-enrichment in both configurations. ATP6V1A, DUSP9 and UMPS had clone-dependent recovery patterns (Figures 5E and 5F; Data S2). Of note, co-purification here identifies candidate associations rather than establishing direct interaction.

## Discussion

Our data suggest that PROM1 contributes to hepatocytic identity programmes and the distribution of transcriptional cell states in PLC/PRF/5 cells. Acute perturbation altered programmes that already characterised the heterogeneity of untreated cells, while sustained activation was associated with a reduced proportion of cells displaying a hepatocyte-like phenotype. Our study thus extends the view of CD133 as a marker to include its potential functional involvement in this state variation. Based on these findings, we hypothesise that CD133 can actively influence intratumoral heterogeneity in HCC along a dedifferentiation axis by regulating hepatocytic identity programmes.

This interpretation builds on the stem- or progenitor-like properties described in CD133-positive HCC cells, including enhanced self-renewal, colony formation, tumour initiation and chemoresistance.^4^ Direct perturbation studies also support a functional contribution of CD133: reducing its expression impaired colony formation, cell-cycle progression and tumorigenicity, while other studies identified roles in autophagy under glucose deprivation and in tumour growth.^19–21^ In models of MASLD-associated HCC, CD133 promoted the stabilisation of active β-catenin through an interaction with MYH9, thereby supporting stem-like properties and tumorigenesis.^9^ Our findings extend these functional associations to the regulation of hepatocytic identity programmes.

The study by Zhou et al.^10^ is particularly relevant to the connection with intratumoral heterogeneity. Using lineage tracing and single-cell RNA sequencing, they showed that PROM1-positive cells in murine HCC models form a heterogeneous tumour-cell population that expands clonally in situ and adopts increasingly dedifferentiated states with pronounced proliferative and stem-cell features during tumour progression. Our study builds on this relationship by examining whether changing PROM1 expression itself influences the hepatocytic identity of HCC cells.

Man et al.^22^ provide a further functional link. They identified SPINK1 as a factor enriched in CD133-positive HCC cells that promotes dedifferentiation, self-renewal and chemoresistance through EGFR–ERK–CDK4/6–E2F2 signalling. Their work thus establishes a functional pathway connecting properties of the CD133-positive compartment to altered differentiation. The ablation of PROM1-positive cells examined in that study removes the labelled compartment, whereas our experiments specifically alter PROM1 expression within the cell population. These approaches therefore complement one another in addressing how CD133-associated heterogeneity arises and is regulated.

CRISPRa-mediated activation of the endogenous PROM1 locus allows this relationship to be investigated within the context of the existing gene structure and cellular RNA processing machinery.^14,23^ This is relevant given the alternative promoters and splice variants of PROM1, because, unlike overexpression of a defined cDNA, this approach does not prescribe a single isoform.^12^ In our data, activation remained restricted to the proximal first exon already used in PLC/PRF/5 cells. The resulting CD133 protein is also available for endogenous post-translational processing. The functional relevance of such processing is illustrated by Liu et al.^24^, who showed that mutation of an N-glycosylation site reduced the interaction with β-catenin and the growth-promoting effect of CD133. Transient and stable PROM1 activation allowed us to relate early transcriptional responses to longer-term changes in the cell population.

The largely reciprocal response to transient PROM1 activation and siRNA-mediated knockdown supports the link between the perturbation and the observed expression changes: 94.4% of the genes significantly altered in both experiments were regulated in opposite directions. The resulting Acute Response Signature formed the basis of the Acute Response Score and enabled us to connect the experimentally observed response to endogenous heterogeneity in untreated cells.

The regulation of hepatocytic identity was central to this response. Following PROM1 activation, hepatocytic identity and functional programmes decreased, whereas EMT-, stress- and inflammation-associated programmes increased. Knockdown predominantly reduced proliferation and cell-cycle programmes and produced changes in the opposite direction in selected liver function-related pathways. HNF4A and HNF1A were among the most strongly altered regulons in both experiments. The reduction in inferred HNF4A regulon activity following PROM1 activation is consistent with the loss of Hnf4a-controlled differentiation regulons in dedifferentiated PROM1-labelled tumour cells in vivo.^10^ Our data recapitulate this relationship in a human system and extend it to targeted perturbation of PROM1 expression itself.

HNF4A is an established central regulator of hepatocytic identity. Its hepatocyte-specific deletion prevents the formation of a hepatic epithelium and disrupts liver morphogenesis.^25^ Together with HNF1A and HNF6, it forms a densely interconnected hepatic regulatory network,^26^ and, together with HNF1A, it represses mesenchymal master regulators whose sustained repression is required to maintain the hepatocytic state.^27^ Particularly informative for the direction of our findings is the previously described reverse relationship: enforced HNF4A expression in hepatoma cells induces differentiation towards hepatocytes and markedly reduces stemness gene expression and the proportion of CD133-positive cells.^28^ Similarly, enforced HNF4A expression suppresses stem-cell markers and β-catenin activation and prevents HCC development.^29^ Thus, whereas restoration of hepatocytic identity reduces the CD133-positive fraction in those studies, changing PROM1 expression alters hepatocytic identity programmes in our experiments. These observations support a functional relationship between CD133 and hepatocytic identity from different directions of intervention.

The single-cell analyses show that the acute PROM1 response aligns with pre-existing cell-state variation. In untreated PLC cells, the Acute Response Score corresponded to the distribution of PROM1-low and PROM1-high subpopulations. The diffusion-based state axis derived after regression of cell-cycle effects also correlated strongly with the score and reflected endogenous PROM1 expression. Because this axis was constructed independently of the perturbation signature, their agreement connects two distinct approaches to population heterogeneity: the experimentally induced response and the pre-existing transcriptional organisation. PROM1 therefore affects programmes that contribute to endogenous state variation and may itself influence their expression.

The coexistence of PROM1-high and PROM1-low cells could reflect a dynamic distribution among interconverting states. Experimental precedents exist for such plasticity: CD133-positive and CD133-negative fractions sorted from HuH7 and HuCCT1 cells regenerated mixed populations during subculture.^30^ Rapid recovery of CD133 surface expression after sorting negative or low-expressing cells has also been reported in colorectal cancer cell lines.^31^ These studies primarily demonstrate the plasticity of the marker phenotype. Further evidence for dynamic state distributions comes from breast cancer models in which sorted fractions re-established characteristic state proportions or non-stem-like cells transitioned into stem-like states.^32^ A bivalent chromatin configuration at the ZEB1 promoter was identified as a basis for signal-dependent plasticity.^33^ Against this background, our data suggest a model in which PROM1 influences the distribution along a pre-existing state axis and, if a dynamic equilibrium exists, modulates its balance.

The results of sustained PROM1 activation are consistent with this model. The proportion of the hepatocyte-like subpopulation, which expressed low levels of PROM1 in controls, was markedly reduced. Accordingly, the hepatocytic end of the control-derived state axis was less densely occupied, whereas cells accumulated at intermediate positions. Acute perturbation and sustained activation thus capture two compatible levels of observation: early changes in state-associated expression programmes and a longer-term shift in population composition.

Experimental and persistent shifts in such distributions have also been described in liver cancer cell lines. Martin et al.^34^ observed an increased proportion of CD133-positive cells following TGF-β treatment that persisted for several cell divisions after stimulus withdrawal and was associated with sustained changes in DNA methylation. Whereas that study described a shift induced by an external signal, our experiments directly target expression from the endogenous PROM1 locus. The early response after two days and in the absence of antibiotic selection strengthens the interpretation of a transcriptional effect preceding the longer-term change in population composition. Together, these findings support the hypothesis that PROM1 contributes to the expression and distribution of the observed cell states.

The complementary functional assays show that this altered state distribution is accompanied by context-dependent phenotypes. The stably PROM1-activated population produced larger mean graft volumes in both CAM experiments, whereas colony and sphere formation were reduced. These assays impose different requirements for adhesion and survival, whereas CAM grafts interact with matrix and vascularised tissue. Enrichment of intermediate states and changes in adhesion programmes could therefore have different consequences across environments.

Reanalysis of the HuH7 data from Zheng et al.^16^ extends the transcriptional relationship between PROM1 and hepatocytic identity beyond PLC/PRF/5. In the original study, stem-cell and HCC markers, including PROM1, displayed heterogeneous expression without defining discrete marker-based clusters. Our analysis shows that this heterogeneity can nevertheless be described as an ordered variation in cell state. The HuH7 axis, constructed analogously to the PLC axis and independently of the Acute Response Signature, followed endogenous PROM1 expression. Unlike in PLC, this relationship was predominantly gradual. Associated pathways and regulons also showed substantial agreement, particularly in the attenuation of hepatocytic programmes with increasing PROM1 expression. Both components of the Acute Response Signature followed this trend in the expected directions. PROM1-associated organisation is therefore also discernible where it does not manifest as a separation into discrete subpopulations.

Analysis of twelve treatment-naïve HCC biopsies from the dataset of Liu et al.^17^ provides a further connection to intratumoral heterogeneity. Among the putative tumour cells classified as aneuploid, HCC05 contained enough PROM1-detected aneuploid cells for a detailed within-tumour comparison. Within this tumour, we ordered cells according to the Acute Response Score derived from the perturbation experiments. PROM1-rich cells accumulated at the pole associated with genes upregulated with PROM1, whereas cells without detectable PROM1 occupied the opposite end. Although PROM1 itself was not used to define the ordering, its expression therefore followed the signature derived in cell culture. This signature can thus reveal a PROM1-associated component of heterogeneity in human HCC. Unlike the cell-line axes constructed independently of the signature, HCC05 represents an application of the signature to a patient tumour.

The similarities between datasets extend beyond the Acute Response Signature. Most genes strongly associated with the PLC axis were directionally concordant in each external dataset, including along the score-based ordering in HCC05. Attenuation of the hepatocytic secretion and synthesis programme was particularly consistent, with corresponding genes decreasing in all three datasets. A substantial component of the state variation described in PLC is therefore also evident in a second cell line and a patient tumour.

Along the PLC/PRF/5 axis, individual genes show distinct patterns that further characterise its terminal region. ANXA3 increased across much of the axis, with a more pronounced rise near its end, whereas PAH decreased and reached particularly low expression at the endpoint. CD44 expression was strongly concentrated towards the PROM1-high end. GPRC5A displayed a similar pattern, although this was less clear in HuH7 and HCC05. The terminal region thus combines accentuated gradual changes with preferential expression of individual markers. Immunofluorescence also supports pronounced CD44 heterogeneity at the protein level: individual strongly positive cells were observed against a largely negative background. This finding suggests that the uneven distribution of CD44 is not solely attributable to missing transcript detection in single-cell RNA sequencing.

This organisation provides a possible transcriptional context for the CD133/CD44 marker combination described in HCC. Zhu et al.^35^ showed that the high tumour-initiating capacity of CD133-positive cells was primarily enriched in the subpopulation that also expressed CD44. Hou et al.^36^ further associated a CD133^+^/CD44-high phenotype with increased metastatic potential. Together with the concentration of CD44 expression at the end of our PROM1-associated continuum, these findings suggest that CD133/CD44-positive cancer stem cells may be preferentially enriched in this terminal region. The broader continuum would thus reflect a gradual attenuation of hepatocytic identity, while CD44 marks a more restricted state potentially associated with CSC properties.

Hu et al.^37^ provide a complementary conceptual framework. Based on transcriptional trajectories and reconstructed clonal relationships, they propose a model in which CSCs emerge as late products of dedifferentiation in established human HCC. This supports the idea that stem-like properties can be acquired during the progressive attenuation of differentiated cellular programmes. Our observation that CD44 expression is concentrated in the terminal region fits within this model: the CD133/CD44-positive subgroup described in previous functional studies may preferentially occupy the dedifferentiated pole of the broader PROM1-associated state variation. Hu et al. provide the developmental framework for this interpretation, whereas the specific connection to CD44 arises from our expression data and the functional studies by Zhu et al. and Hou et al.

Immunoprecipitation of endogenously ALFA-tagged CD133 adds possible molecular connections between membrane organisation and cell state. TUBB6 was the most consistently co-purified candidate apart from CD133, and its enrichment was confirmed by immunoblotting in both tagging configurations. A functional relationship between CD133 trafficking, signalling and differentiation has previously been described for the CD133–HDAC6–β-catenin complex in other tumour models^38^. The detection of ATP6V1A and the quantitative enrichment of the trafficking-associated proteins EHD4, SCYL2, ARFGAP2, ARFGAP3 and SMAP2 further highlight intracellular membrane transport, while DUSP9 provides another candidate connection to signalling regulation.

ELF3 is particularly relevant to the transcriptional findings. It was detected in both tagging configurations but not in the untagged controls. Man et al.^22^ showed that ELF3 directly activates SPINK1 transcription, linking it to the dedifferentiation-promoting pathway discussed above. Co-purification of ELF3 therefore provides a basis for testing a possible connection between CD133-associated protein complexes and this plasticity programme. Together, these candidates motivate testing whether CD133 influences hepatocytic identity by organising and relaying cellular signals.

### Limitations of the study

Our single-cell data capture state distributions but do not demonstrate that individual cells change state. The axis endpoint denotes a transcriptional position, not a temporal endpoint of tumour evolution. It remains unresolved whether PROM1 promotes transitions, alters hepatocytic programmes within existing states, or shifts relative cell proportions through differences in proliferation or survival. A dynamic equilibrium remains a model; transition rates and recovery after sorting were not measured. Likewise, the mechanistic connection to HNF4A/HNF1A and the functional CSC properties of the CD44-rich terminal region remain to be tested. The IP candidates co-purify with CD133 but establish neither direct binding nor a causal contribution to the transcriptional response. The external evidence extends the transcriptional relationship, while the functional perturbation findings remain restricted to PLC. For HCC05, it should also be considered that the ordering was score-based, aneuploidy does not provide definitive evidence of malignancy, and only one tumour contained sufficient PROM1-positive cells.

## Resource availability

### Lead contact

Further information and requests for resources and reagents should be directed to and will be fulfilled by the lead contact, Alexander Scheiter (m).

### Materials availability

This study generated PROM1-activated PLC/PRF/5 cell populations and ALFA-tagged PROM1 clones. These materials are available from the lead contact upon reasonable request, subject to applicable institutional, legal, and regulatory requirements, including any required approvals for material transfer.

### Data and code availability

Bulk RNA-sequencing data from the PROM1 CRISPRa activation and PROM1 siPOOL-mediated silencing experiments in PLC/PRF/5 cells, together with their respective controls, have been deposited in the NCBI Gene Expression Omnibus (GEO) under accession GSE349777. Single-cell RNA-sequencing data generated using Chromium Flex from PROM1 CRISPRa-activated PLC/PRF/5 cells and spacer-free SAM controls have been deposited in GEO under accession GSE350646. Both bulk and single-cell RNA-sequencing datasets are linked under GEO SuperSeries GSE350647.

Public human HCC data were obtained from GEO accession GSE290925.^17^

Public HuH-7 single-cell data were obtained from GEO accession GSE103867 (HuH-7 library GSM2785127)^16^.

Affinity-purification proteomics results are provided in Table S4. Numerical source data are provided in Data S1, and available full-scan immunoblots in Data S2. Custom EGFP probes and ALFA-tagging oligonucleotides are listed in Tables S5 and S6, respectively.

Any additional information required to reanalyze the data reported in this paper is available from the lead contact upon request.

## Acknowledgments

This work was supported by the intramural UR Potentials Stage 1 – Early Independence grant to AS and by the Deutsche Forschungsgemeinschaft (DFG, German Research Foundation; project number 502688960) to AS. We acknowledge Dr. Stefan Tholen and the Proteomics Facility at the Institute of Clinical Pathology, University Medical Center Freiburg, for LC–MS/MS analysis of the affinity-purification samples and MaxQuant-based data processing and quantitative proteomic analysis. Bulk RNA sequencing, including rRNA depletion and strand-specific library preparation, was performed by Eurofins Genomics (Konstanz, Germany) using the INVIEW Transcriptome service. We acknowledge Monika Kerscher, Ingrid Winkel, and Manfred Meyer for their excellent laboratory support. We thank Sara Steinmann for coordinating the CAM experiments. We also thank colleagues in the Regensburg and San Diego laboratories for their advice throughout the project. We acknowledge the use of Gene Set Enrichment Analysis (GSEA) software and the Molecular Signatures Database (MSigDB; Subramanian et al.^39^; http://www.broad.mit.edu/gsea/).

## Author contributions

Conceptualization, T.S. and A.S.; Methodology, T.S., A.S., K.K., A.I., D.H.; Investigation, T.S., A.S., K.K., M.V., C.R., and E.S.L.; Formal analysis, T.S., A.S.; Validation, T.S., A.S.; Visualization, T.S.; Funding acquisition, A.S., G.-S.F., M.E., and D.F.C.; Supervision, K.U., M.E., G.-S.F., and D.F.C.; Writing – original draft, T.S. and A.S.; Writing – review & editing, all authors.

## Declaration of generative AI use

Generative artificial intelligence tools, including ChatGPT (OpenAI) and Claude (Anthropic), assisted language editing, manuscript refinement, programming, computational workflow development and troubleshooting, data visualization, figure preparation and exploratory interpretation. The authors critically reviewed, modified and validated the resulting code, suggestions and visualizations against the underlying data.

Analyses used primary data and established bioinformatic and statistical software. The authors determined the biological interpretation and conclusions and take full responsibility for the integrity of the analyses, figures and manuscript.

## Supplemental information

Document S1. Supplemental Figures S1–S6 and Tables S5 and S6, with legends. Tables S1–S4 are supplied as separate Excel workbooks. Data S1 contains numerical source data, Data S2 contains full-scan immunoblots.

## STAR Methods

### KEY RESOURCES TABLE

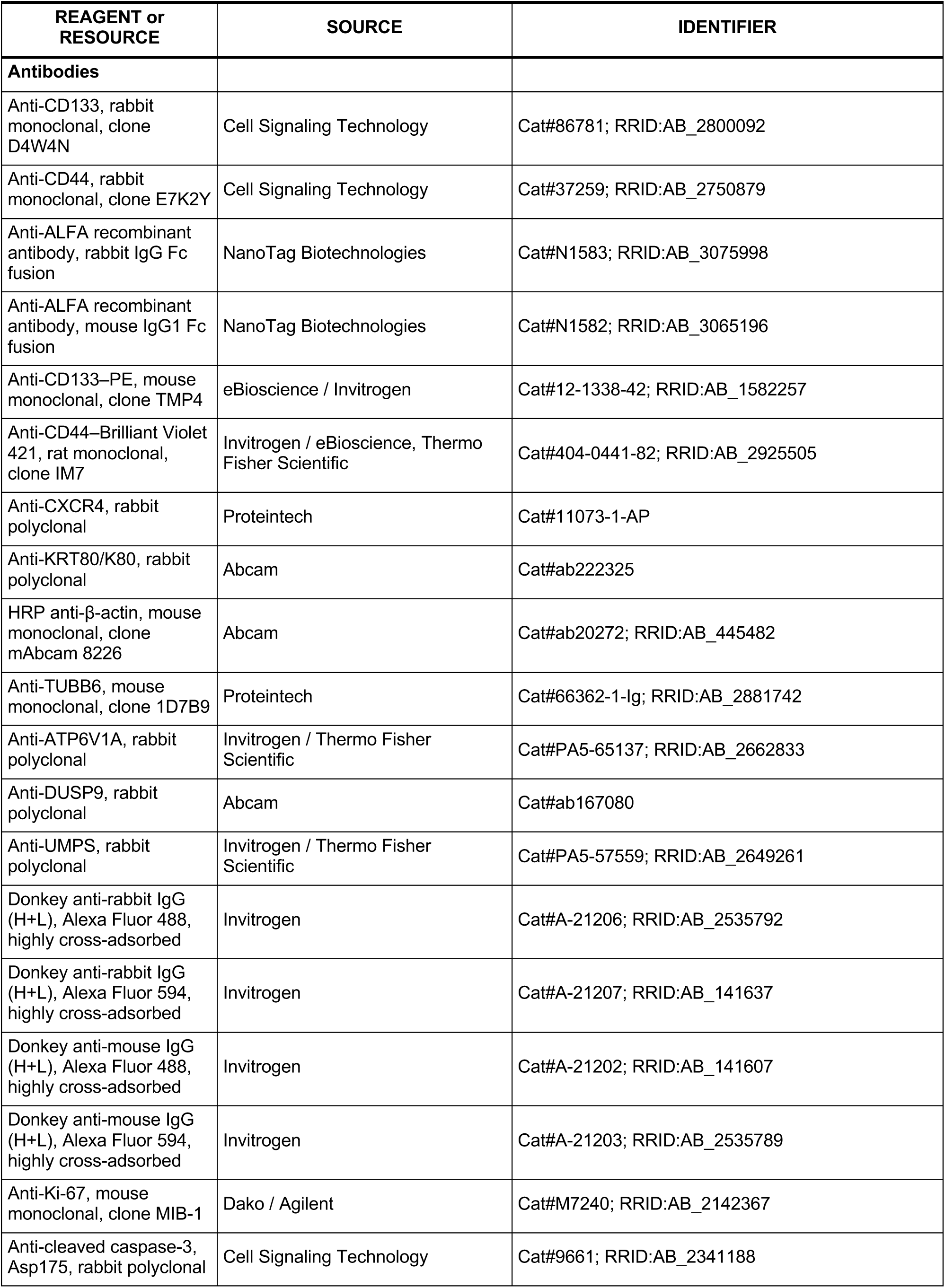
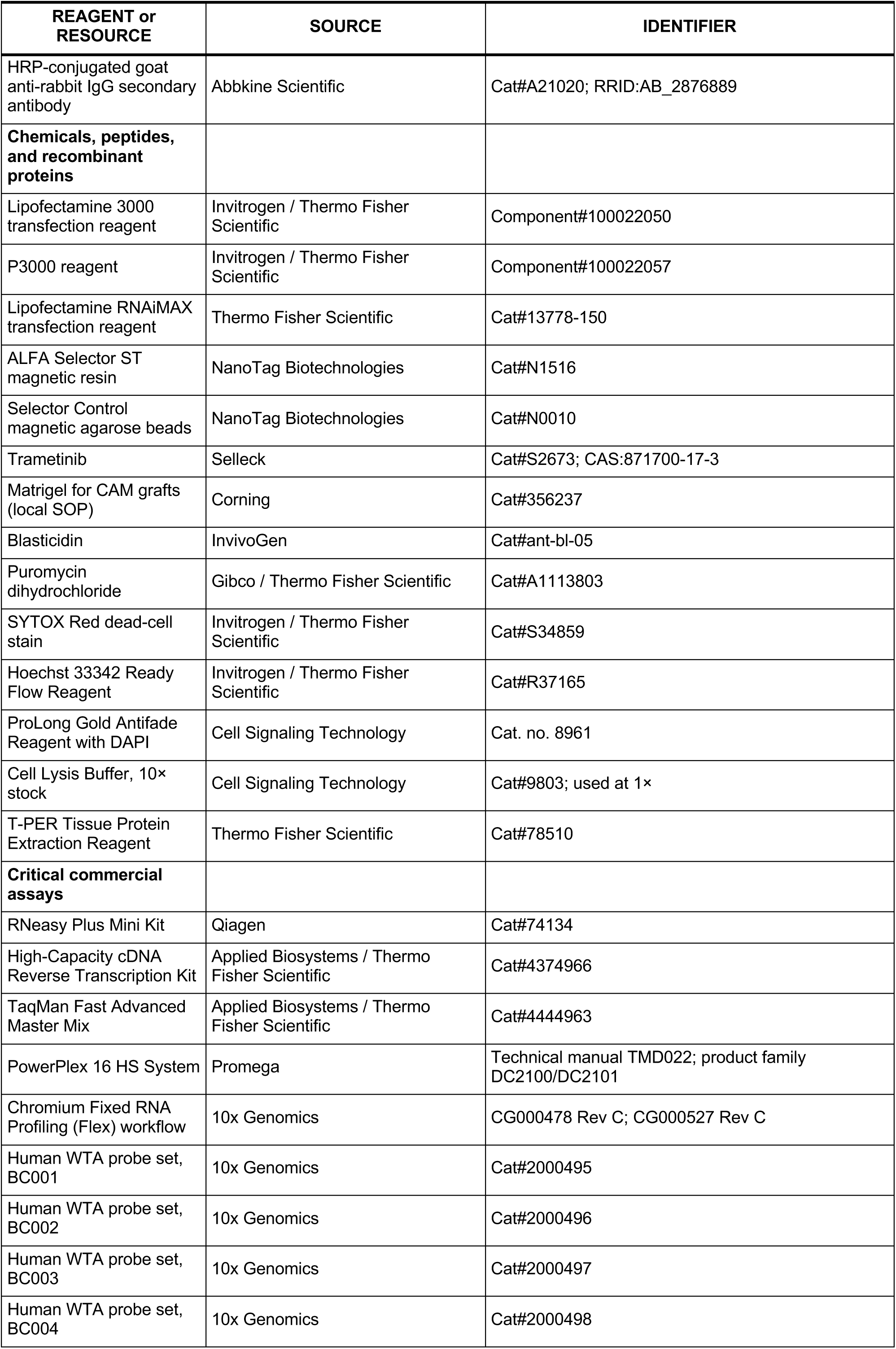
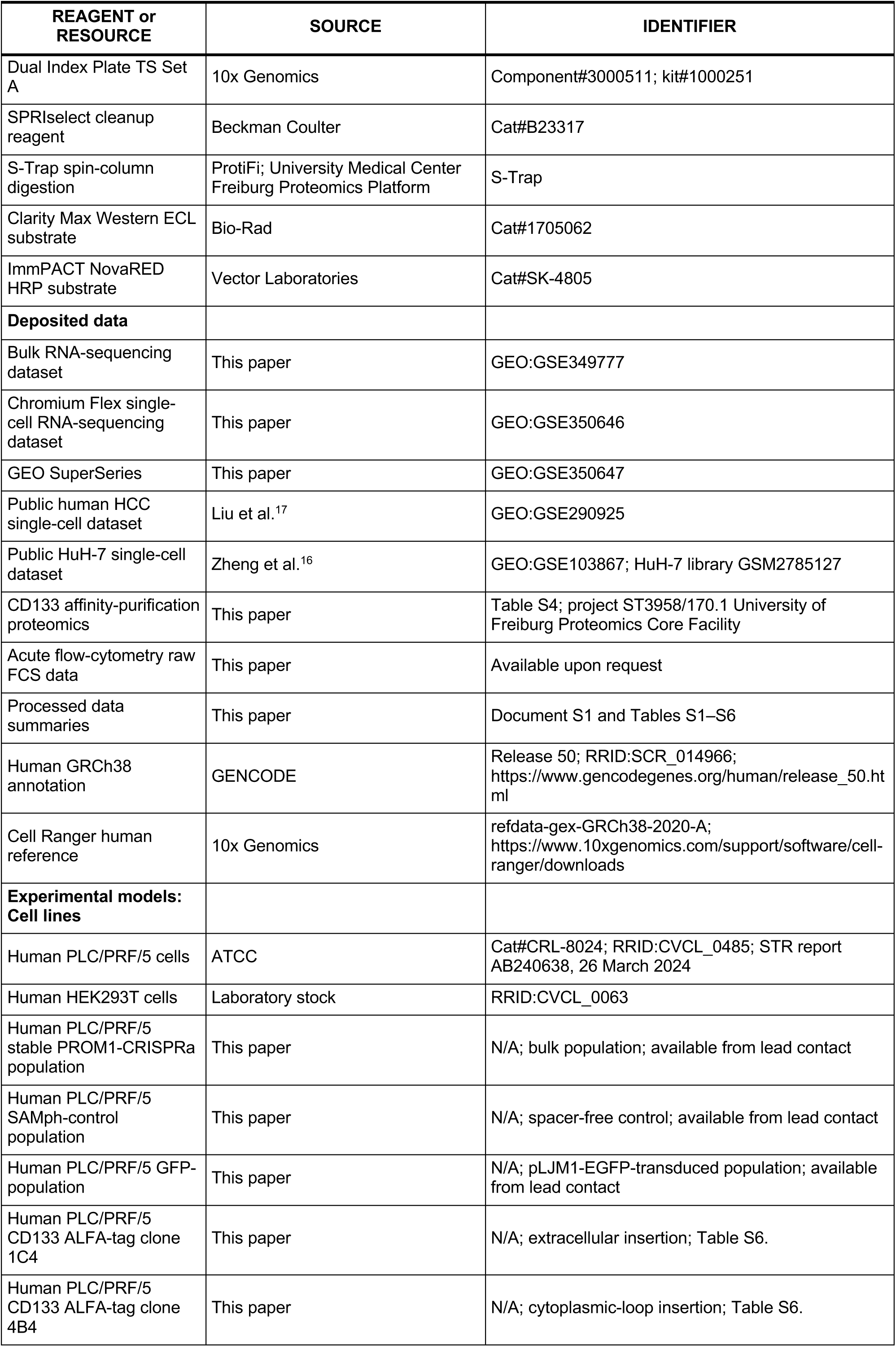
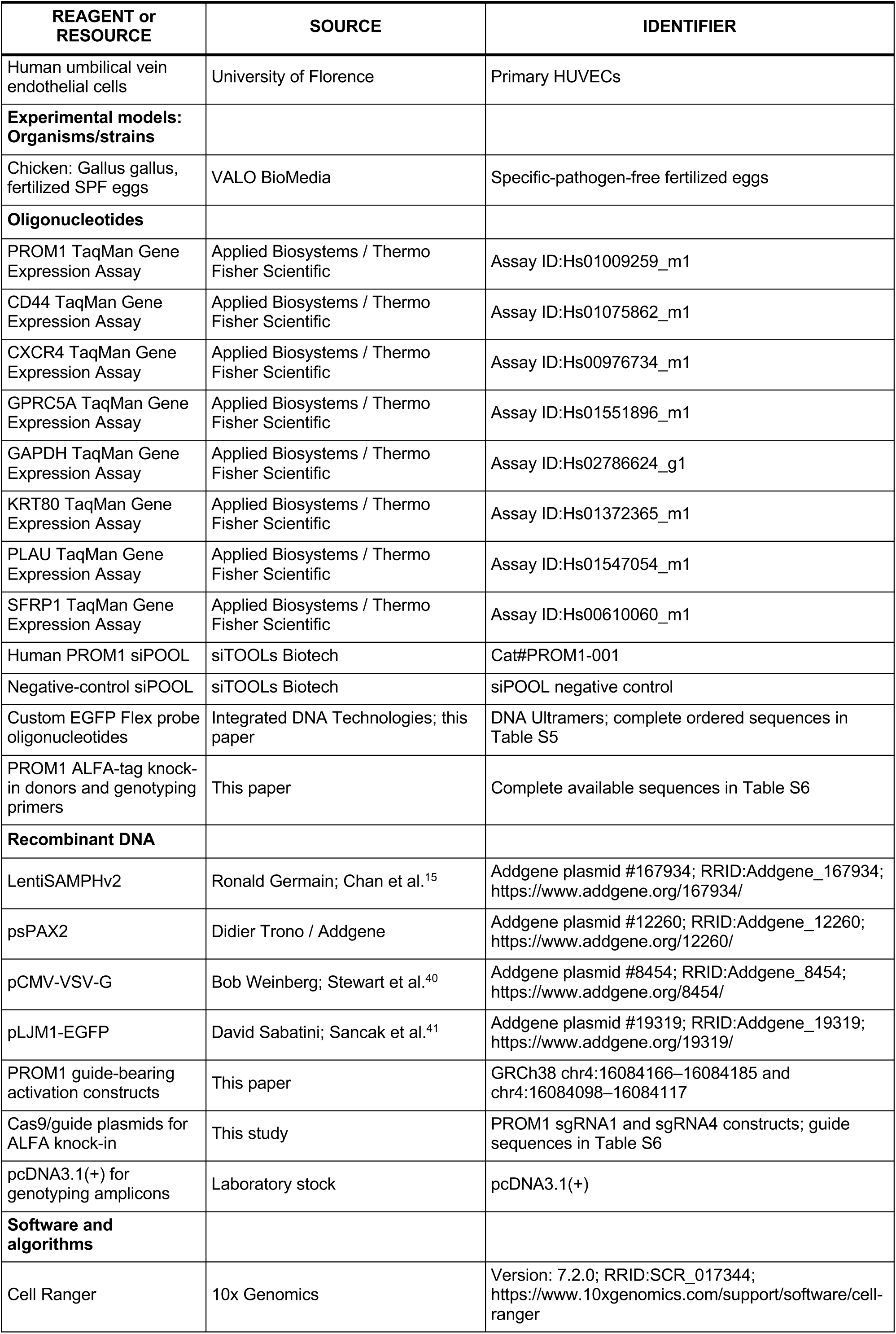
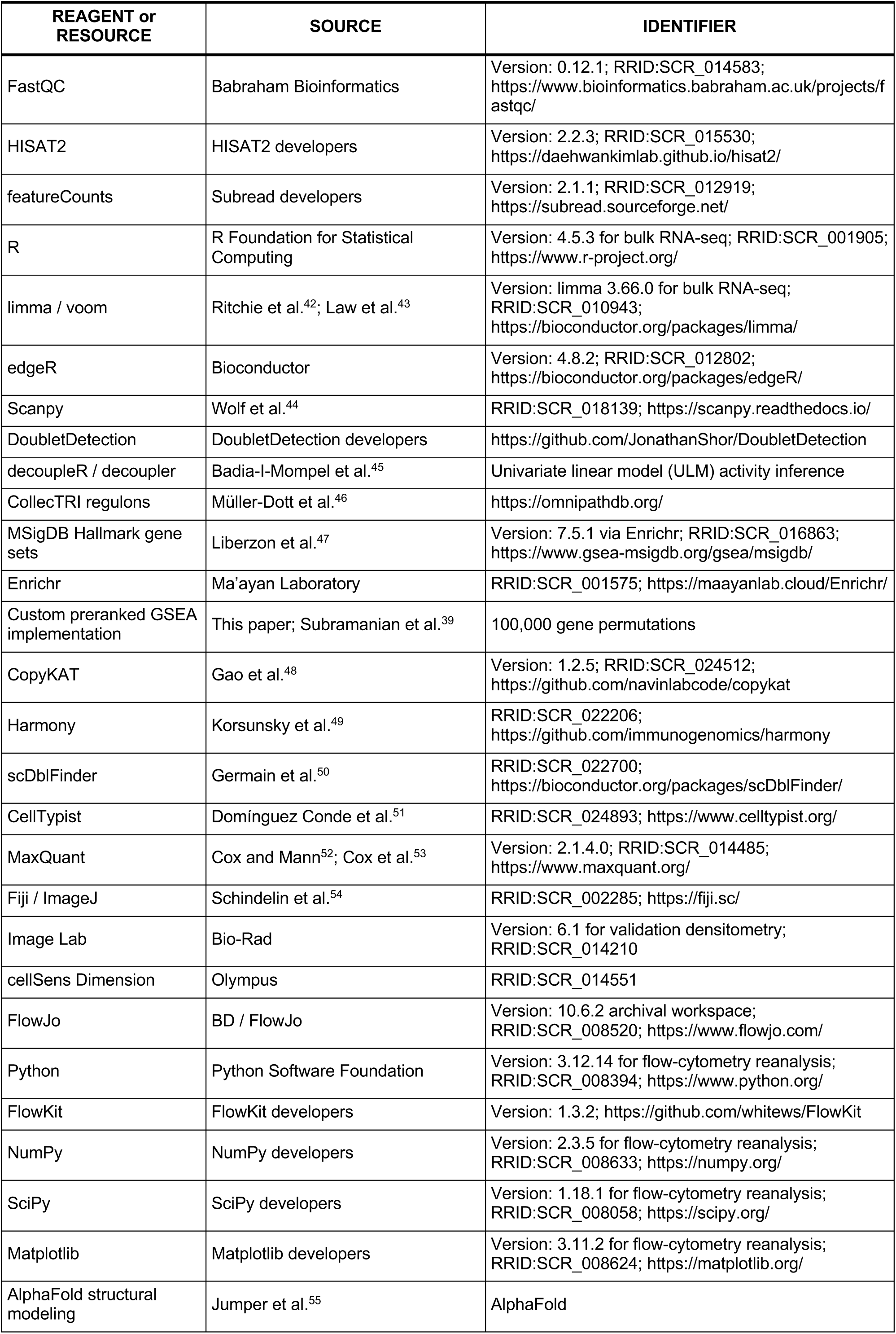
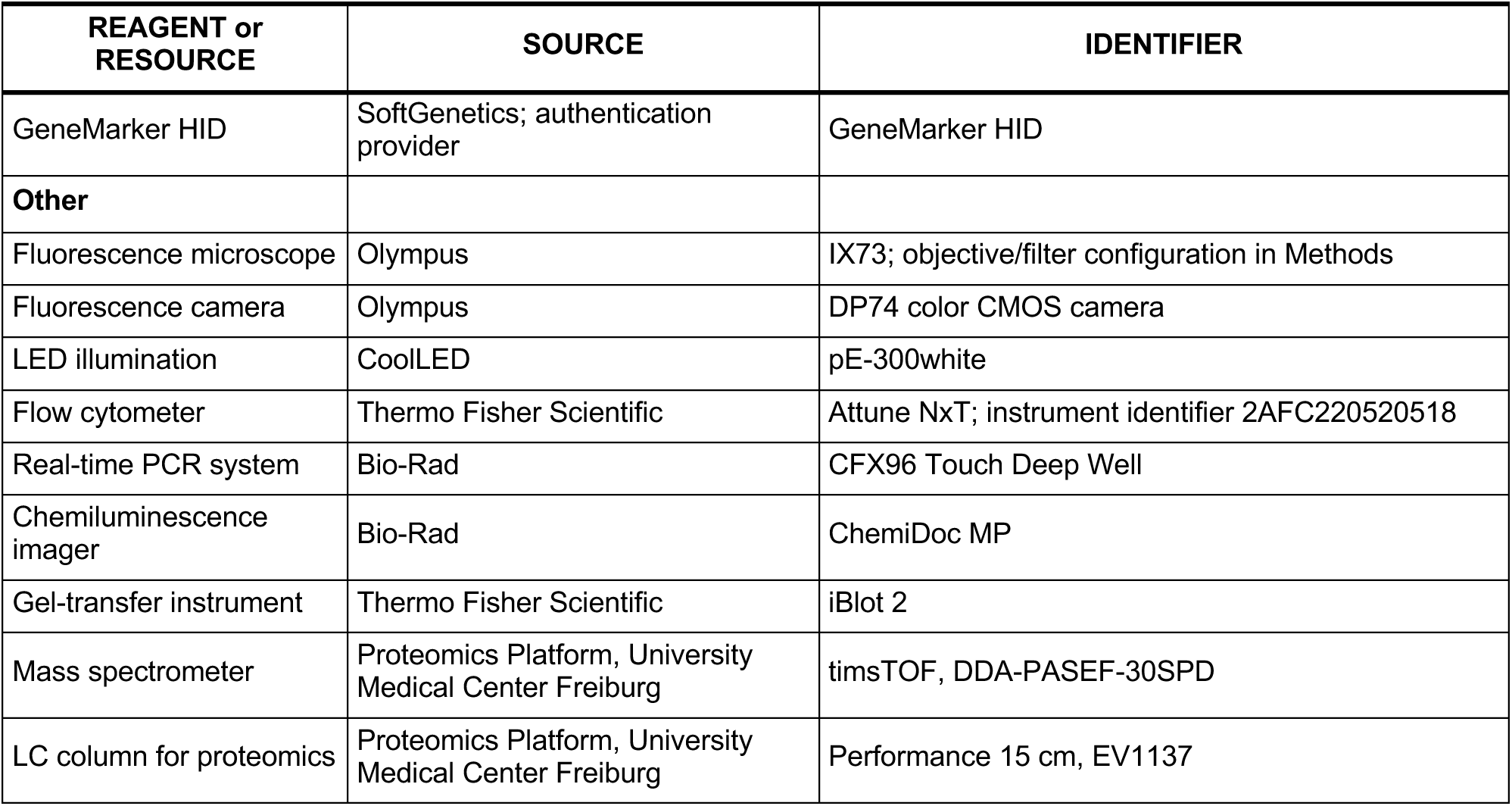

| REAGENT or RESOURCE | SOURCE | IDENTIFIER |
|---|---|---|
| **Antibodies** |  |  |
| Anti-CD133, rabbit monoclonal, clone D4W4N | Cell Signaling Technology | Cat#86781; RRID:AB_2800092 |
| Anti-CD44, rabbit monoclonal, clone E7K2Y | Cell Signaling Technology | Cat#37259; RRID:AB_2750879 |
| Anti-ALFA recombinant antibody, rabbit IgG Fc fusion | NanoTag Biotechnologies | Cat#N1583; RRID:AB_3075998 |
| Anti-ALFA recombinant antibody, mouse IgG1 Fc fusion | NanoTag Biotechnologies | Cat#N1582; RRID:AB_3065196 |
| Anti-CD133–PE, mouse monoclonal, clone TMP4 | eBioscience / Invitrogen | Cat#12-1338-42; RRID:AB_1582257 |
| Anti-CD44–Brilliant Violet 421, rat monoclonal, clone IM7 | Invitrogen / eBioscience, Thermo Fisher Scientific | Cat#404-0441-82; RRID:AB_2925505 |
| Anti-CXCR4, rabbit polyclonal | Proteintech | Cat#11073-1-AP |
| Anti-KRT80/K80, rabbit polyclonal | Abcam | Cat#ab222325 |
| HRP anti-β-actin, mouse monoclonal, clone mAbcam 8226 | Abcam | Cat#ab20272; RRID:AB_445482 |
| Anti-TUBB6, mouse monoclonal, clone 1D7B9 | Proteintech | Cat#66362-1-Ig; RRID:AB_2881742 |
| Anti-ATP6V1A, rabbit polyclonal | Invitrogen / Thermo Fisher Scientific | Cat#PA5-65137; RRID:AB_2662833 |
| Anti-DUSP9, rabbit polyclonal | Abcam | Cat#ab167080 |
| Anti-UMPS, rabbit polyclonal | Invitrogen / Thermo Fisher Scientific | Cat#PA5-57559; RRID:AB_2649261 |
| Donkey anti-rabbit IgG (H+L), Alexa Fluor 488, highly cross-adsorbed | Invitrogen | Cat#A-21206; RRID:AB_2535792 |
| Donkey anti-rabbit IgG (H+L), Alexa Fluor 594, highly cross-adsorbed | Invitrogen | Cat#A-21207; RRID:AB_141637 |
| Donkey anti-mouse IgG (H+L), Alexa Fluor 488, highly cross-adsorbed | Invitrogen | Cat#A-21202; RRID:AB_141607 |
| Donkey anti-mouse IgG (H+L), Alexa Fluor 594, highly cross-adsorbed | Invitrogen | Cat#A-21203; RRID:AB_2535789 |
| Anti-Ki-67, mouse monoclonal, clone MIB-1 | Dako / Agilent | Cat#M7240; RRID:AB_2142367 |
| Anti-cleaved caspase-3, Asp175, rabbit polyclonal | Cell Signaling Technology | Cat#9661; RRID:AB_2341188 |
| REAGENT or RESOURCE | SOURCE | IDENTIFIER |
| HRP-conjugated goat anti-rabbit IgG secondary antibody | Abbkine Scientific | Cat#A21020; RRID:AB_2876889 |
| **Chemicals, peptides, and recombinant proteins** |  |  |
| Lipofectamine 3000 transfection reagent | Invitrogen / Thermo Fisher Scientific | Component#100022050 |
| P3000 reagent | Invitrogen / Thermo Fisher Scientific | Component#100022057 |
| Lipofectamine RNAiMAX transfection reagent | Thermo Fisher Scientific | Cat#13778-150 |
| ALFA Selector ST magnetic resin | NanoTag Biotechnologies | Cat#N1516 |
| Selector Control magnetic agarose beads | NanoTag Biotechnologies | Cat#N0010 |
| Trametinib | Selleck | Cat#S2673; CAS:871700-17-3 |
| Matrigel for CAM grafts (local SOP) | Corning | Cat#356237 |
| Blasticidin | InvivoGen | Cat#ant-bl-05 |
| Puromycin dihydrochloride | Gibco / Thermo Fisher Scientific | Cat#A1113803 |
| SYTOX Red dead-cell stain | Invitrogen / Thermo Fisher Scientific | Cat#S34859 |
| Hoechst 33342 Ready Flow Reagent | Invitrogen / Thermo Fisher Scientific | Cat#R37165 |
| ProLong Gold Antifade Reagent with DAPI | Cell Signaling Technology | Cat. no. 8961 |
| Cell Lysis Buffer, 10× stock | Cell Signaling Technology | Cat#9803; used at 1× |
| T-PER Tissue Protein Extraction Reagent | Thermo Fisher Scientific | Cat#78510 |
| **Critical commercial assays** |  |  |
| RNeasy Plus Mini Kit | Qiagen | Cat#74134 |
| High-Capacity cDNA Reverse Transcription Kit | Applied Biosystems / Thermo Fisher Scientific | Cat#4374966 |
| TaqMan Fast Advanced Master Mix | Applied Biosystems / Thermo Fisher Scientific | Cat#4444963 |
| PowerPlex 16 HS System | Promega | Technical manual TMD022; product family DC2100/DC2101 |
| Chromium Fixed RNA Profiling (Flex) workflow | 10x Genomics | CG000478 Rev C; CG000527 Rev C |
| Human WTA probe set, BC001 | 10x Genomics | Cat#2000495 |
| Human WTA probe set, BC002 | 10x Genomics | Cat#2000496 |
| Human WTA probe set, BC003 | 10x Genomics | Cat#2000497 |
| Human WTA probe set, BC004 | 10x Genomics | Cat#2000498 |
| REAGENT or RESOURCE | SOURCE | IDENTIFIER |
| Dual Index Plate TS Set A | 10x Genomics | Component#3000511; kit#1000251 |
| SPRIselect cleanup reagent | Beckman Coulter | Cat#B23317 |
| S-Trap spin-column digestion | ProtiFi; University Medical Center Freiburg Proteomics Platform | S-Trap |
| Clarity Max Western ECL substrate | Bio-Rad | Cat#1705062 |
| ImmPACT NovaRED HRP substrate | Vector Laboratories | Cat#SK-4805 |
| **Deposited data** |  |  |
| Bulk RNA-sequencing dataset | This paper | GEO:GSE349777 |
| Chromium Flex single-cell RNA-sequencing dataset | This paper | GEO:GSE350646 |
| GEO SuperSeries | This paper | GEO:GSE350647 |
| Public human HCC single-cell dataset | Liu et al.<sup>17</sup> | GEO:GSE290925 |
| Public HuH-7 single-cell dataset | Zheng et al.<sup>16</sup> | GEO:GSE103867; HuH-7 library GSM2785127 |
| CD133 affinity-purification proteomics | This paper | Table S4; project ST3958/170.1 University of Freiburg Proteomics Core Facility |
| Acute flow-cytometry raw FCS data | This paper | Available upon request |
| Processed data summaries | This paper | Document S1 and Tables S1–S6 |
| Human GRCh38 annotation | GENCODE | Release 50; RRID:SCR_014966; https://www.gencodegenes.org/human/release_50.html |
| Cell Ranger human reference | 10x Genomics | refdata-gex-GRCh38-2020-A; https://www.10xgenomics.com/support/software/cell-ranger/downloads |
| **Experimental models: Cell lines** |  |  |
| Human PLC/PRF/5 cells | ATCC | Cat#CRL-8024; RRID:CVCL_0485; STR report AB240638, 26 March 2024 |
| Human HEK293T cells | Laboratory stock | RRID:CVCL_0063 |
| Human PLC/PRF/5 stable PROM1-CRISPRa population | This paper | N/A; bulk population; available from lead contact |
| Human PLC/PRF/5 SAMph-control population | This paper | N/A; spacer-free control; available from lead contact |
| Human PLC/PRF/5 GFP-population | This paper | N/A; pLJM1-EGFP-transduced population; available from lead contact |
| Human PLC/PRF/5 CD133 ALFA-tag clone 1C4 | This paper | N/A; extracellular insertion; Table S6. |
| Human PLC/PRF/5 CD133 ALFA-tag clone 4B4 | This paper | N/A; cytoplasmic-loop insertion; Table S6. |
| REAGENT or RESOURCE | SOURCE | IDENTIFIER |
| Human umbilical vein endothelial cells | University of Florence | Primary HUVECs |
| **Experimental models: Organisms/strains** |  |  |
| Chicken: Gallus gallus, fertilized SPF eggs | VALO BioMedia | Specific-pathogen-free fertilized eggs |
| **Oligonucleotides** |  |  |
| PROM1 TaqMan Gene Expression Assay | Applied Biosystems / Thermo Fisher Scientific | Assay ID:Hs01009259_m1 |
| CD44 TaqMan Gene Expression Assay | Applied Biosystems / Thermo Fisher Scientific | Assay ID:Hs01075862_m1 |
| CXCR4 TaqMan Gene Expression Assay | Applied Biosystems / Thermo Fisher Scientific | Assay ID:Hs00976734_m1 |
| GPRC5A TaqMan Gene Expression Assay | Applied Biosystems / Thermo Fisher Scientific | Assay ID:Hs01551896_m1 |
| GAPDH TaqMan Gene Expression Assay | Applied Biosystems / Thermo Fisher Scientific | Assay ID:Hs02786624_g1 |
| KRT80 TaqMan Gene Expression Assay | Applied Biosystems / Thermo Fisher Scientific | Assay ID:Hs01372365_m1 |
| PLAU TaqMan Gene Expression Assay | Applied Biosystems / Thermo Fisher Scientific | Assay ID:Hs01547054_m1 |
| SFRP1 TaqMan Gene Expression Assay | Applied Biosystems / Thermo Fisher Scientific | Assay ID:Hs00610060_m1 |
| Human PROM1 siPOOL | siTOOLs Biotech | Cat#PROM1-001 |
| Negative-control siPOOL | siTOOLs Biotech | siPOOL negative control |
| Custom EGFP Flex probe oligonucleotides | Integrated DNA Technologies; this paper | DNA Ultramers; complete ordered sequences in Table S5 |
| PROM1 ALFA-tag knock-in donors and genotyping primers | This paper | Complete available sequences in Table S6 |
| **Recombinant DNA** |  |  |
| LentiSAMPHv2 | Ronald Germain; Chan et al.<sup>15</sup> | Addgene plasmid #167934; RRID:Addgene_167934; https://www.addgene.org/167934/ |
| psPAX2 | Didier Trono / Addgene | Addgene plasmid #12260; RRID:Addgene_12260; https://www.addgene.org/12260/ |
| pCMV-VSV-G | Bob Weinberg; Stewart et al.<sup>40</sup> | Addgene plasmid #8454; RRID:Addgene_8454; https://www.addgene.org/8454/ |
| pLJM1-EGFP | David Sabatini; Sancak et al.<sup>41</sup> | Addgene plasmid #19319; RRID:Addgene_19319; https://www.addgene.org/19319/ |
| PROM1 guide-bearing activation constructs | This paper | GRCh38 chr4:16084166–16084185 and chr4:16084098–16084117 |
| Cas9/guide plasmids for ALFA knock-in | This study | PROM1 sgRNA1 and sgRNA4 constructs; guide sequences in Table S6 |
| pcDNA3.1(+) for genotyping amplicons | Laboratory stock | pcDNA3.1(+) |
| **Software and algorithms** |  |  |
| Cell Ranger | 10x Genomics | Version: 7.2.0; RRID:SCR_017344; https://www.10xgenomics.com/support/software/cell-ranger |
| REAGENT or RESOURCE | SOURCE | IDENTIFIER |
| FastQC | Babraham Bioinformatics | Version: 0.12.1; RRID:SCR_014583; https://www.bioinformatics.babraham.ac.uk/projects/fastqc/ |
| HISAT2 | HISAT2 developers | Version: 2.2.3; RRID:SCR_015530; https://daehwankimlab.github.io/hisat2/ |
| featureCounts | Subread developers | Version: 2.1.1; RRID:SCR_012919; https://subread.sourceforge.net/ |
| R | R Foundation for Statistical Computing | Version: 4.5.3 for bulk RNA-seq; RRID:SCR_001905; https://www.r-project.org/ |
| limma / voom | Ritchie et al.<sup>42</sup>; Law et al.<sup>43</sup> | Version: limma 3.66.0 for bulk RNA-seq; RRID:SCR_010943; https://bioconductor.org/packages/limma/ |
| edgeR | Bioconductor | Version: 4.8.2; RRID:SCR_012802; https://bioconductor.org/packages/edgeR/ |
| Scanpy | Wolf et al.<sup>44</sup> | RRID:SCR_018139; https://scanpy.readthedocs.io/ |
| DoubletDetection | DoubletDetection developers | https://github.com/JonathanShor/DoubletDetection |
| decoupleR / decoupler | Badia-I-Mompel et al.<sup>45</sup> | Univariate linear model (ULM) activity inference |
| CollecTRI regulons | Müller-Dott et al.<sup>46</sup> | https://omnipathdb.org/ |
| MSigDB Hallmark gene sets | Liberzon et al.<sup>47</sup> | Version: 7.5.1 via Enrichr; RRID:SCR_016863; https://www.gsea-msigdb.org/gsea/msigdb/ |
| Enrichr | Ma'ayan Laboratory | RRID:SCR_001575; https://maayanlab.cloud/Enrichr/ |
| Custom preranked GSEA implementation | This paper; Subramanian et al.<sup>39</sup> | 100,000 gene permutations |
| CopyKAT | Gao et al.<sup>48</sup> | Version: 1.2.5; RRID:SCR_024512; https://github.com/navinlabcode/copykat |
| Harmony | Korsunsky et al.<sup>49</sup> | RRID:SCR_022206; https://github.com/immunogenomics/harmony |
| scDblFinder | Germain et al.<sup>50</sup> | RRID:SCR_022700; https://bioconductor.org/packages/scDblFinder/ |
| CellTypist | Domínguez Conde et al.<sup>51</sup> | RRID:SCR_024893; https://www.celltypist.org/ |
| MaxQuant | Cox and Mann<sup>52</sup>; Cox et al.<sup>53</sup> | Version: 2.1.4.0; RRID:SCR_014485; https://www.maxquant.org/ |
| Fiji / ImageJ | Schindelin et al.<sup>54</sup> | RRID:SCR_002285; https://fiji.sc/ |
| Image Lab | Bio-Rad | Version: 6.1 for validation densitometry; RRID:SCR_014210 |
| cellSens Dimension | Olympus | RRID:SCR_014551 |
| FlowJo | BD / FlowJo | Version: 10.6.2 archival workspace; RRID:SCR_008520; https://www.flowjo.com/ |
| Python | Python Software Foundation | Version: 3.12.14 for flow-cytometry reanalysis; RRID:SCR_008394; https://www.python.org/ |
| FlowKit | FlowKit developers | Version: 1.3.2; https://github.com/whitews/FlowKit |
| NumPy | NumPy developers | Version: 2.3.5 for flow-cytometry reanalysis; RRID:SCR_008633; https://numpy.org/ |
| SciPy | SciPy developers | Version: 1.18.1 for flow-cytometry reanalysis; RRID:SCR_008058; https://scipy.org/ |
| Matplotlib | Matplotlib developers | Version: 3.11.2 for flow-cytometry reanalysis; RRID:SCR_008624; https://matplotlib.org/ |
| AlphaFold structural modeling | Jumper et al.<sup>55</sup> | AlphaFold |
| REAGENT or RESOURCE | SOURCE | IDENTIFIER |
| GeneMarker HID | SoftGenetics; authentication provider | GeneMarker HID |
| **Other** |  |  |
| Fluorescence microscope | Olympus | IX73; objective/filter configuration in Methods |
| Fluorescence camera | Olympus | DP74 color CMOS camera |
| LED illumination | CoolLED | pE-300white |
| Flow cytometer | Thermo Fisher Scientific | Attune NxT; instrument identifier 2AFC220520518 |
| Real-time PCR system | Bio-Rad | CFX96 Touch Deep Well |
| Chemiluminescence imager | Bio-Rad | ChemiDoc MP |
| Gel-transfer instrument | Thermo Fisher Scientific | iBlot 2 |
| Mass spectrometer | Proteomics Platform, University Medical Center Freiburg | timsTOF, DDA-PASEF-30SPD |
| LC column for proteomics | Proteomics Platform, University Medical Center Freiburg | Performance 15 cm, EV1137 |

### EXPERIMENTAL MODEL AND STUDY PARTICIPANT DETAILS

#### PLC/PRF/5 cells

The human hepatocellular carcinoma (HCC) cell line PLC/PRF/5 (ATCC, Manassas, VA, USA; catalog no. CRL-8024; RRID:CVCL_0485), an adherent epithelial hepatoma line of male-donor origin also known as Alexander cells, was used for PROM1 perturbation, single-cell profiling, and endogenous tagging. PLC/PRF/5 cells were purchased from ATCC and were reported to test negative for mycoplasma. Cells were maintained at 37 °C in a humidified atmosphere containing 5% CO₂ in high-glucose Dulbecco’s modified Eagle medium (DMEM) supplemented with 10% fetal bovine serum (FBS; Gibco, Grand Island, NY, USA), 100 U/mL penicillin, and 100 µg/mL streptomycin.

PLC/PRF/5 identity was assessed by short tandem repeat (STR) profiling by CLS Cell Lines Service (report AB240638, dated 26 March 2024). STR loci were amplified using the PowerPlex 16 HS System (Promega; DC2101/DC2100); fragment analysis was performed on an ABI3730xl (Life Technologies), and profiles were evaluated using GeneMarker HID software (SoftGenetics). The submitted sample showed a reported 100% match to the PLC/PRF/5 reference DNA profile, with no detectable human cell-line admixture reported.

For PLC/PRF/5 cells, high-glucose DMEM without L-glutamine (Anprotec; catalog no. AC-LM-0011) was supplemented with 1 mM sodium pyruvate (1% v/v of a 100 mM stock; Anprotec; catalog no. AC-DS-0023), penicillin–streptomycin (Anprotec; catalog no. AC-AB-0024), 10 mM HEPES (1% v/v of a 1 M stock; Anprotec; catalog no. AC-DS-000), and 2 mM L-glutamine (1% v/v of a 200 mM stock; Thermo Fisher Scientific; catalog no. 35050-038). Medium was replaced every 2–3 days. At approximately 90% confluence, cells were washed with calcium- and magnesium-free Dulbecco’s phosphate-buffered saline (Anprotec; catalog no. AC-BS-0002) and detached using TrypLE Express (1×, without phenol red; Thermo Fisher Scientific; catalog no. 12604-013) for 5 min at 37 °C. Cells were subsequently reseeded at a 1:4 split ratio.

Stable PROM1-CRISPRa and SAMph-control PLC/PRF/5 populations were used for the CAM grafts and ancillary in-vitro experiments. GFP-reference cells and the ALFA-tagged clones 1C4 and 4B4 were derived from the same parental line. The transcriptomic populations were bulk cultures. The study did not assess sex-dependent effects.

#### HEK293T cells

Human HEK293T cells (RRID:CVCL_0063), derived from the female-origin HEK293 lineage, were used for lentiviral production and maintained at 37 °C in 5% CO2 in high-glucose DMEM supplemented with 10% FBS, 100 U/mL penicillin and 100 μg/mL streptomycin.

#### Primary endothelial cells

Human umbilical vein endothelial cells (HUVECs) were used in the conditioned-medium network-formation assay performed by the University of Florence laboratory. Donor sex was not available for this analysis.

#### Monocytes

Monocytes were used as recipient cells in the University of Florence conditioned-medium migration assay. Donor sex was not available for this analysis.

#### Chicken embryos

Fertilized specific-pathogen-free chicken eggs (Gallus gallus; VALO BioMedia, Osterholz-Scharmbeck, Germany) were used for CAM grafting in two independent repeats and maintained at 37 °C in a humidified incubator without supplemental CO2. Implantation was performed on embryonic day 10 and harvest on day 15. Embryo sex was not recorded. Each egg carried one graft and was allocated and treated separately within its repeat.

CAM experiments confined to unhatched chicken embryos and terminated before hatching fall outside the animal-experiment authorization requirements of Directive 2010/63/EU (Article 1(3)–(4)) and the corresponding German provisions (§ 14 TierSchVersV). Such pre-hatch procedures therefore do not require statutory animal-experiment authorization or notification.

#### Public human HCC data

The GSE290925 series contains data from 12 HCC patients^17^ and describes fresh tumor sampling, 10x Genomics library preparation and NovaSeq 6000 sequencing. This study reanalyzed existing data without collecting new specimens.

### METHOD DETAILS

#### Acute endogenous PROM1 activation

PROM1 was activated using a SAM CRISPRa system,^14^ using the all-in-one LentiSAMPHv2 vector (Addgene plasmid no. 167934; RRID:Addgene_167934). The SAM system combines catalytically inactive Cas9–VP64 with an MS2-modified guide scaffold and the MS2–p65–HSF1 activation module to recruit transcriptional activators without intentionally cleaving genomic DNA. Guide-bearing constructs were generated by BsmBI digestion and insertion of annealed guide oligonucleotides. The activation condition contained a mixture of the Pick2 and Pick4 constructs; the spacer-free backbone served as the SAMph control. Pick2 targeted 5′- GTTGCAAGAAGGGAGTGCAG-3′ adjacent to a 5′-GGG-3′ PAM at GRCh38 chr4:16,084,166–16,084,185, and Pick4 targeted 5′-GTGGGCGCCCCAGGGCTACG-3′ adjacent to a 5′-TGG-3′ PAM at chr4:16,084,098–16,084,117. Coordinates are 1-based and inclusive; both protospacers map to the minus strand and are centered approximately 164 and 96 bp upstream of the predominantly used transcription start site. For transient transfection, 300,000 cells were plated per six-well plate well, followed the next day by 2.5 μg plasmid DNA, 5 μL P3000 and 3.75 μL Lipofectamine 3000. Complexes were prepared in Opti-MEM for 15 min, and cells were collected after 48 h. Bulk sequencing comprised three samples per group.

The ordered cloning oligonucleotides were Pick2 forward, 5′- CACCGGTTGCAAGAAGGGAGTGCAG-3′, and reverse, 5′-AAACCTGCACTCCCTTCTTGCAACC-3′; and Pick4 forward, 5′- CACCGGTGGGCGCCCCAGGGCTACG-3′, and reverse, 5′-AAACCGTAGCCCTGGGGCGCCCACC-3′. The CACC/AAAC ends and additional guide-expression/cloning bases are distinct from the 20-nucleotide target sequences given above.

#### PROM1 knockdown

PLC/PRF/5 cells were seeded in six-well plates at 3 × 10^5^ cells per well one day before transfection. Cells were transfected with a human PROM1-targeting siPOOL (siTOOLs Biotech; catalog no. PROM1-001) or the siTOOLs negative-control siPOOL using Lipofectamine RNAiMAX (Invitrogen, Thermo Fisher Scientific; catalog no. 13778-150) in a forward-transfection procedure. Before transfection, the culture medium was replaced with 2 mL Opti-MEM I Reduced Serum Medium (Thermo Fisher Scientific; catalog no. 31985-062). For each well, 40 μL of a 1 μM siPOOL working solution was diluted in 210 μL Opti-MEM. Separately, 4 μL Lipofectamine RNAiMAX was diluted in 246 μL Opti-MEM. The two solutions were combined and incubated for 15 min at room temperature to allow complex formation. The resulting 500 μL mixture was added to each well, delivering 40 pmol total siRNA at a nominal final concentration of 16 nM in 2.5 mL. Negative-control transfections were performed under the same conditions. Cells were harvested 48 h after transfection for RNA extraction and subsequent RNA sequencing and RT–qPCR. Each analysis included three samples per condition.

#### Bulk RNA preparation and sequencing

Bulk RNA sequencing was performed on PLC/PRF/5 cells following acute CRISPRa-mediated activation of endogenous PROM1 and on the corresponding spacer-free SAMPH controls, with three biological replicates per condition. Cells were collected 48 h after transfection, washed with calcium- and magnesium-free PBS, harvested using a cell scraper and pelleted at 350 × g for 5 min at 4°C. Total RNA was isolated using the RNeasy Plus Mini Kit (Qiagen, catalog no. 74134) according to the manufacturer’s instructions. RNA concentration and purity were assessed using a DS-11 FX instrument (DeNovix), and RNA integrity was evaluated using an Agilent 4200 TapeStation with RNA ScreenTape. Recorded RNA integrity number equivalent (RINe) values exceeded 9.

Library preparation and sequencing were performed by Eurofins Genomics, Konstanz, Germany, using the INVIEW Transcriptome service. Six total-RNA samples were submitted in water, and all passed the provider’s entry quality control. Ribosomal RNA depletion was used. Strand-specific RNA-sequencing libraries were prepared.

Libraries were sequenced on an Illumina NovaSeq instrument in paired-end mode with a nominal read length of 2 × 150 bp. Sequencing yielded approximately 8.61–24.88 million read pairs per sample. Paired FASTQ files and MD5 checksums were supplied for downstream analysis.

The workflow assessed reads with FastQC 0.12.1 and performed no additional local trimming. Reads were aligned to GRCh38 with HISAT2 2.2.3 using GENCODE v50 splice sites, and reverse-stranded paired fragments were counted with featureCounts 2.1.1. Differential expression used limma 3.66.0 and edgeR 4.8.2 in R 4.5.3, with filterByExpr, TMM normalization, voom, a condition model and empirical Bayes moderation. Gene-level p values were Benjamini– Hochberg-adjusted within each arm. Each perturbation was compared with its corresponding control, using adjusted p < 0.05 for the reported 452 activation and 3,359 knockdown differentially expressed genes. The reciprocal-response analysis first retained genes with adjusted p < 0.01 in both contrasts, giving 124 genes, and then evaluated their signs, yielding 117 with opposing directions.^42,43^

#### Promoter and off-target analyses

Aligned reads were inspected across PROM1 on GRCh38 using GENCODE release 50. The locus model used the union of annotated exons, and alternative first exons were labeled by their transcription start coordinates and the number of annotated transcripts initiating there. Coverage was pooled across three libraries per condition, scaled per million assigned fragments and displayed on a common vertical scale. The guides targeted chr4:16,084,166–16,084,185 and chr4:16,084,098–16,084,117, each with one perfect genomic match. The proximal TSS used in PLC/PRF/5 was approximately chr4:16,084,012, close to the MANE Select TSS at 16,084,023 (Figure S1A and S1B).

For the local transcriptional analysis, seven expressed neighbors within ±1 Mb of PROM1 (chr4:15.0–17.1 Mb) were plotted at their TSSs. Each gene was compared with the central 95% of log2 fold changes among 1,194–1,870 genome-wide genes of comparable expression; these are descriptive expected ranges, not confidence intervals. The mean absolute fold change of the seven neighbors was assessed against 2,000 expression-matched permutations, yielding an empirical p value of 0.58. Splice junctions in the window were screened for crossing of the PROM1 gene boundary (Figure S1C).

For the separate genome-wide predicted off-target analysis, guide matches were enumerated through four mismatches with an NGG PAM. Candidate sites within 1 kb of an annotated transcription start site were linked to genes, yielding 265 expressed candidates and 249 with differential-expression test results. Candidate-set log2 fold changes were compared with expression-matched gene sets using 2,000 permutations. Table S1 also reports tests for the core promoter window (−400 to −50 bp relative to the TSS), chromosome 4 outside the PROM1 ±1 Mb region, and the Spearman association between promoter similarity and log2 fold change across 15,704 genes.

#### Quantitative PCR and protein validation

Cells were collected 48 h after transfection for RNA extraction and downstream analyses. RNA was isolated using the RNeasy Plus Mini Kit (Qiagen; catalog no. 74134) according to the manufacturer’s instructions. RNA concentration and purity were assessed using a DS-11 FX spectrophotometer/fluorometer. Samples submitted for RNA sequencing underwent additional quality assessment using an Agilent 4200 TapeStation, and only samples with an RNA integrity number greater than 9 were processed. RNA sequencing included three samples per treatment group. TaqMan qPCR measured PROM1, GPRC5A, CXCR4, CD44, PLAU, KRT80 and SFRP1, with technical triplicates averaged before normalization to GAPDH by the ΔΔCt method. Relative values were scaled to the mean of the corresponding control group. Activation qPCR comprised six wells per condition from two independent transfections (three wells per transfection); RNA from one transfection was also used for the bulk sequencing experiment. Knockdown qPCR comprised three wells per condition (Figure S1D and S1E).

For reverse transcription, 400 ng total RNA in 10 μL nuclease-free water was combined with 10 μL reaction mix from the High-Capacity cDNA Reverse Transcription Kit (Applied Biosystems, Thermo Fisher Scientific; catalog no. 4374966). The 20 μL reaction contained 2 μL 10× RT buffer, 0.8 μL 25× dNTP mix, 2 μL 10× random primers, 1 μL MultiScribe reverse transcriptase and 1 μL RNase inhibitor, with nuclease-free water to volume. Reverse transcription was performed at 25 °C for 10 min, 37 °C for 120 min and 85 °C for 5 min; cDNA was stored at −20 °C.

TaqMan reactions contained 1 μL undiluted cDNA, 5 μL 2× TaqMan Fast Advanced Master Mix (Applied Biosystems; catalog no. 4444963), 0.5 μL 20× target assay, 0.16 μL 60× GAPDH reference assay and 3.34 μL nuclease-free water, giving a total volume of 10 μL. Reactions were run in technical triplicate on a CFX96 Touch Deep Well Real-Time PCR Detection System (Bio-Rad), with 95 °C for 2 min followed by 40 cycles of 95 °C for 5 s and 60 °C for 30 s, with fluorescence acquisition at the end of each cycle. Commercial TaqMan Gene Expression Assays (Applied Biosystems/Thermo Fisher Scientific) included Hs01009259_m1 for PROM1, Hs01551896_m1 for GPRC5A, Hs01075862_m1 for CD44, Hs00976734_m1 for CXCR4, Hs01372365_m1 for KRT80, Hs01547054_m1 for PLAU, Hs00610060_m1 for SFRP1 and Hs02786624_g1 for GAPDH.

Immunoblot validation used three separately transfected wells per condition in one experiment for each perturbation arm, with one lysate per well and alternating perturbation and control lanes. For activation, CD133 and β-actin were probed on one membrane; CD44, CXCR4 and KRT80 were assessed on separate membranes loaded with the same lysates and normalized to the β-actin signals from the parallel blot. CXCR4 quantification summed the two upper immunoreactive bands and excluded the lowest band. Band volumes were measured in Image Lab 6.1 with rolling-disk background subtraction (10 mm); the analyzed images had no saturated pixels. Target signals were divided by β-actin and scaled to the corresponding control mean. The same normalization was used for knockdown CD133, with the six displayed and quantified lanes from the right-hand portion of the original nine-lane membrane (Figure S1F and S1G).

#### Stable populations and co-culture

Lentivirus was produced in HEK293T cells in six-well plates by co-transfecting 1 µg of the respective transfer plasmid preparation (Pick2/Pick4 LentiSAMPHv2 constructs for PROM1 activation, spacer-free LentiSAMPHv2 for SAMph controls, or pLJM1-EGFP for GFP-reference cells), 1 µg of the packaging plasmid psPAX2 (Addgene plasmid no. 12260), and 0.5 µg of the envelope plasmid pCMV-VSV-G (Addgene plasmid no. 8454), using Lipofectamine 3000 (Thermo Fisher Scientific). The total DNA input was 2.5 µg per well.

Virus-containing culture supernatant was collected 3 days after transfection, clarified by centrifugation at 3,000 × g for 5 min, and passed through a 0.45 µm filter. PLC/PRF/5 cells were transduced using 300 µL of the clarified, filtered supernatant per culture. Guide-bearing PROM1 activation populations and SAMph control populations were selected with 20 µg/mL blasticidin (InvivoGen; ant-bl-05). GFP-reference cells were generated using pLJM1-EGFP (Addgene plasmid no. 19319) and selected with 5 µg/mL puromycin (Thermo Fisher Scientific; A1113803).

For single-cell profiling, control or PROM1-activated target cells were mixed with GFP-reference cells at a target-to-reference ratio of 3:1 and co-cultured for 72 h in T75 flasks. Four independently transduced cultures were prepared per condition and pooled pairwise after fixation, generating two measured pools per condition.

#### Flex library preparation

After co-culture, cells were fixed in approximately 4% formaldehyde for 20 h at 4°C, quenched, incubated with enhancer and held for 16 h at 4°C. Two independently transduced cultures from the same condition were then pooled, yielding two samples per condition and four samples in total. Fixed cells were processed using the Chromium Fixed RNA Profiling (Flex) workflow (10x Genomics Demonstrated Protocol CG000478 Rev C and User Guide CG000527 Rev C). To identify the pLJM1-EGFP reference cells, three custom left-hand probes (LHS-1–3) targeting EGFP were paired with three sample-specific right-hand probes (RHS-1–3) carrying the corresponding sample barcode (BC001–BC004). The probe oligonucleotides were ordered as DNA Ultramers (Integrated DNA Technologies) and designed according to the 10x Genomics Custom Probe Design Technical Note (CG000621 Rev C); their ordered sequences are listed in Table S5. Oligonucleotide stocks (100 μM) were diluted in IDTE, pH 8, to a 40 nM working solution. For each sample, 1 × 10^6^ cells were resuspended in 80 μL hybridization mix; 20 μL of the corresponding barcoded human whole-transcriptome analysis probe set and 5 μL of the custom EGFP probe mixture (three common LHS probes plus the matching three RHS probes) were added, giving a final concentration of approximately 1.9 nM per custom probe. Hybridization was performed for 20 h at 42°C. After hybridization, the four barcoded samples were pooled, washed, filtered through a 30-μm strainer and resuspended for GEM generation. Chromium iX partitioning was performed with a Next GEM Chip Q, targeting approximately 20,000 recovered cells. Libraries were prepared following the manufacturer’s Fixed RNA Profiling workflow and sequenced on a NovaSeq 6000 SP flow cell. The pooled library concentration was 1.0 nM, the loading concentration was 200 pM, and 15% PhiX was included.

For fixation, 1.5 × 10^6^ cells were washed in PBS containing 0.04% BSA and resuspended in 1 mL fixation buffer containing 791.9 μL nuclease-free water, 100 μL concentrated Fix & Perm Buffer (10x Genomics; 2000517) and 108.1 μL 37% formaldehyde (Sigma-Aldrich; 252549). After fixation, cells were centrifuged at 850 × g for 5 min, resuspended in 1 mL quenching buffer prepared using concentrated Quench Buffer (10x Genomics; 2000516), and supplemented with 100 μL Enhancer (10x Genomics; 2000482). Cell suspensions were passed through 30 μm pre-separation filters (Miltenyi Biotec; 130-041-407). Cell counts and aggregate formation were assessed using a LUNA-FX7 automated cell counter with 2 μL acridine orange/propidium iodide stain added to 18 μL cell suspension. The Human WTA probe sets BC001–BC004 were 10x Genomics products 2000495–2000498.

The recorded Flex pre-amplification program was 98 °C for 3 min, followed by eight cycles of 98 °C for 15 s, 67 °C for 20 s and 72 °C for 1 min, with a final 72 °C extension for 1 min. Index PCR used Dual Index TS Set A (10x Genomics; plate 3000511, kit 1000251) and comprised 98 °C for 45 s, followed by eight cycles of 98 °C for 20 s, 54 °C for 30 s and 72 °C for 20 s, with a final 72 °C extension for 1 min. SPRIselect (Beckman Coulter; B23317) cleanup was performed after pre-amplification and index PCR.

#### Flow cytometry after acute activation

PLC/PRF/5 cells were analyzed 48 h after transient transfection with PROM1-targeting SAM CRISPRa or SAMph control plasmids in a single experiment. 5 µL anti-CD44–Brilliant Violet 421 (clone IM7; Invitrogen, 404-0441-82), 5 µL anti-CD133–PE (clone TMP4; Invitrogen, 12-1338-42) and 1 µL SYTOX Red were applied, with incubation for 30 min at 4°C. Samples were acquired on an Attune NxT Acoustic Focusing Cytometer (Thermo Fisher Scientific). CD133–PE, SYTOX Red and CD44–Brilliant Violet 421 signals were recorded in BL2-A, RL1-A and VL1-A with 488-, 640- and 405-nm excitation, respectively. Detector voltages were FSC, 120 V; SSC, 310 V; BL2, 320 V; RL1, 280 V; and VL1, 220 V. Acquisition flow rates were 200 µL/min for SAMph and 500 µL/min for CRISPRa, yielding 200,000 and 341,544 recorded events. Unstained, single-marker, SYTOX-only, marker-minus and acquisition compensation controls accompanied one fully stained tube per group. Source labels SAM and CRISPR correspond to the SAMph and acute PROM1-CRISPRa conditions, respectively.

The displayed analysis was performed retrospectively. A shared SYTOX-low cutoff was derived from two SYTOX-only controls after scatter and singlet selection. RL1-A values were transformed as asinh(x/150), binned from −7 to 10 in 0.05-unit increments and normalized within this range. The equally weighted group histograms were smoothed with a Gaussian kernel (standard deviation 2 bins; reflected boundaries); the minimum between the low mode (x < 3) and high mode (4 < x < 8) gave a back-transformed upper cutoff of 6262.562475805535. No lower fluorescence cutoff was applied. Selected events are described as SYTOX-low (putatively viable). CD133-plus-SYTOX and CD44-plus-SYTOX tubes served operationally as CD44 and CD133 fluorescence-minus-one (FMO) controls, respectively. For each marker, the higher of the two group-specific 99.5th percentiles after scatter, singlet and SYTOX-low gating was applied to both groups (NumPy linear/type 7 quantile). Positive thresholds were VL1-A ≥ 359.35306965855244 for CD44 and BL2-A ≥ 722.1255760393983 for CD133. Gates were evaluated in native compensated units; asinh(x/150) was used for display (Figure 3F).

The final denominators were 119,536 SAMph and 183,881 CRISPRa SYTOX-low singlets. Four mutually exclusive marker classes were quantified; total marker positivity included the corresponding single-positive and double-positive classes. Analysis used Python 3.12.14, FlowKit 1.3.2, NumPy 2.3.5, SciPy 1.18.1 and Matplotlib 3.11.2. Sensitivity analyses varied marker thresholds, SYTOX selection, upstream gates, compensation and historical reconstruction.

#### Single-cell processing and quality control

Sequencing was performed on an Illumina NovaSeq 6000. FASTQ files were generated from the raw run data using Cell Ranger 7.2.0 mkfastq with the sample sheet and the options -force-single-index and -barcode-mismatches 2, allowing up to two sample-index barcode mismatches during demultiplexing. Gene-expression processing then used Cell Ranger multi with a configuration CSV specifying refdata-gex-GRCh38-2020-A and the custom Chromium_Human_Transcriptome_Probe_Set_v1.0.1_GRCh38-2020-A_GFP.csv probe file, which supplemented Human Transcriptome Probe Set v1.0.1 with three EGFP probe pairs (Table S5). The no-bam option was set to true. Of 20,011 initial assignments, 292 high-occupancy assignments representing 73 GEMs across four barcodes were removed, leaving 19,719 cell calls and 18,083 targeted genes. The two control pools contained 4,065 and 5,628 cells, and the activated pools 4,517 and 5,509 cells before EGFP and subsequent quality filtering.

Cells with ≥1 EGFP count were excluded, removing 2,650 controls and 3,810 activated cells. DoubletDetection was applied per sample with 10 iterations, p threshold 1 × 10^−1^^6^, voter threshold 0.5 and seed 0. Cells with fewer than 1,500 detected genes, more than 10% mitochondrial counts, or upper outlier log UMI or gene counts beyond five median absolute deviations were excluded. Mitochondrial fractions used the 11 mitochondrial targets in the Flex panel. The final sample counts were 2,820 and 3,769 controls, and 2,820 and 3,078 activated cells.

#### Embedding and state robustness

Counts were normalized to 10,000 per cell and log1p-transformed. Variable genes were selected by the Seurat v3 method with sample as batch key, with mitochondrial genes excluded, leaving 1,989 of 2,000 selected genes. S and G2M scores were regressed only for the embedding representation. Values were scaled and clipped at 10; 50 principal components were computed, with 30 used for a 15-neighbor graph. UMAP and Leiden clustering used seed 0, and resolution 0.5 produced six clusters. Conditions were not integrated, and unregressed values were retained for expression analyses. C0 was identified as the cluster with at least 100 cells and the largest decrease in representation after activation.^44^

Robustness analyses repeated Leiden clustering at resolutions 0.3, 0.5, 0.8, 1.0 and 1.2, with three alternative random seeds and five 80% cell subsamples. Preprocessing was also repeated without cell-cycle regression or with PROM1 excluded from variable-gene selection and PCA. In each variant, the state was reselected as the cluster with at least 100 cells showing the largest decrease from control to GoF. Correspondence was quantified as the fraction of cells in the selected state belonging to original C0. Fold differences in Figure S2C used cell-weighted condition fractions; at higher resolution, the selected state could be a subset of original C0.

Figure S2F compared the original Leiden assignment with four alternative estimates. Count splitting used separate count partitions for state definition and evaluation across three random seeds. The frozen-control reference fitted feature selection, cell-cycle regression, scaling and PCA using controls only, projected activated cells, and assigned a cell to C0 when more than half of its 30 nearest control neighbors carried the C0 label. Control-derived expression signatures used 50 C0-enriched and 50 rest-enriched genes selected by pseudobulk limma-voom (FDR < 0.05, |log2 fold change| > 1, concordant direction in both control pools and excluding PROM1). The score contrasted the two gene sets, and the classification threshold was chosen at the Youden point in controls. A further signature excluded genes differentially expressed in the acute bulk activation analysis at FDR < 0.05. Fold differences in Figure S2F were ratios of unweighted mean pool fractions. These estimates reuse control-derived labels and are complementary sensitivity analyses, not independent biological replications.

S and G2M scores and phase assignments used scanpy score_genes_cell_cycle on log-normalized counts with the Tirosh cell-cycle gene sets; Table S2 lists the 42 S-phase and 54 G2M input genes and their use in the expression matrix. All 42 S-phase genes and 52 G2M genes were used; FAM64A and HN1 were not matched under the supplied names. The reciprocal-response score used the 107-gene set defined below. Absolute Spearman correlations of the S, G2M and response scores with PC1–PC10 were calculated before and after cell-cycle regression, together with direct score–score correlations (Table S2). Figure S2H shows the corresponding phase and response-score embeddings. Cycling cells were those assigned to S or G2M (Figure S2E and S2G). Quality overlays showed detected genes, UMIs and mitochondrial-count fractions with color scales limited to the 1st–99th percentiles; C0-versus-other comparisons were stratified by pool (Figure S2D). Mitochondrial fractions refer to the targets included in the Flex probe panel. All state fractions and diagnostics were descriptive, with two measured pools per condition.

CD44 detection sensitivity was assessed descriptively in all quality-controlled cells (6,589 control and 5,898 PROM1-CRISPRa cells) at thresholds of ≥1, ≥2, ≥3 and ≥5 CD44 UMI. Detection at ≥1 UMI was also summarized within quintiles of total UMI counts per cell, using common strata across both conditions. For count-depth sensitivity, cells below the pooled tenth percentile were excluded and the remaining cells were downsampled to 6,532 UMI per cell using Scanpy downsample_counts; detection was then reassessed at ≥1 and ≥2 UMI. Each analysis was additionally repeated among cells outside C0. Values were pooled across the two measured pools per condition and are reported descriptively without inferential testing (Table S2).

#### Pseudobulk and pathway analyses

For the main C0-versus-rest comparison, raw counts were summed within each control pool and state, yielding four pseudobulks. Analysis used limma-voom with filterByExpr, TMM normalization and a design containing pool and state terms. Fold changes are oriented as remaining cells relative to C0. The Figure 2C display required Benjamini–Hochberg FDR < 0.05, |log2 fold change| > 1, mean expression ≥0 log2 CPM and concordant direction in both pools, identifying 219 C0-enriched and 534 rest-enriched genes. State comparisons used the pool as the replication unit.^42,43^ Table S2 reports all 753 eligible genes, with pool-specific fold changes and the 15 labels per direction in Figure 2C.

Hallmark gene sets (the MSigDB_Hallmark_2020 Enrichr library) were evaluated by preranked GSEA on limma-voom moderated t statistics, using the shared weighted-running-sum implementation for the bulk arms and control-state comparison. Sets required at least 15 measured genes. With 100,000 gene permutations, p values were calculated as (b + 1)/(m + 1), where b is the number of null scores at least as extreme and m is the number with the observed sign; Benjamini–Hochberg correction was applied across tested sets. Figures 1C and 1D display the five significant sets with the largest absolute NES per direction at GSEA FDR < 0.05; complete acute-arm results are provided in Table S1. Correlation Adjusted MEan Rank (camera) on the same voom model (inter-gene correlation 0.01) was a separate comparison and was not used to select the Figure 1 display. Figure 2F instead displays the five sets with the largest absolute GSEA NES per direction among sets meeting both GSEA FDR < 0.05 and fixed-correlation camera FDR < 0.05, with concordant direction. Bars show GSEA NES and q labels show GSEA FDR. For example, the acute GoF EMT set had GSEA FDR approximately 0.0002 but camera FDR 0.17.^39,47^ The complete 50-set control-state analysis is provided in Table S2, together with camera using estimated inter-gene correlation and a fry sensitivity analysis. Estimated-correlation camera yielded FDRs of 0.55–0.92; these sensitivity outputs were not used to select the Figure 2F display.

Control-cell transcription-factor activities used CollecTRI with decoupler ULM on log-normalized expression, centered within each cell by the median activity across regulons. Among regulons with at least 25 measured targets, nine per direction were selected by the difference between upper- and lower-decile mean activity, averaged over the two control pools. Bulk ULM enrichment scores used the corresponding moderated t statistics. Directional mroast tests used signed CollecTRI weights, the mean set statistic and 99,999 rotations; cameraPR used inter-gene correlation 0.01 and sign inversion for repressed targets. Each test applied Benjamini– Hochberg correction over the 18 regulons per arm. Both tests had to reach FDR < 0.05 in the score direction for a dark bar in Figure 2G.^45,46^ Tables S1 and S2 provide all 343 axis-evaluated regulons, with the single-cell target coverage and bulk ULM results. Joint mroast/cameraPR flags are supplied only for the 18 displayed regulons; an em dash in the other rows denotes an unevaluated joint criterion rather than a negative result. FOXL2 has bulk ULM scores but no genome-wide adjusted ULM p value because it fell below the bulk target-size threshold; its restricted Figure 2G tests did not meet the joint criterion.

For the separate acute-arm regulon summaries in Table S1, reported ULM enrichment scores were calculated from the bulk moderated-t contrasts. Table S1 reports all evaluated ULM scores and adjusted p values, MLM scores and expression changes of the factors themselves; the joint mroast/cameraPR criterion applies to the displayed bars in Figure 2G.

#### Response signatures and state coordinate

For PLC/PRF/5 single-cell analyses (Figures 2D, 2E and S2H), the reciprocal-response signatures were derived from genes with adjusted p < 0.01 in both acute bulk contrasts and opposite log2 fold-change signs. Of 117 reciprocal genes (Table S1), PROM1 and nine targets not covered in the single-cell analysis were excluded, leaving 34 activation-up/silencing-down and 73 activation-down/silencing-up genes. No additional fold-change cutoff was imposed. Log-normalized expression was standardized per gene across all displayed cells; each component score was the mean of its gene-wise z-scores. The combined score was the increased-component mean minus the decreased-component mean. Figure 2D shows illustrative gene profiles along the control coordinate, and Figure 2E shows embeddings of the combined response score and state coordinate; the selected heatmap rows do not define the full score. The human-tumor score panels (Figures 4E–4G) used the same 107-gene reciprocal signature, with 34 up-with-PROM1 and 73 down-with-PROM1 genes and standardization within the stated human-cell analysis population. Figure S6 used the 85 signature genes outside the displayed axis-gene set, as described below. PROM1 itself was excluded from both component scores. These signatures characterize the independently constructed control-cell diffusion coordinate.

The root-free coordinate was the first nontrivial diffusion component of the control-cell graph (Scanpy diffmap; 15 neighbors, 30 PCs, 15 components, seed 0), constructed in the shared PCA representation. Its sign was oriented towards increasing endogenous PROM1 expression, placing C0 at the low end. Control quantiles were (average rank − 0.5)/number of controls. Activated-cell projections averaged the signed diffusion coordinates of 30 nearest control neighbors in PCA space before applying the empirical control cumulative distribution. Control self-projection used 30 control neighbors excluding the cell itself. Occupancy comparisons and the stated interval fractions used projected coordinates in both conditions. The reference quantile is approximately uniform by construction and does not establish a density gap. Figure 3C uses 60 equal-width bins, Gaussian smoothing with SD 1.5 bins for display and unweighted means across pools.

Cells were flagged outside reference neighborhoods when their mean distance to the 30 nearest control neighbors exceeded the 95th percentile of control leave-one-out neighbor distances; 14.2% of activated cells met this criterion. The main occupancy curves include these cells, with exclusion assessed separately.

For the PLC/PRF/5 coordinate robustness checks (Table S2), Spearman correlations with UMI count, detected-gene count, mitochondrial fraction, doublet score, residual S/G2M scores and marker expression were computed in all control cells and within each pool. Marker associations were additionally evaluated as partial rank correlations after removing the linear dependence on log10 UMI count. Pool-associated variation was summarised by the two-sample Kolmogorov–Smirnov statistic and the fraction of coordinate variance explained by pool (η^2^). Recomputed variants used 10 or 30 neighbours, 20 or 40 PCs, exclusion of the acute reciprocal-response genes including PROM1 from the variable-gene set, additional regression of log10 UMIs and mitochondrial fraction, or regression, scaling and PCA fitted on control cells alone, with and without signature exclusion. The first three nontrivial diffusion components were evaluated to detect component swaps; reference correlations and sign-aligned marker correlations are reported descriptively, without treating cells as independent experimental replicates.

For the cross-experiment analysis in Figure S3, limma-voom tables were reduced to one entry per gene symbol, retaining the entry with the highest average expression. Genes with Benjamini–Hochberg-adjusted p < 0.01 in both acute arms formed a shared set of 124 genes, of which 117 had opposite and seven concordant signs. Single-arm sets comprised all genes meeting this threshold in GoF (215) or LoF (2,056), regardless of significance in the other arm. After restriction to the single-cell probe panel and exclusion of PROM1 from correlations, the shared, GoF and LoF sets contained 112, 182 and 1,823 genes, respectively. PROM1 was displayed separately in the heatmap. CD44 was included as an additional reference row outside the shared set; it was not included in the shared-set correlations or the reciprocal score. Figure S3A therefore contains 114 displayed rows (113 measured shared genes plus CD44).

Axis profiles used unregressed log1p counts per 10,000 from the 6,589 control cells. The control-quantile coordinate was divided into 40 equal-width bins. Mean expression was calculated within each pool and bin; bins containing fewer than 25 cells in a pool were masked, pool profiles were averaged, and each gene was z-scored across bins. Rows were ordered by the bin of maximum expression within each response block. Gene-wise axis association was the Spearman correlation between expression and coordinate within each control pool, using average ranks for ties and then averaging the two pool-specific correlations. Spearman correlations across genes summarized agreement between these associations and bulk log2 fold changes. Because genes were selected using the bulk results, are co-expressed, and were evaluated along an axis derived from the same control cells, these correlations were descriptive and no p values were reported.

#### Reanalysis of HuH-7 single-cell data

The published 10x Genomics count matrix from GSE103867 was used to extract the HuH-7 library (GSM2785127; barcode suffix -2; Zheng et al., 2018)^16^. Counts for duplicate gene symbols were summed. Cells with at least 500 detected genes and less than 20% mitochondrial counts were retained (2,075 of 2,082 cells). The legacy annotation lists ADGRG6 as GPR126, SELENOP as SEPP1 and ARFGEF3 as KIAA1244; CD24 and SLC38A3 are absent. Signature scores used genes present under the current symbols, whereas former symbols were mapped for gene-level heatmaps.

Counts were normalised to 10,000 per cell and log1p-transformed. S and G2M scores and phases were assigned with Scanpy score_genes_cell_cycle using the Tirosh et al.^56^ gene lists. A 15-neighbour graph and Leiden clustering at resolution 0.5 separated two disconnected transcriptional compartments (clusters 0–2, 1,274 cells; clusters 3–4, 801 cells), treated as subpopulations without inferring genetic clonality. Gene-set scores were means of gene-wise z-scored log1p-normalised expression, omitting genes without variance. The 117-gene reciprocal response yielded 107 genes after PLC assay restriction and PROM1 exclusion, of which 31 activation-upregulated and 68 activation-downregulated genes were measurable in HuH-7. The C0 marker sets were the 50 strongest genes per direction in the PLC/PRF/5 control pseudobulk contrast, ranked by moderated t after requiring FDR < 0.05, |log2 fold change| > 1, concordant pool-specific directions and mean log2 CPM ≥ 0, with PROM1 excluded. Only genes present and variable in HuH-7 contributed to the marker scores. The full reciprocal-response and C0 marker lists shared 12 genes.

Within each subpopulation, 2,000 highly variable genes were selected from counts using the Seurat v3 procedure, excluding mitochondrial genes. S and G2M scores were regressed out, values were scaled with a maximum of 10, and 50 principal components were computed. A graph using 15 neighbours and 30 PCs supported a 15-component diffusion map. The first nontrivial component defined a root-free coordinate, converted to a rank quantile within each subpopulation. Its sign was oriented so that the correlation with the C0-minus-remaining-cell marker score was negative; PROM1 increased in this orientation. This choice determined the sign rather than the cell ordering. Pooled displays combined cells by their within-subpopulation quantiles.

Coordinate robustness was examined using correlations with log10 UMIs, detected-gene counts, mitochondrial fraction, residual cell-cycle scores and PROM1, CD44 and APOB expression, both pooled and within subpopulations (Table S3). Partial marker correlations adjusted for log10 UMIs; cell-cycle-phase effects were summarised as η^2^. Variants changed the neighbour count to 10 or 30, removed the union of the 117 reciprocal-response genes (including PROM1) and 100 C0 markers (205 unique genes) from the variable-gene space, additionally regressed log detected-gene and log UMI counts, omitted cell-cycle regression, or used G1-phase cells alone. Reference correlations were evaluated across the first three diffusion components to detect component swaps (Table S3). Further analyses used depth-adjusted partial rank correlations, a linear model including PROM1, depth, cell-cycle scores and subpopulation with HC3 standard errors, Mann–Whitney comparisons of signature scores at PROM1 thresholds of 1, 2, 3 or 5 UMIs versus zero, and within-phase associations with and without regression (Table S3).

PROM1-high cells (≥3 UMIs, n = 193) were compared with PROM1-undetected cells (n = 1,077) using limma-trend on log1p-normalised expression. Genes detected in more than 1% of compared cells were retained, with PROM1 excluded. The model included group, log detected-gene count, S score, G2M score and subpopulation; subpopulation-specific analyses omitted the latter term. Moderated-t statistics ranked genes for preranked GSEA (100,000 permutations, empirical plus-one correction and Benjamini–Hochberg adjustment across sets) and camera (inter-gene correlation 0.01). Hallmark sets and both directions of the full reciprocal-response signature after PROM1 exclusion were evaluated with the same implementation used for the PLC analyses; these sets were included in one Benjamini–Hochberg adjustment family per contrast. Displayed Hallmarks required GSEA FDR < 0.05 with camera support in the adjusted pooled comparison; row-specific support is indicated separately in Figure 4D.

CollecTRI regulons were evaluated with decoupler ULM and MLM on the moderated-t statistics, requiring at least 15 measured targets and Benjamini–Hochberg adjustment across regulons. Directional cameraPR used inter-gene correlation 0.01 and inverted the signs of repressed targets. For the 18 regulons selected to describe the PLC axis, of which 17 were measurable in HuH-7, weighted mroast used the mean set statistic and 99,999 rotations, with cameraPR corroboration and Benjamini–Hochberg adjustment within this family. To compare cell lines on the same scale in Figure 4C, the same ULM/cameraPR procedure was applied to the PLC remaining-cell-versus-C0 moderated-t contrast (limma-voom; four pseudobulks from two control pools; model ∼ pool + group). Thus, this comparison uses contrast-based ULM enrichment scores rather than the axis-position differences displayed in Figure 2G.

Cross-model concordance was summarised across all 50 Hallmark sets and 312 jointly measured regulons (Table S3). For 9,539 genes detected in more than 1% of cells in both lines, excluding PROM1, expression–coordinate Spearman correlations were calculated in the 6,589 PLC/PRF/5 control cells and 2,075 HuH-7 cells. Agreement across genes was compared with 10,000 permutations of gene labels within ten detection-rate strata. Genes with |ρ| > 0.3 in PLC/PRF/5 (145 genes) were displayed over 40 coordinate bins, with PLC profiles first averaged within each pool and then across pools, followed by gene-wise standardisation across bins. Directional concordance and cross-gene correlation were accompanied by 95% confidence intervals from 2,000 gene bootstrap resamples. A sign-weighted mean z-score of these 145 genes was evaluated against the HuH-7 coordinate and PROM1 at the cell level; Pearson correlation of score and PROM1 means over 40 bins was reported separately. A sensitivity projection used the 100 genes at each end of the PLC axis after excluding the response-signature and C0 marker genes.

#### Public data analysis and case selection

The public-data analysis started from 149,021 barcodes and 36,601 features in GSE290925. Cells with fewer than 200 genes, fewer than 500 UMIs or more than 20% mitochondrial counts were removed; high UMI and gene-count outliers were identified using a five-MAD threshold on log-transformed values. scDblFinder was applied per patient, leaving 121,556 cells. Genes detected in at least three cells were retained (32,457 genes). Counts were normalized to 10,000 and log1p-transformed. Two thousand variable genes were selected with patient as the batch key, excluding mitochondrial genes. Fifty principal components were calculated, the first 30 were integrated with Harmony, and a 15-neighbor graph and Leiden resolution 1 were used for the cohort overview.^49,50^

For the epithelial-candidate overview, genes detected in at least 10 cells were retained, and 2,000 highly variable genes were selected without a batch key. Thirty principal components were integrated with Harmony (maximum 30 iterations), followed by a 15-neighbor graph and UMAP with seed 0. Cell annotations used fixed canonical marker panels listed in Table S3. Per-cell scores were computed with the Scanpy score_genes algorithm on log1p counts per 10,000 as the panel mean minus the mean of expression-matched control genes, using 25 expression bins, up to 50 control genes per bin and seed 0. Each Leiden cluster was assigned the panel with the highest mean score when this score was ≥0.3 and exceeded the next-highest score by ≥0.1; otherwise the cluster remained unresolved. Both epithelial panels defined epithelial identity, with the dominant panel specifying the hepatocyte-like or ductal-like subgroup. PROM1 was absent from all panels. A separate four-gene cell-cycle score was not used for assignment. Hepatocyte-like, ductal-like and unresolved epithelial candidates were retained for copy-number assessment so that candidate selection did not require preserved hepatocyte differentiation. Labels were compared with CellTypist Healthy_Human_Liver and Immune_All_High predictions and reviewed. A mixed low-quality cluster of 2,408 cells was removed, leaving 15,748 epithelial or unresolved candidates for copy-number assessment.^51,57^

CopyKAT 1.2.5 was run per patient with candidate cells and up to 1,000 T/NK reference cells, partitioned into 800 reference and 200 held-out calibration cells where available. Candidate calls comprised 5,399 aneuploid, 6,278 not-aneuploid and 4,071 unclassified cells. HCC07 failed reference calibration, with 68.2% of held-out cells called aneuploid, and was excluded from tumor-cell case analyses. HCC09 had no aneuploid candidates. Per-patient candidate, aneuploid and PROM1-detected counts and held-out-reference calibration are provided in Table S3. Aneuploid calls were used to enrich for malignant cells; a not-aneuploid call was not taken as proof of non-malignancy.^48^

PROM1 detection was defined as at least one UMI. Optimal one-to-one matching without replacement was performed within patient, with calipers of 0.20 on log1p UMI count and log1p detected-gene count. Matching assessed within-patient comparability because transcript detection can depend on sequencing depth and the number of detected genes. The maximum absolute standardized mean difference across these two covariates was recorded for the matched pairs (Table S3). Cases were considered eligible for detailed within-case analysis when at least 100 matched pairs were available. Only HCC05 met this threshold, with 272 pairs and a maximum absolute standardized mean difference of 0.15; the next-largest number was 57 pairs in HCC06. Figures 4F, 4G and S6 show all 866 HCC05 aneuploid cells, including 301 PROM1-detected and 565 PROM1-undetected cells, rather than only the matched subset.

The reciprocal-response programs were scored as the mean of gene-wise z-scores of log1p counts per 10,000, using the same 34 up-with-PROM1 and 73 down-with-PROM1 genes as the PLC/PRF/5 analysis. Scaling was within the HCC05 aneuploid population for the case panels and across candidate cells for the cohort overview; overview component scores were additionally standardized for display. For the cross-dataset comparison in Figure S6, the 145 genes with |Spearman ρ| > 0.3 along the PLC axis were displayed across 40 quantile bins of each dataset’s ordering. PLC/PRF/5 and HuH-7 used their diffusion coordinates. HCC05 used an up-minus-down score formed from the 85 reciprocal-signature genes absent from the displayed axis-gene set (27 up-with-PROM1 and 58 down-with-PROM1 genes). Expression profiles were shown as gene-wise z-scores across bins, with rows ordered by the HuH-7 gradient. PROM1 detection fractions and expression were displayed above the profiles. Directional agreement with PLC was summarized descriptively; the HCC05 ordering was score-based and did not use any of the 145 displayed genes. Table S3 provides the gene-level correlations, binned summaries and HCC05 score statistics.

#### Endogenous ALFA tagging

ALFA-tag sequences were introduced into the endogenous PROM1 locus in PLC/PRF/5 cells by CRISPR/Cas9-mediated editing using single-stranded donor oligonucleotides. The ALFA-tag/nanobody epitope system was described by Götzke et al.^18^ Two insertion sites, located in an extracellular region and the first cytoplasmic loop of CD133, were selected with guidance from AlphaFold structural predictions to minimize potential disruption of protein structure.^55^ Cells were cotransfected with a plasmid encoding Cas9 and the corresponding guide RNA (sgRNA1 for the extracellular site or sgRNA4 for the intracellular site), together with 5 μL of donor oligonucleotide stock at 4 pmol/μL (20 pmol). Donors contained the ALFA-tag coding sequence flanked by sequences homologous to the target locus. Transfected bulk populations were selected with 5 μg/mL puromycin and screened before clonal isolation. Guide target sequences and ordered single-stranded donor oligonucleotide (ssODN) sequences are listed in Table S6. The sgRNA1 donor placed the insertion two nucleotides downstream of the Cas9 cleavage site, and the sgRNA4 donor placed it one nucleotide downstream, to preserve the reading frame. The sgRNA4 donor design also changed two linker glycine residues to serine to disrupt the PAM sequence.

Bulk populations were assessed by ALFA/CD133 immunofluorescence and PCR. Outside/outside PCR, using primers flanking the insertion site, yielded expected product sizes of 557 bp and 632 bp for untagged and tagged alleles, respectively, at the sgRNA1 site, and 925 bp and 1,000 bp at the sgRNA4 site. An additional inside/outside PCR, pairing an ALFA-internal primer with a PROM1-flanking primer, was used for genotyping at the sgRNA4 site. Primer sequences are provided in Table S6.

Clones were isolated from the edited bulk populations by limiting dilution in 96-well plates. All wells except A1 were initially filled with 100 μL of culture medium, and 200 μL of a cell suspension containing 2 × 10^4^ cells/mL was added to A1. Serial twofold dilutions were prepared down the first column by transferring 100 μL between successive wells, with gentle mixing after each transfer. After discarding 100 μL from H1, 100 μL of fresh medium was added to each well of the first column, and serial twofold dilutions were continued across the remaining columns. Final well volumes were adjusted to 200 μL. Plates were incubated at 37 °C in a humidified CO₂ incubator. Colonies were identified microscopically after 4–5 days, and wells were monitored repeatedly to select those containing a single visible colony.

Expanded clones were screened by PCR and ALFA-mediated immunoprecipitation. Selected PCR bands were excised, purified, cloned into pcDNA3.1(+), and sequenced to verify the insertion and examine the amplified PROM1 coding sequence for unintended changes. One clone per insertion site was selected: clone 1C4 carried the extracellular insertion, and clone 4B4 carried the insertion in the first cytoplasmic loop. No additional mutations were detected within the sequenced coding regions of the selected clones.

#### Affinity purification and immunoblotting

Affinity purification was performed using the manufacturer’s analytical-scale protocol for magnetic ALFA Selector Resins (NanoTag Biotechnologies, Göttingen, Germany), with modifications to the preclearing, washing and elution steps as described below. Cell pellets were resuspended in 400 μL of 1× Cell Lysis Buffer (Cell Signaling Technology; catalog no. 9803), homogenized on ice using a Dounce homogenizer (100 gentle strokes), and then clarified at 14,000 × g for 10 min at 4 °C. An input aliquot of approximately 20 μL was retained. Selector Control magnetic agarose beads (NanoTag Biotechnologies; catalog no. N0010) were equilibrated in 1× CST lysis buffer, and 20 μL of control beads was added to each clarified lysate for preclearing for 1 h at 4°C with end-over-end rotation. Control beads were removed using a magnetic rack, and the precleared supernatant was retained for affinity capture. For each purification, 20 μL of ALFA Selector ST magnetic resin slurry (10 μL packed beads; NanoTag Biotechnologies, cat. no. N1516) was equilibrated twice with 1 mL lysis buffer, with magnetic separation after each equilibration. Precleared lysates were incubated with the equilibrated resin for 1 h at 4°C with end-over-end rotation. Beads were collected using a magnetic rack and the unbound fraction was removed. Beads were resuspended in 1 mL lysis buffer and collected by magnetic separation. This wash was repeated three times, giving four washes in total. A final wash was performed with 1 mL Tris-buffered saline (TBS), and the supernatant was removed completely after magnetic separation. For immunoblotting, bound proteins were eluted in loading buffer for 10 min at 70°C, as recorded for this experiment. Bead-bound material for mass spectrometry was processed separately as described below. Four samples per group were compared from untagged PROM1-activated cells, clone 1C4 and clone 4B4.

At 1× concentration, the manufacturer’s formulation of Cell Lysis Buffer no. 9803 contains 20 mM Tris-HCl (pH 7.5), 150 mM NaCl, 1 mM Na₂EDTA, 1 mM EGTA, 1% Triton, 2.5 mM sodium pyrophosphate, 1 mM β-glycerophosphate, 1 mM sodium orthovanadate and 1 μg/mL leupeptin.

General whole-cell immunoblotting used T-PER tissue protein extraction reagent (Thermo Fisher Scientific; catalog no. 78510) and Bradford protein quantification. The recorded protocol loaded 5 μg protein per lane in a 15 μL preparation containing 1.5 μL reducing agent and 3.75 μL LDS sample buffer. Samples were heated at 70 °C for 10 min and separated on Bolt 4–12% Bis-Tris gels (Thermo Fisher Scientific) at 130 V for 60–90 min. Proteins were transferred using an iBlot 2 Gel Transfer Device (Thermo Fisher Scientific) with sequential steps of 20 V for 1 min, 23 V for 4 min and 25 V for 2 min. Membranes were assessed by Ponceau staining, blocked in Bio-Rad blocking buffer for 20 min at room temperature and incubated with primary antibodies overnight at 4 °C. After three 10 min washes in TBS containing 0.1% Tween-20 (TBST), rabbit primary antibodies were detected with HRP-conjugated goat anti-rabbit IgG (Abbkine Scientific; catalog no. A21020; RRID:AB_2876889) at 1:20,000 for 1 h at room temperature, followed by three further 10 min TBST washes. Chemiluminescence was developed using Clarity Max Western ECL reagents (Bio-Rad; catalog no. 1705062) and acquired with a ChemiDoc MP system (Bio-Rad). CD133 and ALFA reagents are listed in the key resources table. The recorded HRP-conjugated β-actin reagent was the mouse monoclonal antibody mAbcam 8226 (Abcam; catalog no. ab20272). Figure 5F summarizes immunoblot IP/input ratios for TUBB6, ATP6V1A, DUSP9 and UMPS, expressed relative to the corresponding mean ratio in the untagged PROM1-activated control (WTOE). Adjusted total band volumes were used to calculate matched IP/input ratios, normalized to the arithmetic mean of the four WTOE ratios for each protein. Bars show arithmetic means, error bars show SD, and points show the four IP measurements per group.

Two-sided unpaired Student t tests were performed on log2-transformed normalized ratios, with Holm–Šidák adjustment across the extracellular-tag-versus-WTOE and intracellular-tag-versus-WTOE comparisons separately for each protein. CD133 was displayed as the bait immunoblot and was not included in the ratio tests. The two tagged groups each represent one edited clone, so the four measurements do not constitute four independent clones.

#### Mass spectrometry

CD133-associated proteins were investigated by affinity purification–mass spectrometry in PLC/PRF/5 cells with CRISPRa-mediated activation of endogenous PROM1. Three conditions were analyzed: untagged cells, cells carrying an extracellular ALFA tag in endogenous CD133 (clone 1C4), and cells carrying an intracellular ALFA tag (clone 4B4). Four immunoprecipitation samples were analyzed per condition, giving 12 samples in total. Untagged cells underwent the same affinity-purification procedure and served as controls for nonspecific recovery. Affinity purification, including 1 h preclearing with magnetic control beads, was performed using ALFA Selector ST magnetic resin as described above.

Bead-bound material was submitted to the Proteomics Platform at the Institute of Surgical Pathology, University Medical Center Freiburg. Proteins were eluted in 5% SDS and 50 mM triethylammonium bicarbonate (TEAB), pH 7.5, at 95°C for 10 min. Samples were reduced, alkylated and digested using an S-Trap spin-column workflow (ProtiFi). Peptides were analyzed using a timsTOF mass spectrometer operated in data-dependent acquisition with parallel accumulation–serial fragmentation (DDA-PASEF), using the facility’s 30-SPD method and a 15-cm Performance column (EV1137).

#### Protein identification and quantification

Raw mass-spectrometry data were processed using MaxQuant version 2.1.4.0. Database searches specified trypsin digestion with up to two missed cleavages, cysteine carbamidomethylation as a fixed modification and no variable modifications. Match between runs was enabled, and at least one unique peptide was required for quantification. Protein abundance was assessed using MaxQuant label-free quantification (LFQ) intensities.^52,53^

The exported proteinGroups table contained 1,519 protein groups. The facility-provided statistical input contained exactly the 907 groups with at least one positive LFQ intensity among the 12 samples. Positive LFQ intensities were transformed to log2 values and zero LFQ intensities were represented as missing values; no imputed intensities were present in the supplied input matrix. Differential enrichment was assessed using limma, with pairwise comparisons between each tagged clone and the untagged control, a comparison between tagged clones, and an overall F-test. Benjamini–Hochberg adjustment was applied separately to the finite p values within each contrast and to the overall F-test. The supplied outputs contained 758 estimable control-versus-1C4 comparisons, 774 control-versus-4B4 comparisons and 709 finite overall F-tests; these adjustment families were verified from the exported unadjusted and adjusted p values.^42^

For presentation, clone-versus-control fold changes were oriented so that positive values indicated enrichment in the tagged samples. Quantitative candidates were selected by positive log₂ fold changes and adjusted P values <0.05 in both tagged-clone-versus-control comparisons, together with an adjusted F-test P value <0.05. Protein groups annotated as contaminants were excluded. This selection yielded 19 candidates; no additional minimum fold-change threshold was applied. Table S4 reports each retained protein-group identifier, the number of nonmissing LFQ observations per condition, both clone-versus-control log2 fold changes, contrast-specific adjusted p values and the adjusted overall F-test p value. Displayed linear fold enrichments were calculated as 2 raised to the clone-versus-control log2 fold change. Selection used unrounded values; no zero or pseudocount was substituted for an entirely unquantified control group.

#### Detection-based proteomic screening

A separate detection-based analysis was performed using the MaxQuant proteinGroups “Intensity” values rather than LFQ intensities. Detection was defined as an intensity greater than zero. After exclusion of protein groups containing reverse-sequence or contaminant identifiers, groups were retained if they were undetected in all four untagged-control samples and detected in all four samples from at least one tagged clone. This criterion yielded 15 protein groups, including CD133, displayed in Figure 5D and Table S4. Protein groups were retained as exported, including semicolon-separated accessions; representative gene labels were not treated as proof of isoform-specific identification. Table S4 distinguishes positive raw-intensity counts, direct MS/MS identification counts and positive LFQ counts for each condition; unique-peptide ranges were taken only from samples with direct MS/MS identification. Recovered proteins were considered candidate CD133-associated proteins. The detection-based and quantitative screens used different eligibility criteria and had no overlapping protein groups.

#### Immunofluorescence and microscopy

For fixed-cell immunofluorescence, PLC/PRF/5 cells were cultured in ibidi μ-Slide eight-well chambers, washed three times with PBS and fixed with 4% paraformaldehyde for 20 min at room temperature. After three PBS washes, nonspecific binding was blocked with 5% BSA in PBS for 1 h at room temperature in a humidified chamber. Primary antibodies were diluted 1:100 in PBS containing 3% BSA and 0.3% Triton X-100 and incubated overnight at 4 °C. After three PBS washes, fluorescent secondary antibodies were applied at 1:300 in the same BSA/Triton buffer for 1 h at room temperature. Where nuclear counterstaining was performed, Hoechst 33342 was applied at 1:2,000 in PBS for 5 min. Cells were washed three times with PBS and kept in PBS until imaging.

For live-cell surface immunofluorescence of ALFA-tagged CD133, unfixed cells were incubated with anti-CD133 and anti-ALFA primary antibodies, each diluted 1:100 in fresh DMEM, for 30 min at 37 °C. Cells were washed twice with culture medium and incubated with the appropriate fluorescent secondary antibodies at 1:300 in DMEM for 15 min at room temperature. After two further medium washes, fresh medium was added and cells were imaged immediately without fixation or detergent permeabilization. Phenol-red-free DMEM (Gibco; catalog no. 21063-029) was used for fluorescence imaging. This primary-plus-secondary procedure is indirect immunofluorescence on living cells.

Fluorescence reagents included PE-conjugated mouse anti-human CD133 (clone TMP4; eBioscience/Invitrogen; catalog no. 12-1338-42), rabbit anti-CD133 (clone D4W4N; Cell Signaling Technology; catalog no. 86781), rabbit anti-CD44 (clone E7K2Y; Cell Signaling Technology; catalog no. 37259), and recombinant anti-ALFA antibodies with rabbit IgG Fc or mouse IgG1 Fc domains (NanoTag Biotechnologies; catalog nos. N1583 and N1582, respectively). Highly cross-adsorbed donkey secondaries against rabbit or mouse IgG (H+L) included Alexa Fluor 488 conjugates (Invitrogen; A-21206 and A-21202, respectively) and Alexa Fluor 594 conjugates (A-21207 and A-21203, respectively).

Fluorescence microscopy was performed using an Olympus IX73 inverted microscope equipped with a DP74 color CMOS camera, a CoolLED pE-300white LED illumination source and cellSens Dimension acquisition software (Olympus). The configured objectives were UPLFLN10X2PH-2 (10×/0.30 NA), LUCPLFLN20XPH (20×/0.45 NA), LUCPLFLN40XPH (40×/0.60 NA) and UPLXAPO60XOPH (60×/1.42 NA, oil immersion). The system included DAPI filters with excitation/emission passbands of 349–389/416–480 nm, GFP/FITC filters with passbands of 459–487/500–540 nm, and Texas Red filters with passbands of 559–591/609–641 nm, together with a FITC/Texas Red dual-band filter.

Images were processed and analyzed using Fiji (ImageJ).^54^ Channels were assigned display colors as indicated in each figure. Localization was assessed qualitatively.

The supporting images in Figure S2A show PROM1-CRISPRa cells co-cultured with GFP-reference PLC/PRF/5 cells. Individual-channel and merged images are shown at two field scales; the higher-magnification field is not a crop of the overview. Figure 3E shows a CD44-stained overview of a PROM1-activated culture and two indicated regions. For figure assembly, white separator gutters were removed and the channel tiles rearranged without changing the supplied image pixel intensities. Original scale bars were retained: 100 µm and 20 µm in the two Figure S2A co-culture fields, and 500 µm in the Figure 3E overview with 100 µm insets. The CD44 panel uses an intensity pseudocolor scale from blue to green. These images were assessed descriptively without image-based quantification.

#### CAM assay and trametinib exposure

Fertilized specific-pathogen-free chicken eggs (VALO BioMedia, Osterholz-Scharmbeck, Germany) were used for the CAM assay. Two independent experimental repeats were analyzed. The recorded schedule defined incubation start as embryonic day (ED) 0, followed by shell puncture on ED8, shell opening and cell–Matrigel graft preparation on ED9, implantation on ED10 and harvest on ED15, 5 days after implantation. Eggs were maintained at 37 °C in a humidified incubator without supplemental CO2. The procedure was carried out in accordance with the description in Gäbele et al.^58^ 5 × 10^6^ cells per graft were suspended in 20 μL culture medium and mixed with 20 μL Matrigel (Corning, catalog no. 356237) on ice. Grafts were allowed to set for 10 min at 37 °C, the dish was inverted for a further 20 min, and grafts were covered with culture medium and precultured overnight. One graft was placed onto the CAM within a silicone ring, covered with 20 μL Matrigel and incubated until harvest; egg windows were sealed with Durapore tape (3M). Stable PROM1-CRISPRa and guide-free SAMph control cells were studied with and without trametinib. Each egg carried one graft and was separately allocated to a cell-population and treatment group within its repeat; treatment or control exposure was administered individually. Trametinib (Selleck Chemicals; catalog no. S2673) was applied at 50 nM topically to the CAM/graft through the egg window during incubation.

At harvest, grafts were excised with the surrounding CAM and measured in three perpendicular dimensions. Ellipsoid volumes were calculated from the recorded length, width and height as V = (π/6) × length × width × height, with dimensions in millimeters and volume in cubic millimeters. Calculations used the unrounded dimensions. Repeat 1 included Ca2–Ca6 (CRISPRa; n = 5), CaT1, CaT3, CaT4 and CaT8 (CRISPRa plus trametinib; n = 4), Sa1, Sa2 and Sa4–Sa7 (SAMph; n = 6), and SaT1–SaT6 (SAMph plus trametinib; n = 6). Repeat 2 included Ca20–Ca28 (CRISPRa; n = 9), CaT20–CaT25 (CRISPRa plus trametinib; n = 6), Sa20–Sa24 (SAMph; n = 5), and SaT20–SaT24 (SAMph plus trametinib; n = 5). All 46 grafts with complete dimensions in these blocks were included.

#### CAM histology and immunohistochemistry

Excised CAM grafts were fixed in 4% paraformaldehyde overnight to 24 h at room temperature according to the local SOP, processed and embedded in paraffin. Hematoxylin and eosin staining was used to assess tumor morphology, tissue viability and evidence of infection. Immunohistochemistry examined CD133, CD44, Ki-67 (MIB1), and cleaved caspase-3. The local IHC SOP includes 2 µm paraffin sections on Superfrost slides, heating at 70 °C for at least 30 min for fresh sections, deparaffinization in xylene (three 15-min incubations), and rehydration through 100–70% alcohol. Heat-induced antigen retrieval is performed in citrate buffer (pH 6.0) or Tris–EDTA (pH 9.0; Vector) for 8–12 min at 110 °C, followed by cooling. The protocol specifies Dako S3006 wash buffer, peroxidase suppression for 10–15 min, and overnight primary-antibody incubation at room temperature in Dako Antibody Diluent S2022 in a humidified chamber. Detection uses Cell Signaling anti-rabbit/mouse polymer for 60 min at room temperature and ImmPACT NovaRED chromogen (Vector Laboratories; catalog no. SK-4805), followed by Mayer’s hemalum counterstaining. The illustrated PROM1-CRISPRa (Ca) and SAMph (Sa) CAM graft sections were assessed by qualitative visual inspection of H&E, CD133, CD44, cleaved-caspase-3 and Ki-67 (MIB1) images.

#### Colony formation

For colony formation, cells were plated at low density and colonies were documented and counted after 10 days in one experiment with duplicate wells per population. The assay follows the general colony-outgrowth format used previously, in which low-density cultures in complete medium were fixed, stained with crystal violet and counted with ImageJ.^59^

#### Sphere formation

Sphere formation was assessed after 5 days in two experiments, with five wells per condition in each experiment. The first experiment used 0% methylcellulose and the second used 2% methylcellulose. Phase-contrast images were acquired at 20× objective magnification; readouts included the mean sphere number per microscopic field and sphere area relative to the control population. The source summaries are displayed descriptively, and the two medium formulations are identified separately in the image panel. The procedure was carried out as published previously: a framework of nonadherent culture in serum-free DMEM/F12 containing B27 without vitamin A (Life Technologies), EGF and bFGF (R&D Systems), both at 20 ng/mL.^60^

#### Conditioned medium and monocyte migration

Conditioned medium from SAMph and PROM1-CRISPRa cultures was used to test effects on monocyte migration and endothelial network formation. Monocyte migration was assessed at 6 h in one experiment, including negative and positive controls and conditioned medium from each population. Migrated cells were visualized at 20× and reported as the mean number per field. This design is consistent with the conditioned-medium chemotaxis approach used previously by Raggi and colleagues.^61^

#### Endothelial network formation

Human umbilical vein endothelial cells (HUVECs) were exposed to conditioned medium from SAMph or PROM1-CRISPRa cultures in one experiment. Images were recorded after 3, 6 and 24 h; network quantification was available at 6 and 24 h. Basal endothelial medium, labeled 0% EBM in the source data, served as an additional comparator. Readouts were the numbers of master segments, master junctions, meshes, branches and segments. Meshes were defined as enclosed network areas, segments as elements joining junctions, and branches as elements between a junction and a terminal extremity. Related work from the Raggi laboratory uses a Matrigel-based HUVEC tube-formation assay to evaluate tumor-conditioned medium.^61^

### QUANTIFICATION AND STATISTICAL ANALYSIS

Acute bulk sequencing comprised three independently transfected wells per group within one experiment for each arm, with a separately prepared transfection mixture for each well and separate controls for activation and knockdown. Activation qPCR comprised six transfected wells per group across two independent transfections (three wells per experiment); knockdown qPCR comprised three wells per group. Technical qPCR triplicates were averaged within each well. Immunoblot validation comprised three separately transfected wells per group in one experiment per arm. Stable single-cell profiling comprised two measured pools per condition, each formed from two cultures. Cell-state fractions and projection-bin occupancies are reported descriptively with pool-level values; individual cells were not treated as independent treatment replicates.^62^

The acute flow-cytometry experiment comprised one fully stained tube per condition, measured 48 h after transient transfection without GFP-reference cells. Marker fractions and analysis-setting sensitivities are descriptive. Event counts, staining controls and reanalysis variants do not constitute biological replicates; no inferential between-condition test was performed.

For Figure S1D–G, bars show arithmetic means, error bars SD and points individual transfected wells. Validation testing used unpaired, two-sided Student t tests on log2-transformed relative values. Holm–Šidák correction was applied separately across seven transcripts in each qPCR panel and across four proteins in the activation immunoblot panel; the single-protein knockdown test was unadjusted. Significance labels indicate p < 0.05, p < 0.01, p < 0.001 and p < 0.0001. These are exploratory well-level tests. The activation qPCR pooled wells from two experiments without an experiment term, and the immunoblots represent one experiment per perturbation arm; their p values therefore do not establish experiment-level reproducibility.

Gene-level adjusted p values refer to the multiple-testing procedure applied to each contrast; pathway FDRs refer to the enrichment analysis. The reciprocal-response set used adjusted p < 0.01 in both acute contrasts, followed by a sign comparison. The 94.4% reciprocal fraction is conditional on significance in both arms. The PLC/PRF/5 reciprocal-response score used the 117 reciprocal genes followed by assay restriction and PROM1 exclusion, yielding 107 genes (34 activation-up/silencing-down and 73 with the inverse response). The human-tumor score panels used this same 107-gene reciprocal signature, with gene-wise standardization within the relevant human-cell population. For the cross-dataset heatmap in Figure S6, the HCC05 ordering used the 85 signature genes outside the 145 displayed axis genes. Figure S3 instead used all measurable genes meeting adjusted p < 0.01 in both arms or in each arm separately, without a fold-change cutoff. Its Spearman correlations across selected genes are descriptive and do not quantify genome-wide concordance or an independent treatment effect. Figure S2 state fractions, robustness measures and score correlations likewise do not use individual cells as independent treatment replicates.

The HuH-7 reanalysis comprised one public single-cell library, with cells serving as analytical units. Its p values describe variation within that culture and do not estimate between-culture reproducibility. Pathway and regulon tests used the analysis-specific false-discovery-rate corrections described above; axis and cross-model correlations are interpreted as exploratory associations. Gene-level permutation and bootstrap procedures assume exchangeability and can understate uncertainty when genes are correlated. Decile- or bin-level summaries were distinguished from individual-cell associations.

For affinity-purification proteomics, four IP samples per condition were analyzed using the facility-provided limma models of log2 LFQ intensity. Quantitative selection required enrichment in both tagged clones, Benjamini–Hochberg-adjusted p < 0.05 in each clone-versus-control contrast, and an adjusted overall F-test p < 0.05. Nonmissing LFQ counts and all selection statistics are provided in Table S4. The raw-intensity detection screen was descriptive and did not assign p values. These proteomics tests are separate from the immunoblot IP/input comparisons in Figure 5F.

For the CAM model, the egg was the experimental unit because each egg carried one graft and was separately allocated and treated within its repeat. All 46 grafts with recorded dimensions were analyzed; the unmeasured Ca29 entry was excluded without assigning a zero volume. Positive graft volumes were natural-log transformed to reduce right skew and express effects on a proportional scale. Ordinary least-squares regression included repeat as a fixed blocking factor, cell population (SAMph or PROM1-CRISPRa), trametinib exposure, and the population-by-treatment interaction. Repeat was treated as fixed because the analysis concerned the two observed runs. Heteroskedasticity-consistent HC3 covariance estimates were used to accommodate unequal residual variances and unequal group sizes. Contrast estimates were divided by their robust standard errors, and two-sided p values and 95% confidence intervals were calculated using an approximate Student t reference distribution with 41 residual degrees of freedom (46 eggs minus five fitted coefficients).

The exploratory CAM analysis evaluated five contrasts: CRISPRa versus SAMph without trametinib; CRISPRa versus SAMph with trametinib; trametinib versus no trametinib within SAMph; trametinib versus no trametinib within CRISPRa; and the population-by-treatment interaction. Holm correction controlled family-wise error across all five tests, with adjusted p < 0.05 used as the significance criterion. Exponentiated log contrasts and confidence limits are reported as adjusted geometric-mean ratios; the interaction is the CRISPRa treated/untreated ratio divided by the SAMph treated/untreated ratio. The reported 95% confidence intervals are pointwise and were not adjusted for multiplicity. As a sensitivity analysis, a fully interacted repeat-by-population-by-treatment model allowed a separate mean for each of the eight repeat– population–treatment combinations. The corresponding log contrasts were averaged equally across the two repeats, with HC3 covariance, 38 residual degrees of freedom and a separate Holm correction across the same five tests. The primary model assumes common population and treatment contrasts across repeats, whereas this sensitivity analysis permits them to vary.

