## Supplemental Information for "Endogenous PROM1 activation redistributes liver cancer cells along a preexisting differentiation axis"

Document S1. Figures S1–S6, Tables S5 and S6, and supplemental item legends.

#### Contents

Figure S1 — PROM1 locus and acute perturbation validation

Figure S2 — Single-cell quality control and robustness

Figure S3 — Reciprocal response genes along the control axis

Figure S4 — In vitro functional assays

Figure S5 — CAM graft volume and histology

Figure S6 — State-axis genes across PLC/PRF/5, HuH-7 and HCC05

Table S5 — Custom EGFP probes

Table S6 — Endogenous ALFA tagging oligonucleotides

Separate supplemental files

Long data tables are supplied as separate Excel workbooks (Tables S1–S4). Uncropped immunoblots are supplied as Data S2. Their item descriptions are listed at the end of this document.

### Figure S1

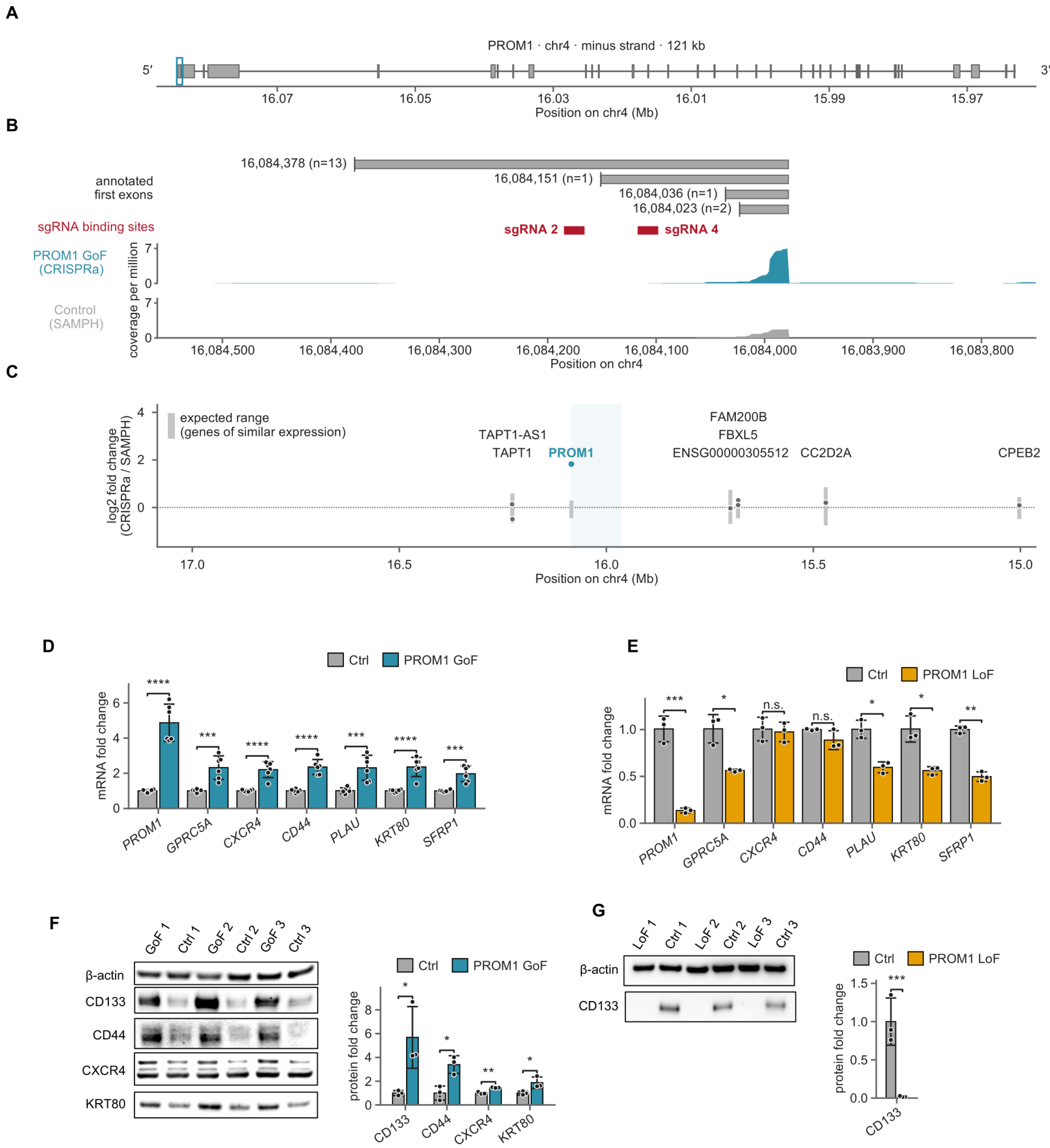
A
PROM1 · chr4 · minus strand · 121 kb
5′
3′
16.07
16.05
16.03
16.01
15.99
15.97
Position on chr4 (Mb)
B
16,084,378 (n=13)
16,084,151 (n=1)
16,084,036 (n=1)
16,084,023 (n=2)
annotated first exons
sgRNA binding sites
sgRNA 2
sgRNA 4
PROM1 GoF (CRISPRa)
Control (SAMPH)
coverage per million
16,084,500
16,084,400
16,084,300
16,084,200
16,084,100
16,084,000
16,083,900
16,083,800
Position on chr4
C
expected range (genes of similar expression)
TAPT1-AS1
TAPT1
PROM1
FAM200B
FBXL5
ENSG00000305512
CC2D2A
CPEB2
log2 fold change (CRISPRa / SAMPH)
17.0
16.5
16.0
15.5
15.0
Position on chr4 (Mb)
D
Ctrl
PROM1 GoF
mRNA fold change
PROM1
GPRC5A
CXCR4
CD44
PLAU
KRT80
SFRP1
E
Ctrl
PROM1 LoF
mRNA fold change
n.s.
F
GoF 1
Ctrl 1
GoF 2
Ctrl 2
GoF 3
Ctrl 3
β-actin
CD133
CD44
CXCR4
KRT80
protein fold change
G
LoF 1
Ctrl 1
LoF 2
Ctrl 2
LoF 3
Ctrl 3
β-actin
CD133
Ctrl
PROM1 LoF
protein fold change

#### Figure S1. The PROM1 locus, specificity of CRISPR activation, and validation of PROM1 gain and loss of function. Related to Figure 1.

(A and B) Gene model of PROM1 on chromosome 4 (minus strand, 121 kb; GENCODE 50), oriented 5′→3′ (A), with the boxed 5′ region enlarged in (B). Upper track, annotated alternative first exons, each labeled with its transcription start site (TSS) and the number of transcripts initiating there. Red, target sites of the two promoter-directed sgRNAs, 164 and 96 bp upstream of the TSS predominantly used in PLC/PRF/5 cells. Lower tracks, RNA-seq coverage after activation (teal) and in control cells (gray) from the same run, pooled across the three libraries per condition, scaled to million assigned fragments and plotted on identical axes.

(C) Local specificity of activation. Dots, log2 fold change of every expressed gene within ±1 Mb of PROM1, placed at its TSS. Gray bars, the central 95% of log2 fold changes among genome-wide genes of comparable expression (1,194–1,870 matched genes per neighbor; descriptive, not a confidence interval). Light shading, PROM1 gene body. PROM1 is the only gene outside its expected range.

(D and E) TaqMan qPCR of PROM1 and six response genes after activation (D) and knockdown (E), normalized to GAPDH by the ΔΔCt method and expressed relative to the corresponding control mean.

(F and G) Immunoblots after activation (F) and knockdown (G) with their densitometry. CD133 is the protein encoded by PROM1; β-actin served as loading control. Lysates were loaded in alternating lanes; targets on separate membranes were normalized to β-actin of the parallel blot of the same lysates. For CXCR4, the two upper immunoreactive bands were quantified together. Band volumes were divided by β-actin and scaled to the control mean.

Cells were harvested 48 h after transfection; each well received its own transfection mix. (D) $n = 6$ independently transfected wells per condition, from two experiments of three wells each; the RNA of one experiment is the RNA sequenced in Figure 1. (E) $n = 3$ independently transfected wells per condition, one experiment; this is the RNA sequenced in Figure 1. (F and G) $n = 3$ independently transfected wells per condition, one experiment. Bars, mean; error bars, SD; dots, individual wells. Statistics (D–G): unpaired two-tailed Student's t test on log2-transformed relative values, Holm-Šidák-corrected across the genes (D and E, seven genes) or proteins (F, four proteins) of each panel; (G) single protein, uncorrected. *$p < 0.05$, **$p < 0.01$, ***$p < 0.001$, ****$p < 0.0001$; n.s., not significant. Source values for (B) and (D–G) are in Data S1; uncropped blots with lane assignment and exposure times in Data S2.

#### Figure S2

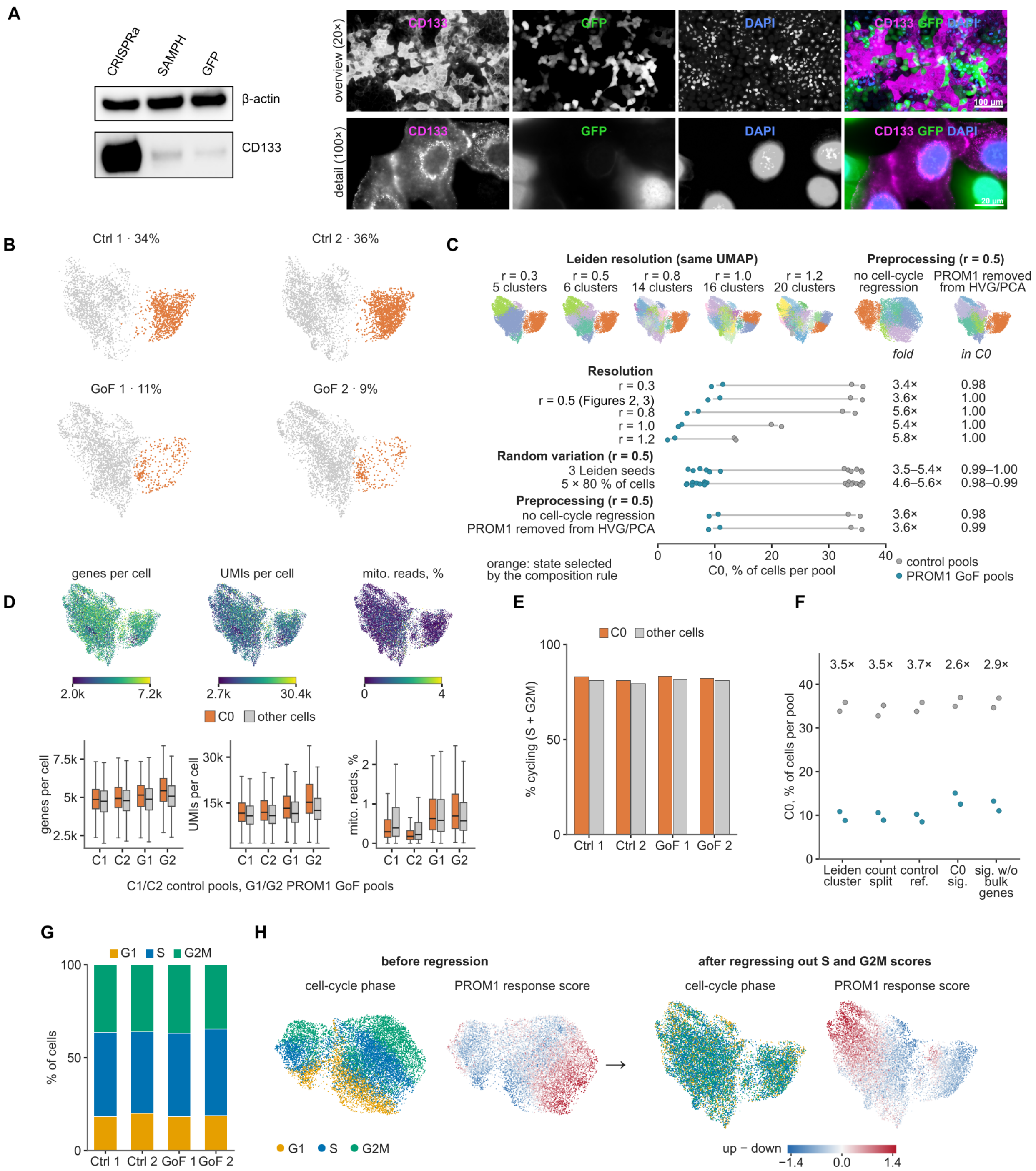
A
CRISPRa
SAMPH
GFP
β-actin
CD133
overview (20×)
detail (100×)
CD133
GFP
DAPI
CD133 GFP DAPI
100 µm
20 µm
B
Ctrl 1 · 34%
Ctrl 2 · 36%
GoF 1 · 11%
GoF 2 · 9%
C
Leiden resolution (same UMAP)
r = 0.3 5 clusters
r = 0.5 6 clusters
r = 0.8 14 clusters
r = 1.0 16 clusters
r = 1.2 20 clusters
Preprocessing (r = 0.5)
no cell-cycle regression
PROM1 removed from HVG/PCA
fold
in C0
Resolution
r = 0.3
r = 0.5 (Figures 2, 3)
r = 0.8
r = 1.0
r = 1.2
Random variation (r = 0.5)
3 Leiden seeds
5 × 80 % of cells
Preprocessing (r = 0.5)
no cell-cycle regression
PROM1 removed from HVG/PCA
3.4× 0.98
3.6× 1.00
5.6× 1.00
5.4× 1.00
5.8× 1.00
3.5–5.4× 0.99–1.00
4.6–5.6× 0.98–0.99
3.6× 0.98
3.6× 0.99
orange: state selected by the composition rule
C0, % of cells per pool
control pools
PROM1 GoF pools
D
genes per cell
UMIs per cell
mito. reads, %
2.0k 7.2k
2.7k 30.4k
0 4
C0
other cells
C1 C2 G1 G2
C1/C2 control pools, G1/G2 PROM1 GoF pools
E
C0
other cells
% cycling (S + G2M)
Ctrl 1 Ctrl 2 GoF 1 GoF 2
F
3.5× 3.5× 3.7× 2.6× 2.9×
C0, % of cells per pool
Leiden cluster
count split
control ref.
C0 sig.
sig. w/o bulk genes
G
G1 S G2M
% of cells
Ctrl 1 Ctrl 2 GoF 1 GoF 2
H
before regression
after regressing out S and G2M scores
cell-cycle phase
PROM1 response score
G1 S G2M
up − down
−1.4 0.0 1.4

#### Figure S2. Quality control and robustness of the single-cell analysis. Related to Figures 2 and 3.

(A) Left, immunoblot of CD133 in stable CRISPRa, SAMPH control and GFP bystander cells; β-actin, loading control. Right, immunofluorescence of CD133 (magenta), GFP (green) and DAPI (blue) in the co-culture, as overview (20×) and detail (100×). Scale bars, 100 μm and 20 μm.

(B) C0 on the UMAP of each pool, with its share of cells.

(C) Robustness of the C0 depletion to clustering and preprocessing. Top, UMAPs at five Leiden resolutions and after two preprocessing changes (no cell-cycle regression; PROM1 removed from highly variable genes and PCA), with the resulting number of clusters; orange, the cluster selected by the composition rule. Bottom, C0 per pool for each variant, including three clustering seeds and five 80% subsamples, with the control/CRISPRa ratio and the overlap with the cluster used in the figures. Gray, control pools; teal, CRISPRa pools.

(D) Sequencing quality: genes per cell, UMIs per cell and mitochondrial reads on the UMAP, and per pool for C0 versus the remaining cells. Boxes, interquartile range with median; whiskers, 1.5× interquartile range; outliers not shown.

(E) Cycling cells (S or G2M phase) per pool, C0 versus the remaining cells.

(F) C0 per pool from five estimators: the Leiden clustering, an independently split half of the counts, assignment to a frozen control reference, a C0 signature score, and the same score after removing every gene changed by activation in the bulk experiment. Numbers, control/CRISPRa ratio.

(G) Cell-cycle phase composition per pool.

(H) Cell-cycle phase and PROM1 response score on the UMAP, before and after regressing out the S and G2M scores.

n = 2 pools per condition, each from two independently transduced cultures; (A) one experiment. Descriptive; no inferential test of condition. Values in Table S2.

### Figure S3

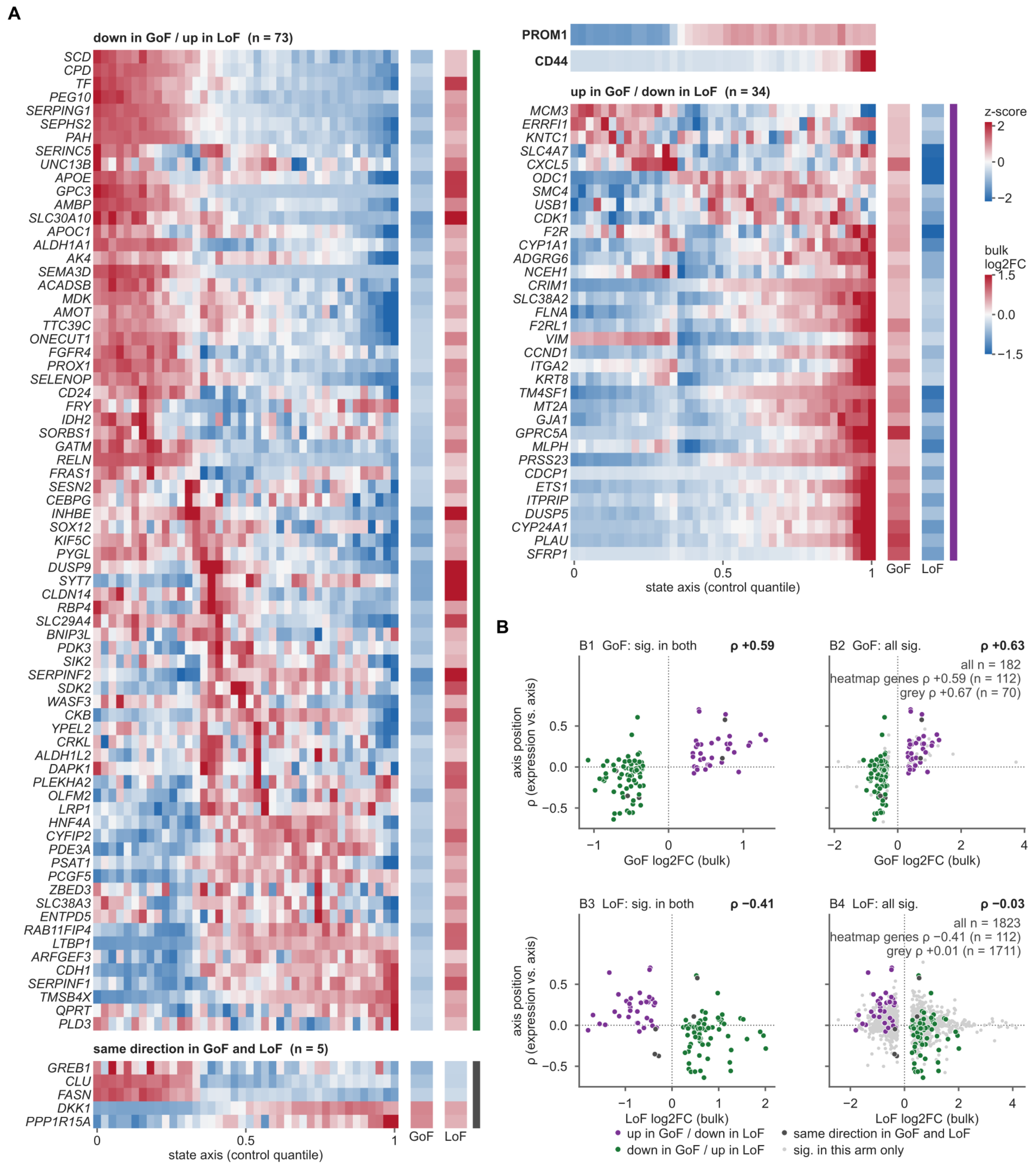
A
down in GoF / up in LoF (n = 73)
SCD
CPD
TF
PEG10
SERPING1
SEPHS2
PAH
SERINC5
UNC13B
APOE
GPC3
AMBP
SLC30A10
APOC1
ALDH1A1
AK4
SEMA3D
ACADSB
MDK
AMOT
TTC39C
ONECUT1
FGFR4
PROX1
SELENOP
CD24
FRY
IDH2
SORBS1
GATM
RELN
FRAS1
SESN2
CEBPG
INHBE
SOX12
KIF5C
PYGL
DUSP9
SYT7
CLDN14
RBP4
SLC29A4
BNIP3L
PDK3
SIK2
SERPINF2
SDK2
WASF3
CKB
YPEL2
CRKL
ALDH1L2
DAPK1
PLEKHA2
OLFM2
LRP1
HNF4A
CYFIP2
PDE3A
PSAT1
PCGF5
ZBED3
SLC38A3
ENTPD5
RAB11FIP4
LTBP1
ARFGEF3
CDH1
SERPINF1
TMSB4X
QPRT
PLD3
same direction in GoF and LoF (n = 5)
GREB1
CLU
FASN
DKK1
PPP1R15A
0
0.5
1
GoF
LoF
state axis (control quantile)
PROM1
CD44
up in GoF / down in LoF (n = 34)
MCM3
ERRFI1
KNTC1
SLC4A7
CXCL5
ODC1
SMC4
USB1
CDK1
F2R
CYP1A1
ADGRG6
NCEH1
CRIM1
SLC38A2
FLNA
F2RL1
VIM
CCND1
ITGA2
KRT8
TM4SF1
MT2A
GJA1
GPRC5A
MLPH
PRSS23
CDCP1
ETS1
ITPRIP
DUSP5
CYP24A1
PLAU
SFRP1
0
0.5
1
GoF
LoF
state axis (control quantile)
z-score
2
0
−2
bulk
log2FC
1.5
0.0
−1.5
B
B1 GoF: sig. in both
ρ +0.59
B2 GoF: all sig.
ρ +0.63
all n = 182
heatmap genes ρ +0.59 (n = 112)
grey ρ +0.67 (n = 70)
ρ (expression vs. axis)
axis position
0.5
0.0
−0.5
−1
0
1
GoF log2FC (bulk)
−2
0
2
4
GoF log2FC (bulk)
B3 LoF: sig. in both
ρ −0.41
B4 LoF: all sig.
ρ −0.03
all n = 1823
heatmap genes ρ −0.41 (n = 112)
grey ρ +0.01 (n = 1711)
ρ (expression vs. axis)
axis position
0.5
0.0
−0.5
−1
0
1
2
LoF log2FC (bulk)
−2
0
2
4
LoF log2FC (bulk)
up in GoF / down in LoF
down in GoF / up in LoF
same direction in GoF and LoF
sig. in this arm only

#### Figure S3. Genes regulated by both acute PROM1 perturbations are aligned with the control-derived state axis. Related to Figures 1 and 2.

(A) Expression along the state axis of the genes significantly changed in both bulk arms (CRISPRa versus SAMPH and PROM1 siPOOL versus control siPOOL; limma-voom, adjusted $p < 0.01$ in each arm, irrespective of direction). Of 124 such genes, the 113 present in the single-cell probe panel are shown. Left, genes decreased by gain and increased by loss of function (n = 73) and, below them, the five genes changed in the same direction in both arms; right, PROM1 and an additional CD44 reference track, followed by genes increased by gain and decreased by loss of function (n = 34). Columns, 40 equal-width bins of the control-quantile state axis (0, hepatocyte-like C0 end; see Figure 2), control cells only (n = 6,589). Values are mean log-normalized expression per bin and pool, averaged across the two pools and z-scored per gene; bins with fewer than 25 cells in a pool are masked. Rows are ordered by the bin of maximal expression within each block. Side columns, bulk log2 fold change in the gain-of-function (GoF) and loss-of-function (LoF) arm.

(B) Axis position of each gene versus its bulk response. Axis position is the Spearman correlation between a gene's expression and the axis coordinate across control cells, computed per pool and averaged. B1 and B3, the genes of (A) except PROM1 (n = 112) against the GoF and the LoF log2 fold change; purple, increased by GoF and decreased by LoF; green, the reverse; gray, same direction in both arms. B2 and B4, all genes with adjusted $p < 0.01$ in the GoF arm alone (n = 182) or in the LoF arm alone (n = 1,823), with the genes of (A) colored on top; $\rho$ above each plot, Spearman correlation across all plotted genes; insets in B2 and B4, $\rho$ for the colored and the light-gray genes separately.

Genes were preselected on the bulk results and are co-expressed, and the axis derives from the same control cells used to compute axis positions; all correlations are therefore descriptive and no p values are reported. Bulk arms, n = 3 independently transfected wells per condition; single-cell controls, n = 2 pools. Values in Table S1 (both_arms_FDR01) and Table S2 (axis_vs_acute_effects, reciprocal_genes_readouts).

#### Figure S4

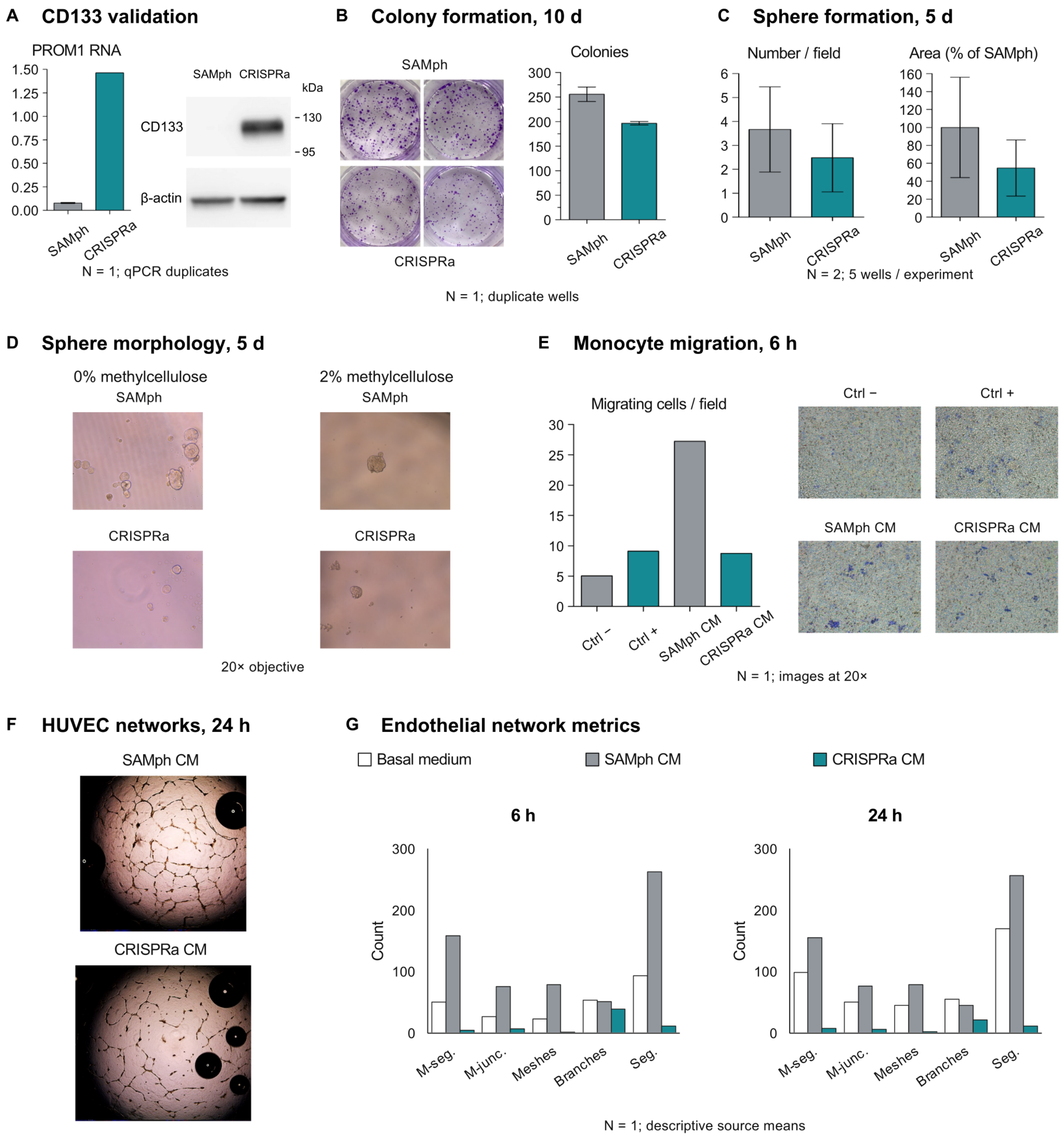
A CD133 validation
PROM1 RNA
1.50
1.25
1.00
0.75
0.50
0.25
0.00
SAMph
CRISPRa
N = 1; qPCR duplicates
SAMph CRISPRa
kDa
CD133
130
95
β-actin
B Colony formation, 10 d
SAMph
CRISPRa
Colonies
300
250
200
150
100
50
0
SAMph
CRISPRa
N = 1; duplicate wells
C Sphere formation, 5 d
Number / field
6
5
4
3
2
1
0
Area (% of SAMph)
160
140
120
100
80
60
40
20
0
SAMph
CRISPRa
SAMph
CRISPRa
N = 2; 5 wells / experiment
D Sphere morphology, 5 d
0% methylcellulose
SAMph
CRISPRa
2% methylcellulose
SAMph
CRISPRa
20× objective
E Monocyte migration, 6 h
Migrating cells / field
30
25
20
15
10
5
0
Ctrl −
Ctrl +
SAMph CM
CRISPRa CM
Ctrl −
Ctrl +
SAMph CM
CRISPRa CM
N = 1; images at 20×
F HUVEC networks, 24 h
SAMph CM
CRISPRa CM
G Endothelial network metrics
Basal medium
SAMph CM
CRISPRa CM
6 h
24 h
Count
300
200
100
0
M-seg.
M-junc.
Meshes
Branches
Seg.
N = 1; descriptive source means

#### Figure S4. Context-dependent effects of PROM1 activation in in vitro functional assays. Related to Figures 1 and 3.

(A) Validation of the cells used in (B–G) and Figure S5: PROM1 mRNA by qPCR (left) and CD133 immunoblot with β-actin as loading control (right) in PROM1-CRISPRa and SAMPH control cells.

(B) Colony formation after 10 d, colonies per well.

(C) Sphere formation after 5 d: spheres per field (left) and sphere area as percentage of the SAMPH mean (right).

(D) Sphere morphology after 5 d in 0% and 2% methylcellulose, 20× objective.

(E) Transwell migration of monocytes after 6 h towards conditioned medium (CM) of SAMPH or CRISPRa cells, with medium alone (Ctrl −) and a chemoattractant (Ctrl +) as references; migrating cells per field, with representative fields at 20×.

(F and G) HUVEC network formation in basal medium or in conditioned medium of SAMPH or CRISPRa cells: representative fields after 24 h (F) and network metrics after 6 h and 24 h (G; master segments, master junctions, meshes, branches and segments per field).

Panels (A–G) are exploratory experiments with the replicate numbers given in each panel (A, B, E and G, one experiment; C, two experiments with five wells each) and are shown descriptively without inferential testing. Available source values are in Data S1.

#### Figure S5

##### A CAM graft volume

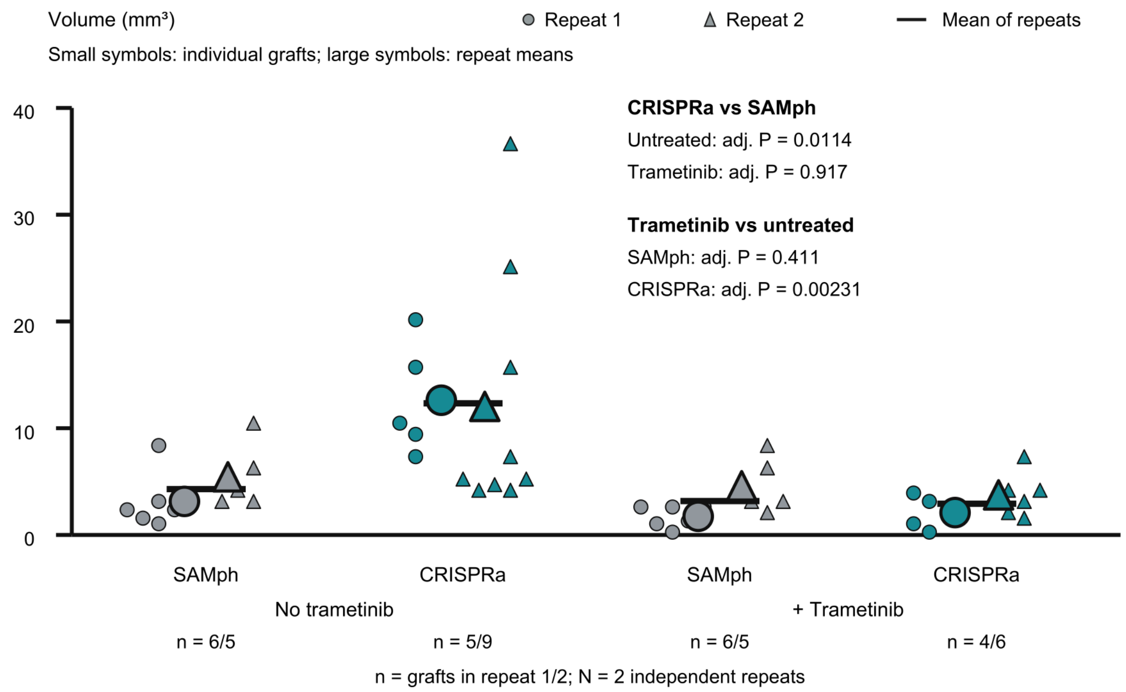


##### B CAM histology

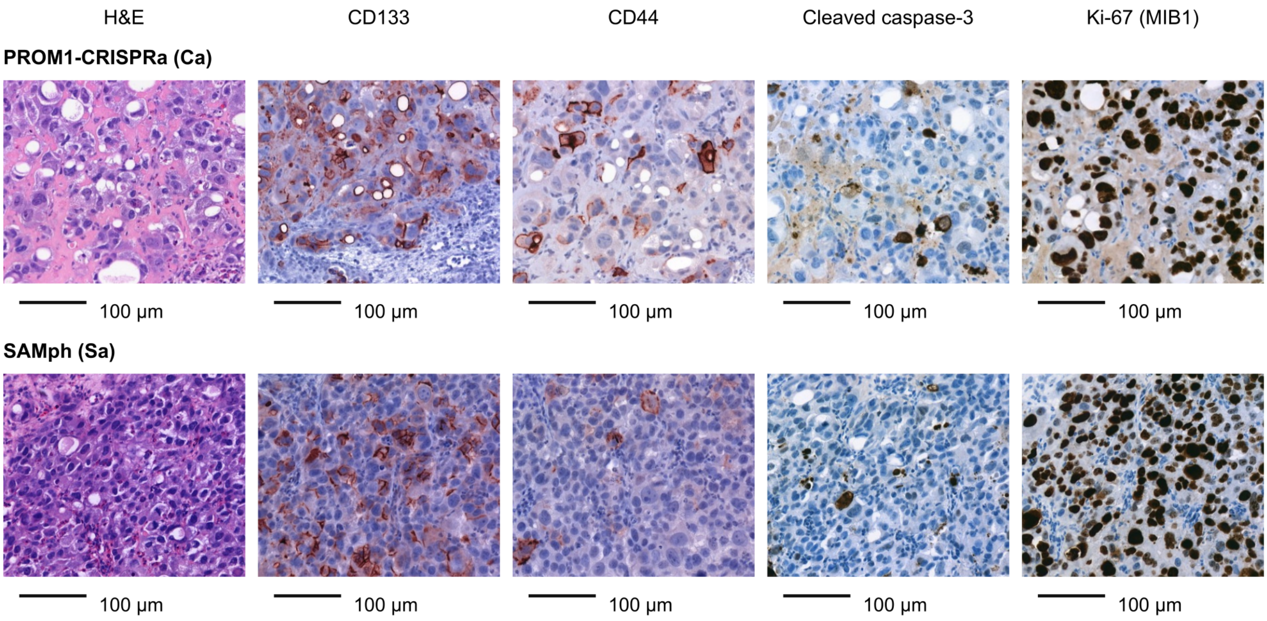

#### Figure S5. PROM1 activation and trametinib exposure in the chick chorioallantoic membrane model. Related to Figure 3.

(A) Volume of PROM1-CRISPRa and SAMPH grafts on the chick chorioallantoic membrane, without and with trametinib. Small symbols, individual grafts; large symbols, the mean of each repeat; n per group is given below the axis as grafts in repeat 1/repeat 2.

(B) Sections of CRISPRa (Ca) and SAMPH (Sa) grafts: H&E, CD133, CD44, cleaved caspase-3 and Ki-67 (MIB1). Scale bars, 100 μm.

(A) n = 2 independent repeats; adjusted p values are given in the panel. Each independently allocated and treated egg carried one graft. Primary inference used linear regression of log-transformed volume with cell population, trametinib, their interaction and repeat as fixed effects, HC3 heteroskedasticity-robust standard errors, and Holm correction across four simple contrasts and the population-by-treatment interaction. Circles and triangles distinguish repeats 1 and 2; large symbols show repeat-specific arithmetic means; black horizontal lines show the means of the two repeat means. Histology (B) was assessed visually without formal quantification. Source values and CAM statistics are in Data S1.

#### Figure S6

**A**

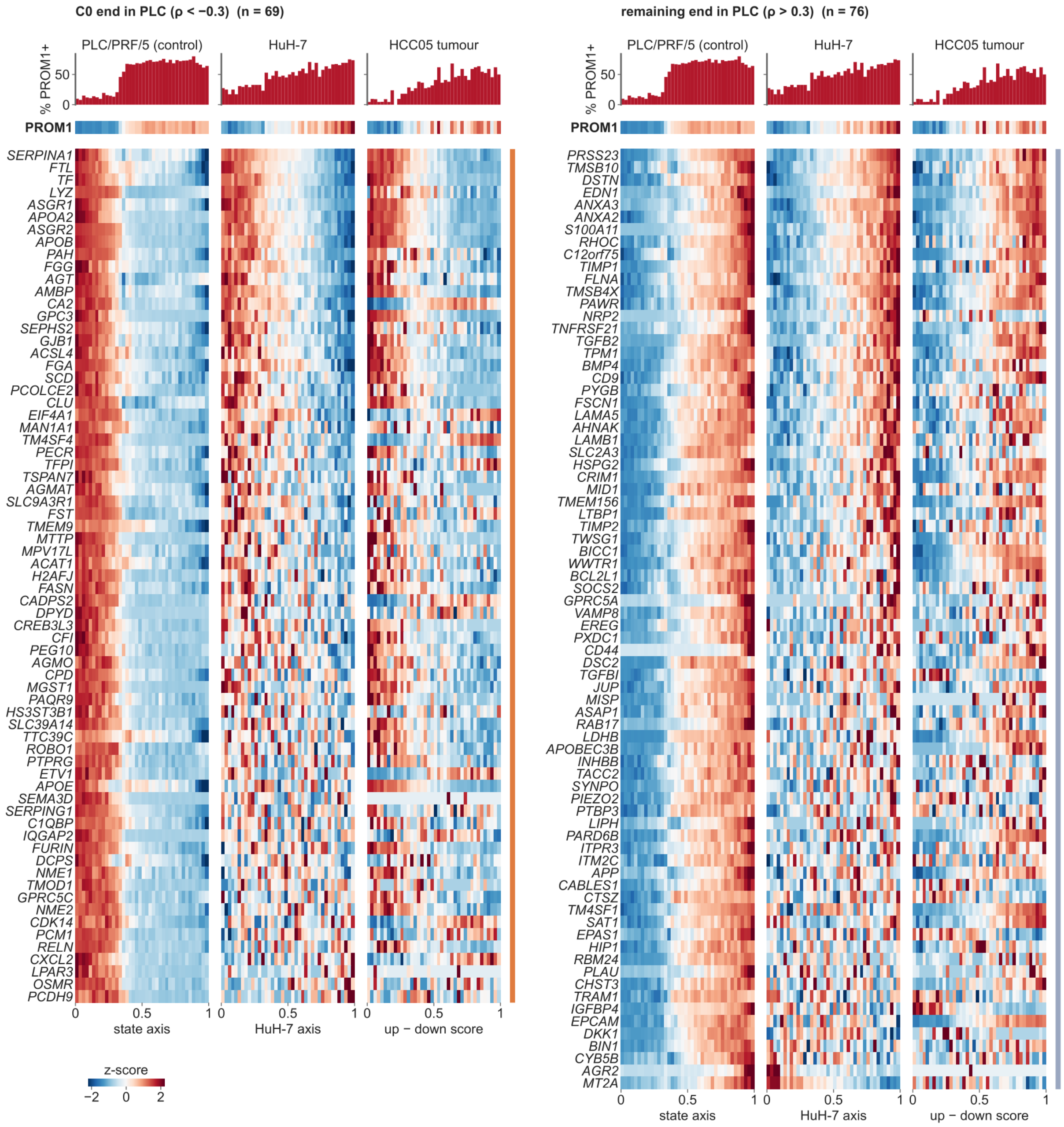
C0 end in PLC (ρ < −0.3) (n = 69)
remaining end in PLC (ρ > 0.3) (n = 76)
PLC/PRF/5 (control)
HuH-7
HCC05 tumour
% PROM1+
PROM1
SERPINA1
FTL
TF
LYZ
ASGR1
APOA2
ASGR2
APOB
PAH
FGG
AGT
AMBP
CA2
GPC3
SEPHS2
GJB1
ACSL4
FGA
SCD
PCOLCE2
CLU
EIF4A1
MAN1A1
TM4SF4
PECR
TFPI
TSPAN7
AGMAT
SLC9A3R1
FST
TMEM9
MTTP
MPV17L
ACAT1
H2AFJ
FASN
CADPS2
DPYD
CREB3L3
CFI
PEG10
AGMO
CPD
MGST1
PAQR9
HS3ST3B1
SLC39A14
TTC39C
ROBO1
PTPRG
ETV1
APOE
SEMA3D
SERPING1
C1QBP
IQGAP2
FURIN
DCPS
NME1
TMOD1
GPRC5C
NME2
CDK14
PCM1
RELN
CXCL2
LPAR3
OSMR
PCDH9
PRSS23
TMSB10
DSTN
EDN1
ANXA3
ANXA2
S100A11
RHOC
C12orf75
TIMP1
FLNA
TMSB4X
PAWR
NRP2
TNFRSF21
TGFB2
TPM1
BMP4
CD9
PYGB
FSCN1
LAMA5
AHNAK
LAMB1
SLC2A3
HSPG2
CRIM1
MID1
TMEM156
LTBP1
TIMP2
TWSG1
BICC1
WWTR1
BCL2L1
SOCS2
GPRC5A
VAMP8
EREG
PXDC1
CD44
DSC2
TGFBI
JUP
MISP
ASAP1
RAB17
LDHB
APOBEC3B
INHBB
TACC2
SYNPO
PIEZO2
PTBP3
LIPH
PARD6B
ITPR3
ITM2C
APP
CABLES1
CTSZ
TM4SF1
SAT1
EPAS1
HIP1
RBM24
PLAU
CHST3
TRAM1
IGFBP4
EPCAM
DKK1
BIN1
CYB5B
AGR2
MT2A
state axis
HuH-7 axis
up − down score
z-score

#### Figure S6. Genes bound to the PLC/PRF/5 state axis behave similarly in HuH-7 cells and in HCC05 tumour cells. Related to Figure 4.

(A) The 145 genes correlated with the PLC/PRF/5 state axis ($|\rho| > 0.3$ across control cells, PROM1 excluded), as two blocks: genes of the hepatocyte-like C0 end ($\rho < -0.3$; n = 69, left) and of the opposite end ($\rho > +0.3$; n = 76, right). Each block shows the same genes in the same row order in three datasets: PLC/PRF/5 control cells along the state axis (n = 6,589; Figure 2), HuH-7 cells along their own diffusion axis (n = 2,075; Figure 4B), and HCC05 aneuploid tumour cells ordered by a PROM1 signature score (up − down; n = 866, including 301 PROM1-detected cells; computed as in Figure 4F, but from the 85 signature genes that are not shown in this panel, so the ordering does not reuse the displayed genes). Columns, 40 quantile bins of the respective ordering; values are mean log-normalized expression per bin (PLC/PRF/5 computed per pool, then averaged), z-scored per gene across bins. Rows are sorted by the HuH-7 gradient, with genes running against the direction of their PLC block placed at the bottom. Bars above each heatmap, percentage of PROM1+ cells (≥ 1 UMI) per bin; the row between bars and heatmap, PROM1 itself. Directional concordance with the PLC/PRF/5 blocks was 59 of 69 and 65 of 76 genes in HuH-7, and 53 of 69 and 65 of 76 genes in HCC05, respectively. Colored side bars identify the C0 and remaining-cell gene blocks.

HuH-7 and the HCC05 case are re-analyses of public data (GSE103867 and GSE290925) comprising one single-cell library and one patient, so cells are the units and the comparison is descriptive. The genes were selected on the PLC/PRF/5 axis, so the PLC/PRF/5 panel is circular by construction, whereas the HuH-7 analysis uses an independently derived axis and HCC05 is ordered using signature genes outside the displayed set. Values in Table S3 (axis_genes_three_datasets, axis_bins_three_datasets, hcc05_score_vs_prom1).

#### Table S5. Custom EGFP probe oligonucleotides for Chromium Flex, related to Figures 2, 3 and S2 and STAR Methods.

Sequences are shown 5′ to 3′ as ordered. The three LHS probes were common to all samples; each sample received the three RHS probes bearing its assigned barcode (BC001–BC004). LHS, left-hand-side EGFP-targeting probe, paired with the matching RHS probes; RHS, right-hand-side EGFP-targeting probe; EGFP, enhanced green fluorescent protein.

| Oligonucleotide | Sequence (5′ to 3′) | Role / sample barcode |
|---|---|---|
| EGFP-LHS-1 | CCTTGGCACCCGAGAATTCCAGGTAGTGGTCGGCGAGCTGCACGCT | LHS; all samples. |
| EGFP-LHS-2 | CCTTGGCACCCGAGAATTCCAAGGGTGTCGCCCTCGAACTTCACCT | LHS; all samples. |
| EGFP-LHS-3 | CCTTGGCACCCGAGAATTCCAATGGTGCGCTCCTGGACGTAGCCTT | LHS; all samples. |
| BC001-EGFP-RHS-1 | /5Phos/GCCGTCCTCGATGTTGTGGCGGATCACGCGGTTAGCACGTANNACTTTAGGCGGTCCTAGCAA | RHS; BC001; 5′ phosphate. |
| BC001-EGFP-RHS-2 | /5Phos/CGGCGCGGGTCTTGTAGTTGCCGTCACGCGGTTAGCACGTANNACTTTAGGCGGTCCTAGCAA | RHS; BC001; 5′ phosphate. |
| BC001-EGFP-RHS-3 | /5Phos/CGGGCATGGCGGACTTGAAGAAGTCACGCGGTTAGCACGTANNACTTTAGGCGGTCCTAGCAA | RHS; BC001; 5′ phosphate. |
| BC002-EGFP-RHS-1 | /5Phos/GCCGTCCTCGATGTTGTGGCGGATCACGCGGTTAGCACGTANNAACGGGAACGGTCCTAGCAA | RHS; BC002; 5′ phosphate. |
| BC002-EGFP-RHS-2 | /5Phos/CGGCGCGGGTCTTGTAGTTGCCGTCACGCGGTTAGCACGTANNAACGGGAACGGTCCTAGCAA | RHS; BC002; 5′ phosphate. |
| BC002-EGFP-RHS-3 | /5Phos/CGGGCATGGCGGACTTGAAGAAGTCACGCGGTTAGCACGTANNAACGGGAACGGTCCTAGCAA | RHS; BC002; 5′ phosphate. |
| BC003-EGFP-RHS-1 | /5Phos/GCCGTCCTCGATGTTGTGGCGGATCACGCGGTTAGCACGTANNAGTAGGCTCGGTCCTAGCAA | RHS; BC003; 5′ phosphate. |
| BC003-EGFP-RHS-2 | /5Phos/CGGCGCGGGTCTTGTAGTTGCCGTCACGCGGTTAGCACGTANNAGTAGGCTCGGTCCTAGCAA | RHS; BC003; 5′ phosphate. |
| BC003-EGFP-RHS-3 | /5Phos/CGGGCATGGCGGACTTGAAGAAGTCACGCGGTTAGCACGTANNAGTAGGCTCGGTCCTAGCAA | RHS; BC003; 5′ phosphate. |
| BC004-EGFP-RHS-1 | /5Phos/GCCGTCCTCGATGTTGTGGCGGATCACGCGGTTAGCACGTANNATGTTGACCGGTCCTAGCAA | RHS; BC004; 5′ phosphate. |
| BC004-EGFP-RHS-2 | /5Phos/CGGCGCGGGTCTTGTAGTTGCCGTCACGCGGTTAGCACGTANNATGTTGACCGGTCCTAGCAA | RHS; BC004; 5′ phosphate. |
| BC004-EGFP-RHS-3 | /5Phos/CGGGCATGGCGGACTTGAAGAAGTCACGCGGTTAGCACGTANNATGTTGACCGGTCCTAGCAA | RHS; BC004; 5′ phosphate. |

#### Table S6. Oligonucleotides for endogenous PROM1 ALFA-tag knock-in and genotyping, related to Figure 5 and STAR Methods.

Guide spacers and primer sequences are written 5′ to 3′. Donor sequences are shown in the ordered 5′ to 3′ orientation; lowercase letters in the source sequence identify the ALFA-tag cassette and were retained. Expected product sizes are those specified in the supplied design records. ssODN, single-stranded donor oligonucleotide; o/o, outside/outside; i/o, inside/outside; bp, base pairs.

| Oligonucleotide | Sequence (5′ to 3′) | Purpose / expected product |
|---|---|---|
| sgRNA1 target segment | CAGATGCTCCTAAGGCTTGG | PROM1 coding-strand orientation; extracellular insertion site with a reverse-oriented guide. PAM excluded. |
| sgRNA4 spacer | TGTCGTTGCTGTAACAAATG | Forward-oriented guide; PAM excluded. |
| sgRNA1 ssODN donor | TGGGTCTCATAATTTGTTGCAGGCAATTCATAATTCCAAGCCTTAGGAG Ccgagccaccccccgcccggttcggtcaggcggcggcgcagttcttcttccaggcggctcggagagcca cctcctcCATCTGTGGATGAAGGCTGCCCTCCTGAAAAGGAGTTCCCGCA CAGCCCCAGC | 5′→3′; extracellular-site insertion. |
| sgRNA4 ssODN donor | CTCAGGAAGGGCCCATTTTCCTTCTGTCGCTGGTGCATTTCTCCACCA Cacgagccaccccccgcccggttcggtcaggcggcggcgcagttcttcttccaggcggctcggagagcc accagatcCTTTGTTACAGCAACGACACATACAAAAGAAATACCCCACCAG AGGCATCAG | 5′→3′; first intracellular-loop insertion. |
| sgRNA1 o/o primer 1 | CAAGCCAGCCTCAGACAGAA | Pair with sgRNA1 o/o primer 2; expected 632 bp tagged and 557 bp untagged. |
| sgRNA1 o/o primer 2 | TCCTCGCAACCTATGTAACCTG | Pair with sgRNA1 o/o primer 1. |
| sgRNA4 o/o primer 1 | CAGTGTGCAAAGCGCCTTAT | Pair with sgRNA4 o/o primer 2; expected 1,000 bp tagged and 925 bp untagged. |
| sgRNA4 o/o primer 2 | TGTGAGACACTCTTCATCAGCC | Pair with sgRNA4 o/o primer 1. |
| ALFA-tag inside primer (i/o) | CGAGCCGCCTGGAAGAAGAA | Shared internal primer; forward orientation in the sgRNA4 inside/outside assay. |
| sgRNA4 outside primer (i/o) | TGAGACACTCTTCATCAGCCT | Outside reverse primer paired with the ALFA-tag inside primer; sequence-predicted tagged product, 211 bp. |

#### Separate supplemental files

##### Table S1. Acute PROM1 perturbation transcriptomics, related to Figure 1 and Figures S1 and S3.

Differential expression, reciprocal response genes, pathway and regulon analyses, and PROM1 locus specificity.

##### Table S2. PLC/PRF/5 single-cell state analysis, related to Figures 2 and 3 and Figures S2 and S3.

Cell-state markers, control-axis associations, regulons, pathway scores and robustness analyses.

##### Table S3. HuH-7 and human HCC analyses, related to Figure 4 and Figure S6.

HuH-7 state-axis, pathway and regulon analyses; HCC cohort annotations and HCC05 scores; cross-dataset analysis of 145 PLC-associated genes. HCC05 heatmap ordering uses 85 response-signature genes outside the displayed set. Cell-type marker annotation and cell-cycle scoring were informed by published single-cell studies of liver and tumor populations.[1,2–5]

##### Table S4. CD133 affinity-purification proteomics, related to Figure 5.

Detection-based and LFQ candidate screens, peptide-identification support and quantitative model outputs.

##### Data S1. Numerical source data, related to Figures 1–5 and Figures S1–S6.

Available values underlying validation assays, immunoblot ratios, CAM graft measurements and statistical models, and recoverable HUVEC graph summaries. Workbook sheet names identify each panel; source-data availability is documented in its README.

##### Data S2. Uncropped immunoblots, related to Figure 5 and Figures S1 and S2.

Original blot images with the supplied lane, target and molecular-weight annotations; blue outlines indicate displayed or quantified regions where supplied. These source images support the cropped figure panels.
