## Supplementary material for "Endogenous PROM1 activation redistributes liver cancer cells along a preexisting differentiation axis": Data_S2

Data S2. Uncropped immunoblots

#### Figure S1G + S2A — Ponceau S, whole membrane (uncropped)

Blot 2024-08-20, total protein before immunodetection. Two ladder lanes; between and right of them the three stable-line lanes (Figure S2A) and the six knockdown lanes (Figure S1G).

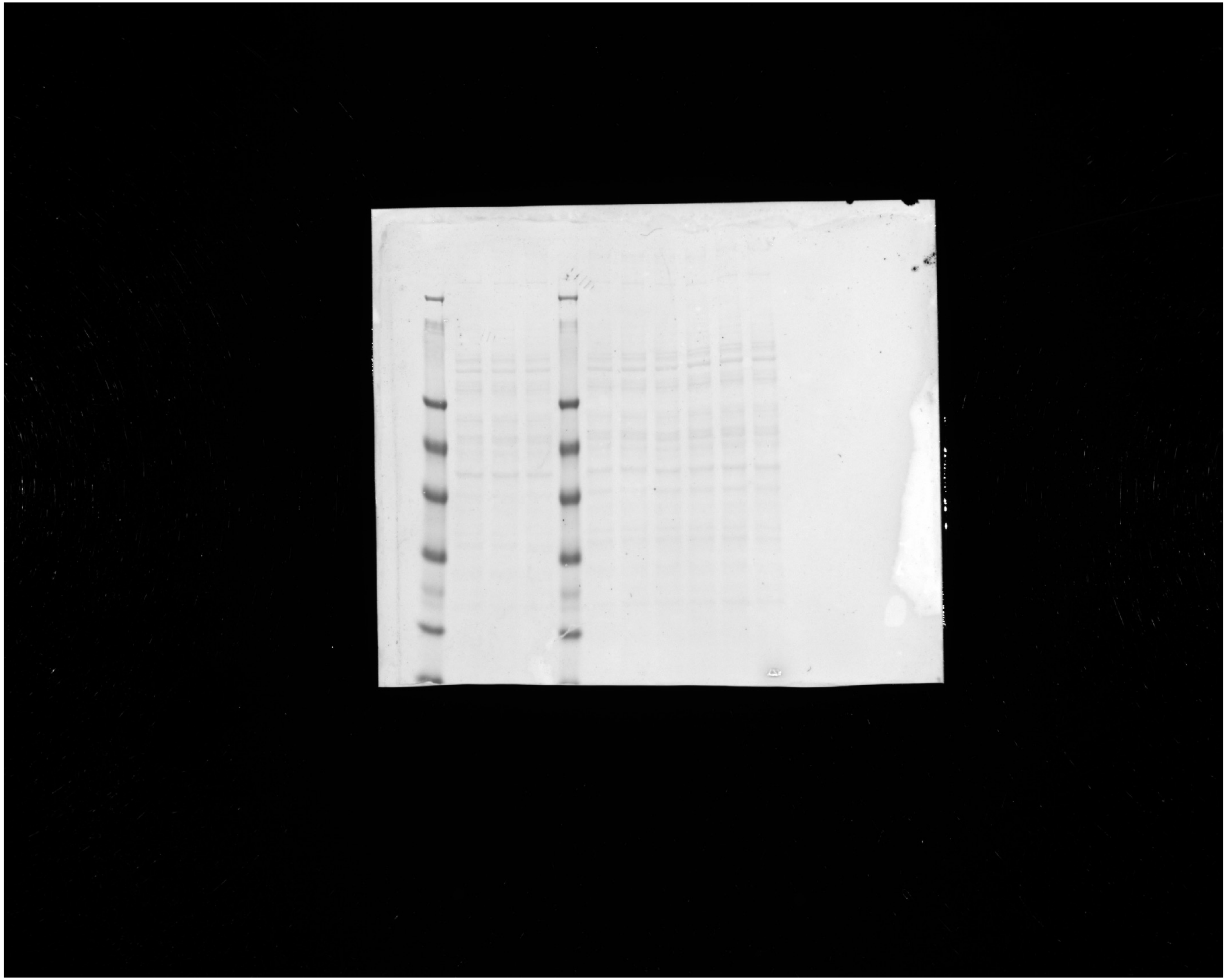

Data S2. Uncropped immunoblots

#### Figure S1G + S2A — β-Actin, whole membrane (uncropped)

Blot 2024-08-20. Left three lanes: stable CRISPRa / SAMph / GFP (Figure S2A). Right six lanes: LoF 1, Ctrl 1, LoF 2, Ctrl 2, LoF 3, Ctrl 3 (Figure S1G).

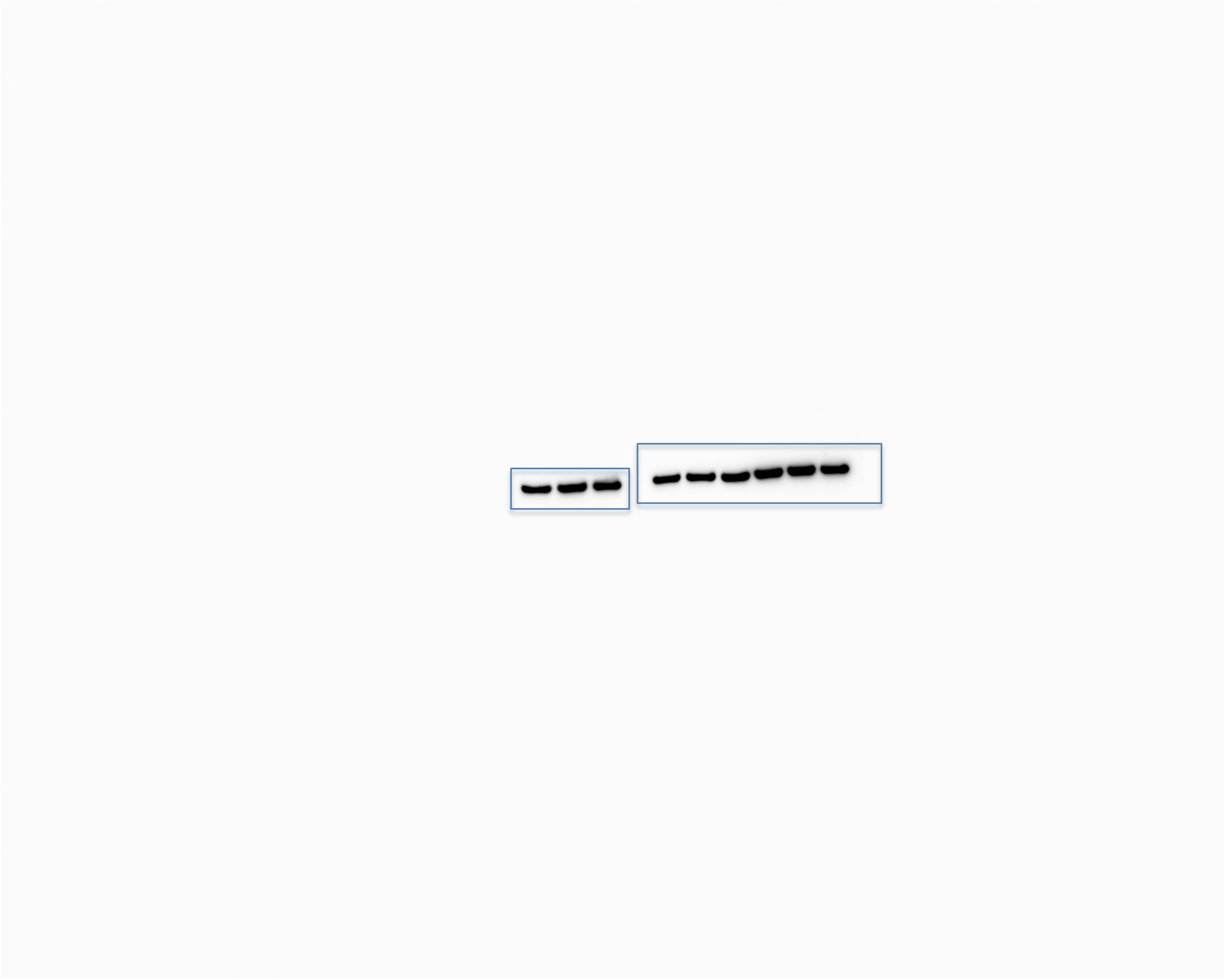

Data S2. Uncropped immunoblots

#### Figure S1G + S2A — CD133, whole membrane (uncropped)

Same membrane and lane order as the β-Actin slide.

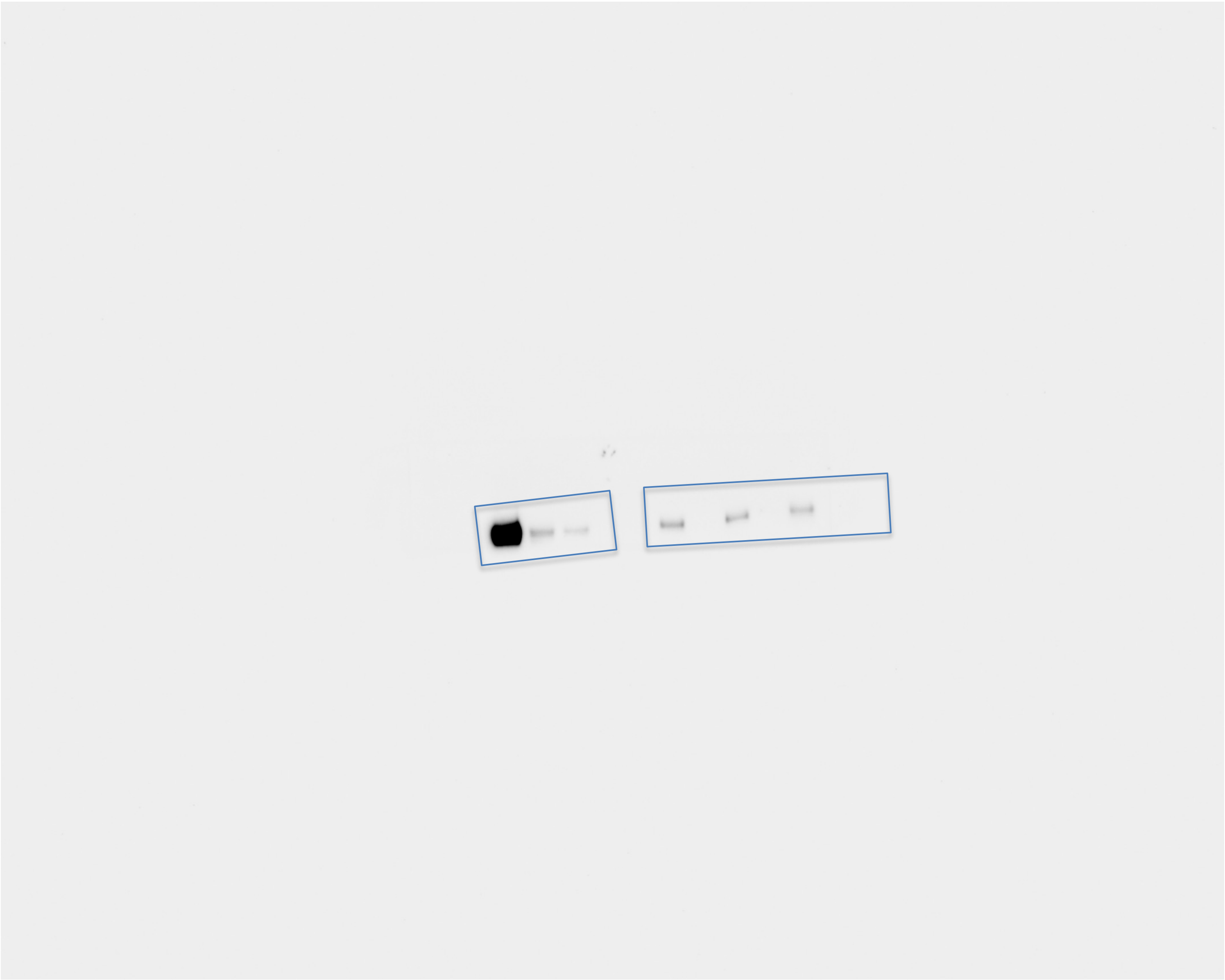

Data S2. Uncropped immunoblots

### Figure S1F — Ponceau S, membrane 1 (CD133 + β-Actin), whole membrane (uncropped)

Lanes alternate CaT 1, SAMPHT 1, CaT 2, SAMPHT 2, CaT 3, SAMPHT 3.
Lanes, left to right: GoF 1, Ctrl 1, GoF 2, Ctrl 2, GoF 3, Ctrl 3 — CRISPRa and SAMph lysates in alternating lanes, labelled as in Figure S1F. Same order on all Figure S1F slides.

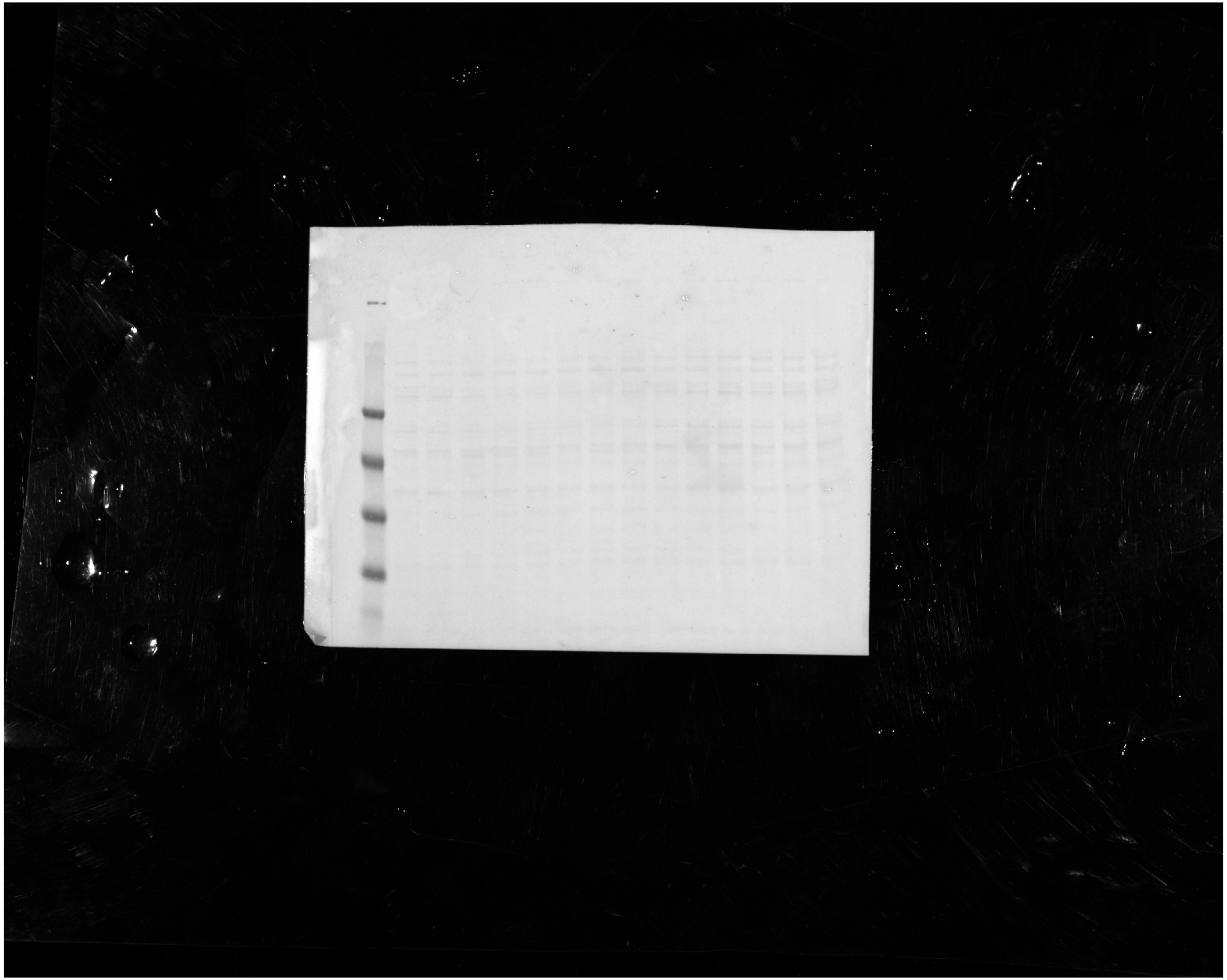

Data S2. Uncropped immunoblots

### Figure S1F — CD133, developed strip (merged: colorimetric + chemiluminescence)

Physically cut strip of membrane 1, therefore not a whole membrane.
Lanes, left to right: GoF 1, Ctrl 1, GoF 2, Ctrl 2, GoF 3, Ctrl 3 — CRISPRa and SAMph lysates in alternating lanes, labelled as in Figure S1F. Same order on all Figure S1F slides.

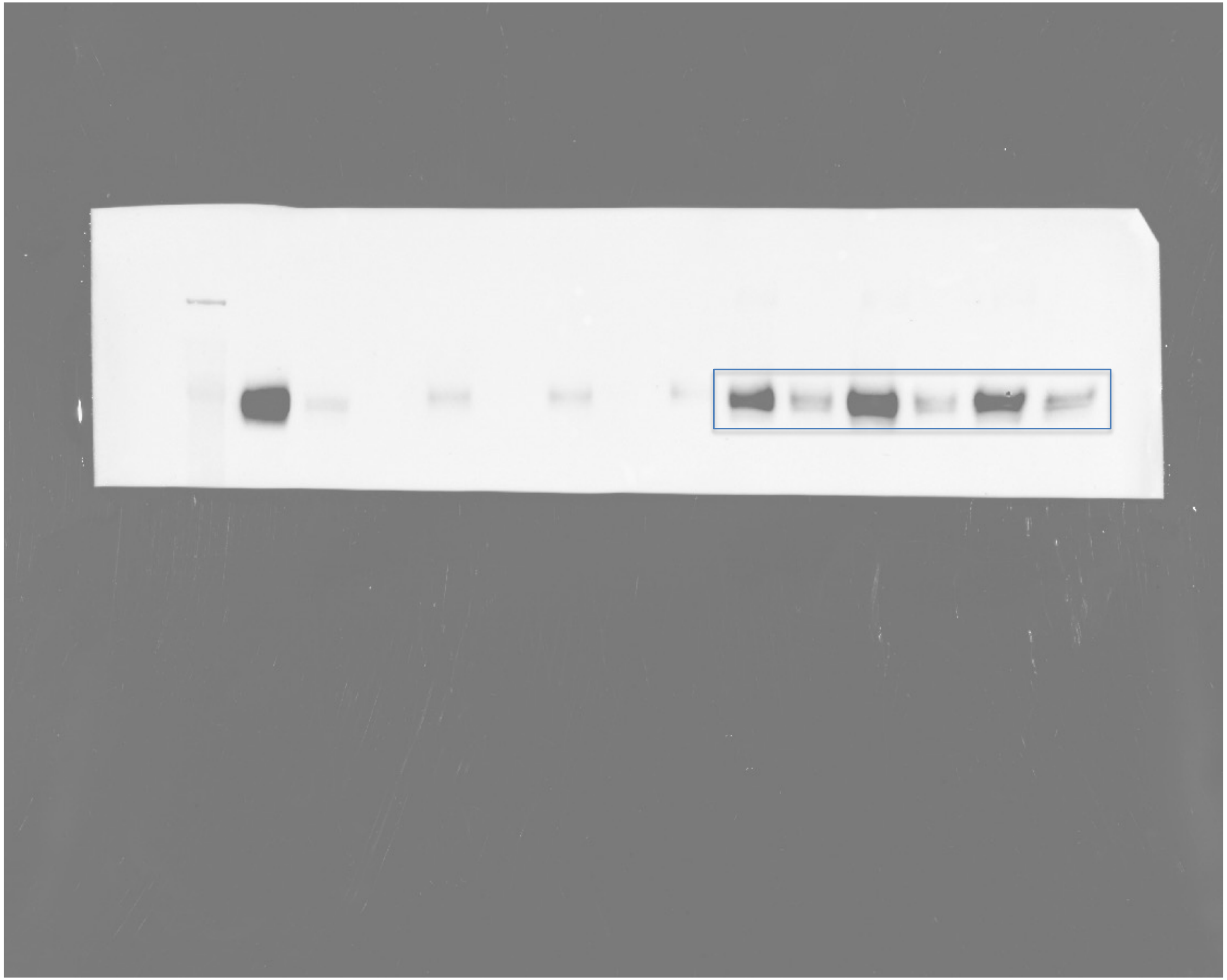

Data S2. Uncropped immunoblots

### Figure S1F — β-Actin, developed strip (merged)

Lower strip of the same membrane as CD133.
Lanes, left to right: GoF 1, Ctrl 1, GoF 2, Ctrl 2, GoF 3, Ctrl 3 — CRISPRa and SAMph lysates in alternating lanes, labelled as in Figure S1F. Same order on all Figure S1F slides.

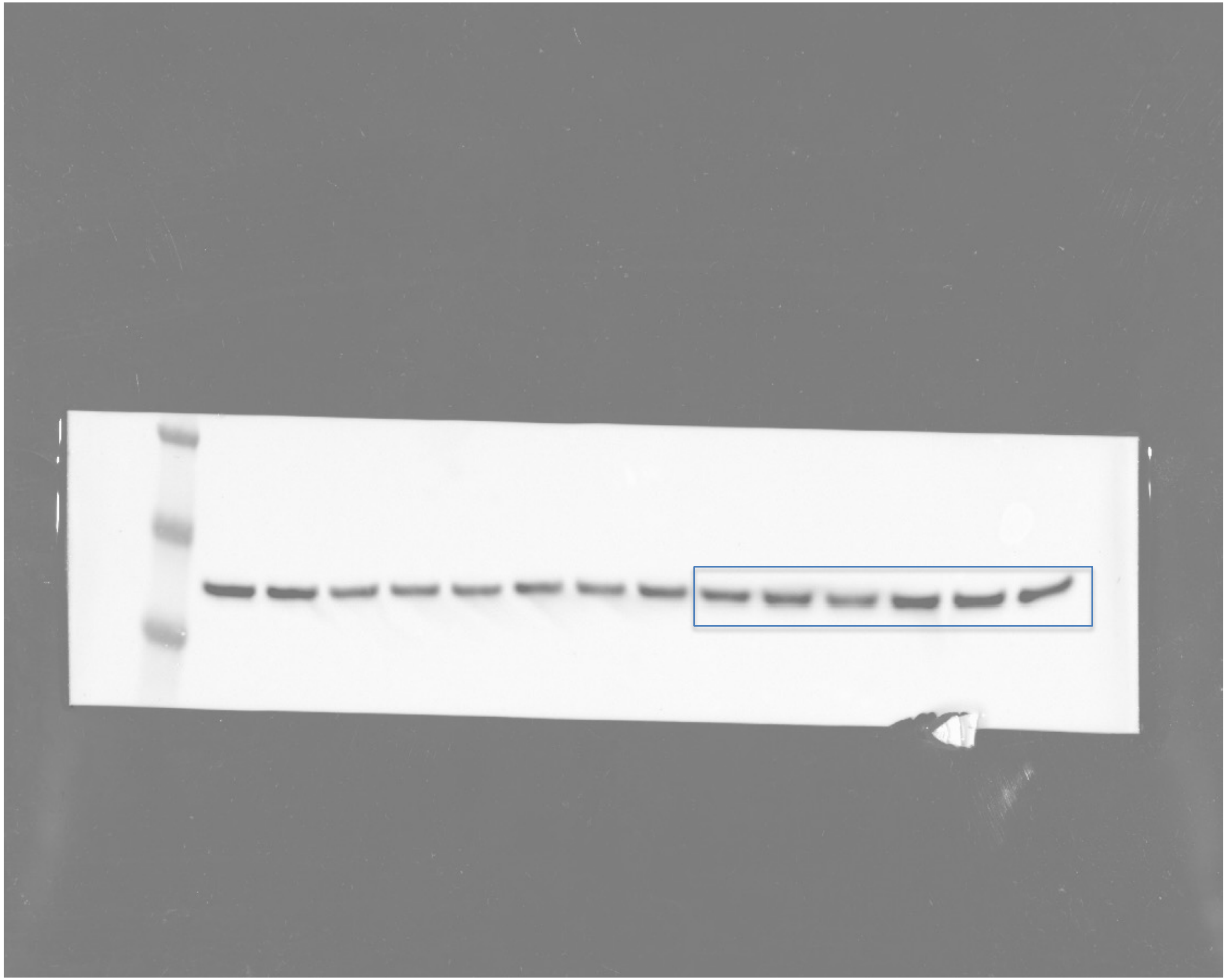

**Data S2. Uncropped immunoblots**

### Figure S1F — Ponceau S, CD44 membrane, whole membrane (uncropped)

Lanes, left to right: GoF 1, Ctrl 1, GoF 2, Ctrl 2, GoF 3, Ctrl 3 — CRISPRa and SAMph lysates in alternating lanes, labelled as in Figure S1F. Same order on all Figure S1F slides.

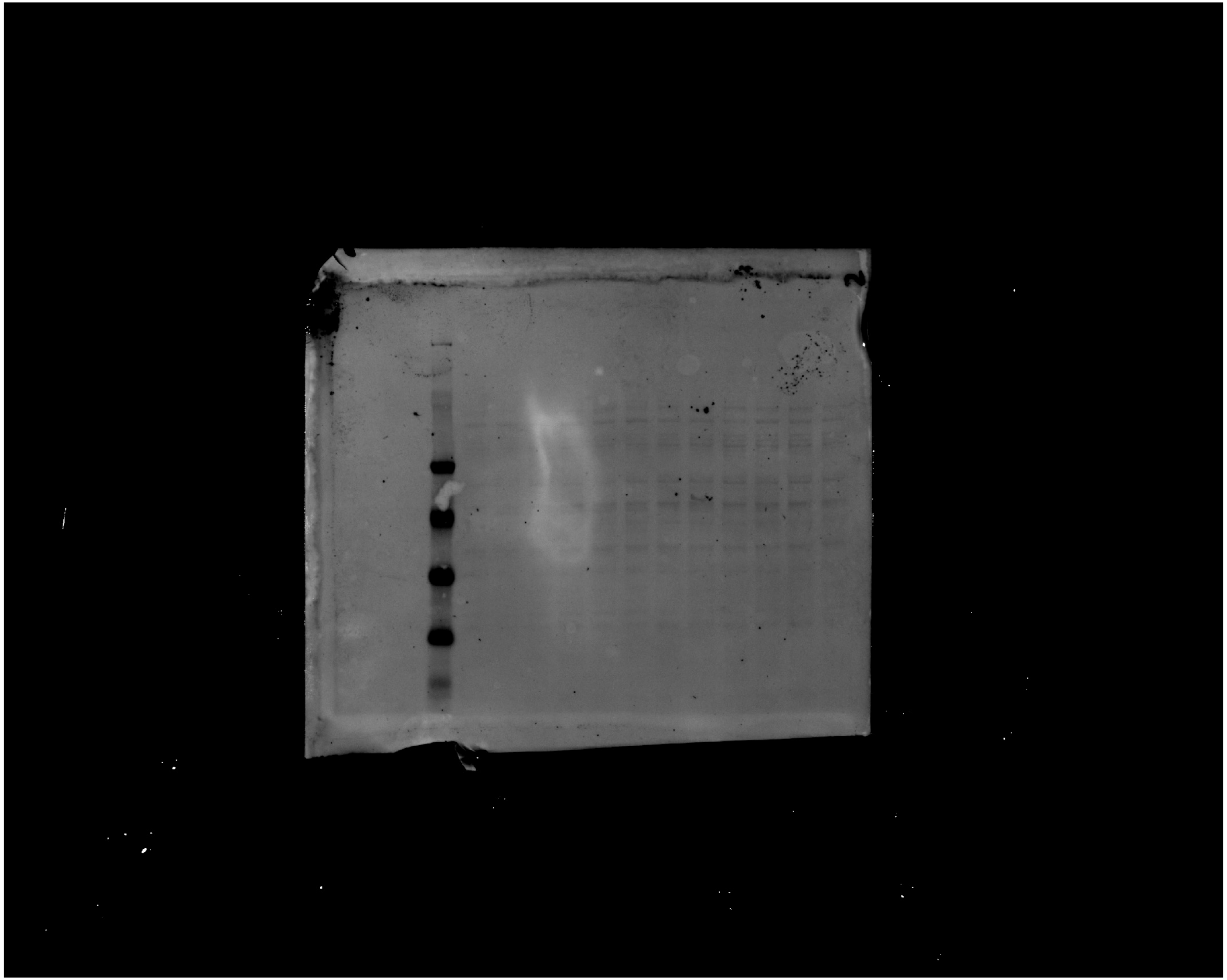

**Data S2. Uncropped immunoblots**

#### Figure S1F — CD44, developed strip (merged)

Lanes, left to right: GoF 1, Ctrl 1, GoF 2, Ctrl 2, GoF 3, Ctrl 3 — CRISPRa and SAMph lysates in alternating lanes, labelled as in Figure S1F. Same order on all Figure S1F slides.

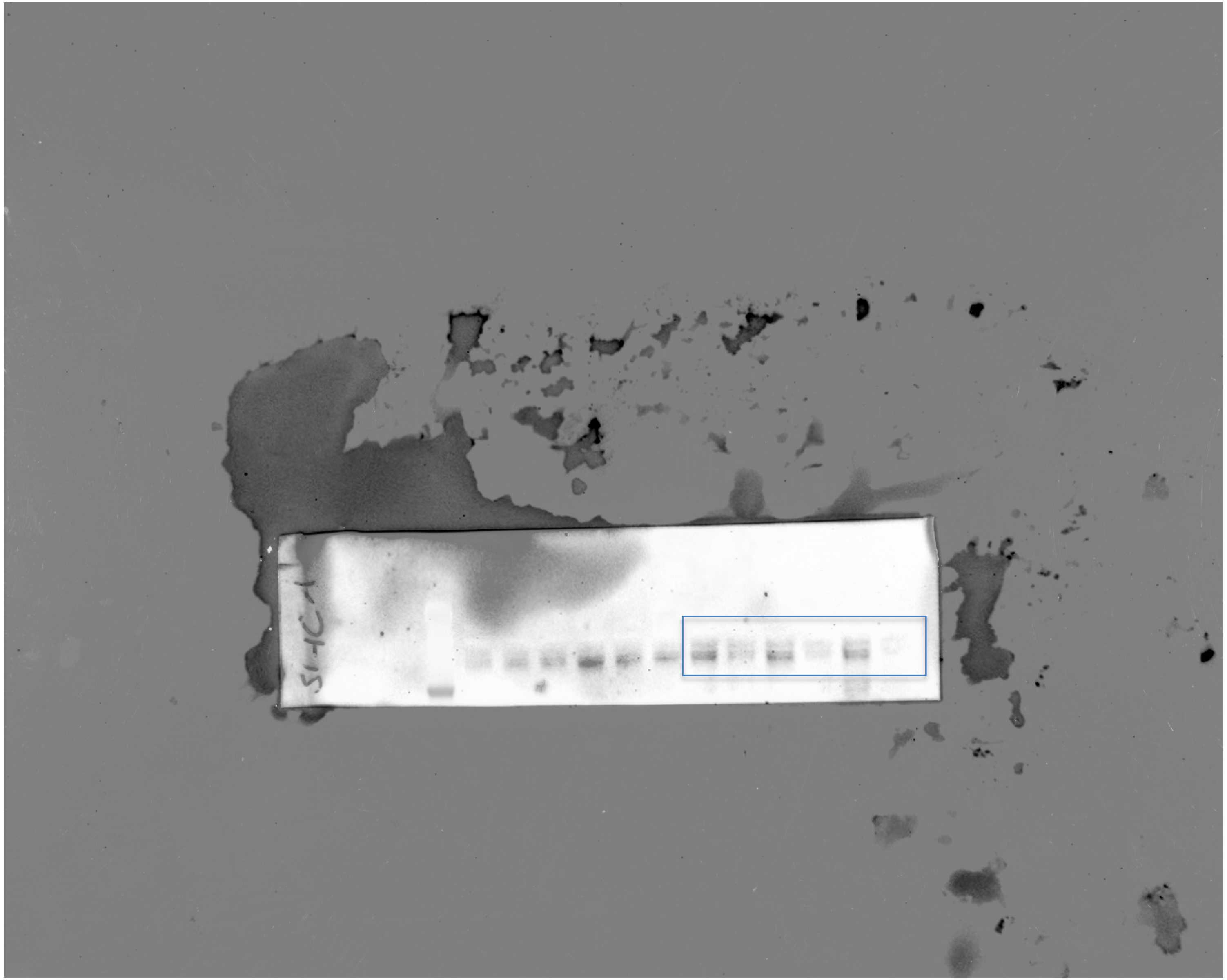

Data S2. Uncropped immunoblots

#### Figure S1F — Ponceau S, CXCR4 membrane, whole membrane (uncropped)

Lanes, left to right: GoF 1, Ctrl 1, GoF 2, Ctrl 2, GoF 3, Ctrl 3 — CRISPRa and SAMph lysates in alternating lanes, labelled as in Figure S1F. Same order on all Figure S1F slides.

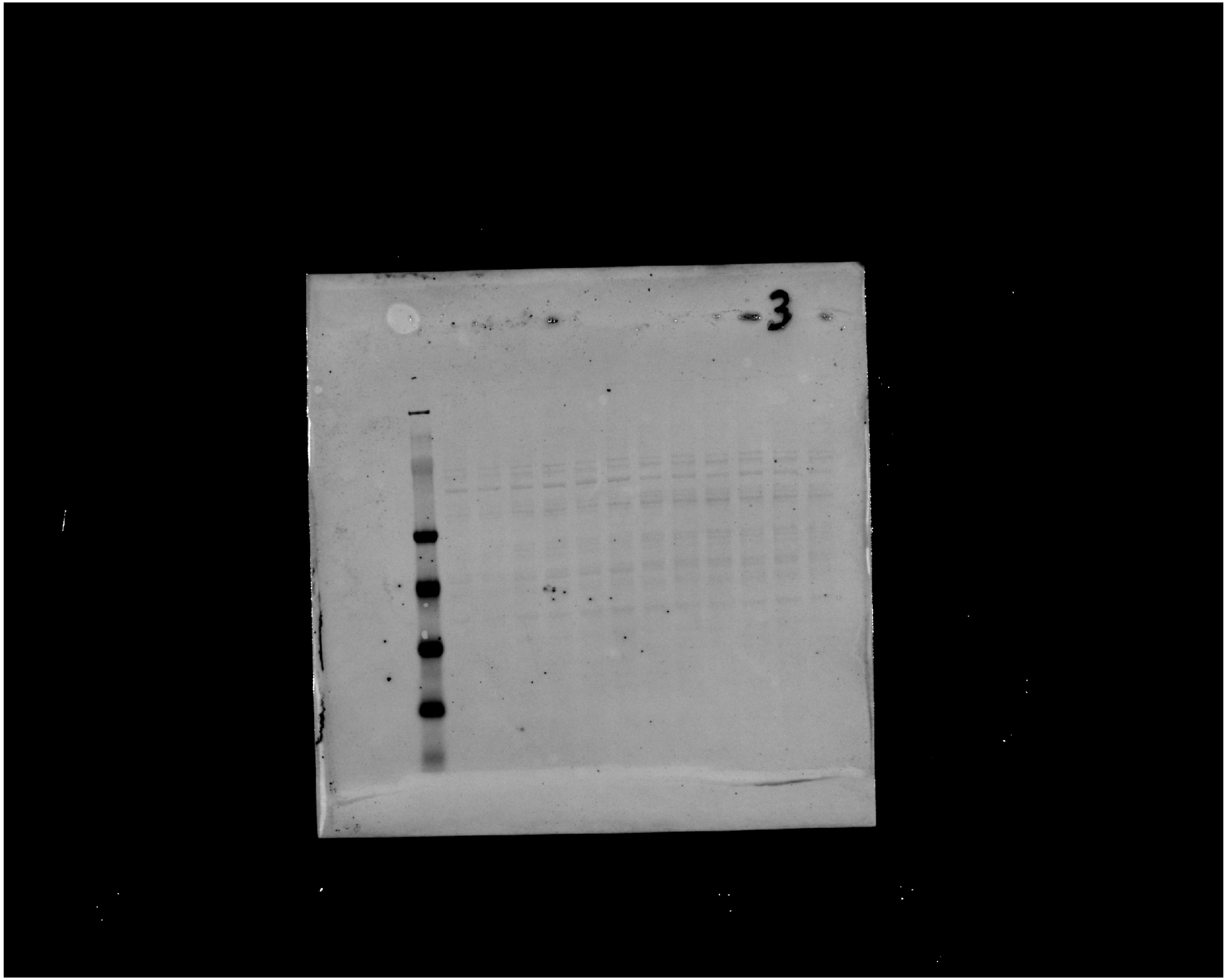

Data S2. Uncropped immunoblots

#### Figure S1F — CXCR4, developed strip (merged)

The two upper immunoreactive bands were quantified together; the lowest band was excluded.
Lanes, left to right: GoF 1, Ctrl 1, GoF 2, Ctrl 2, GoF 3, Ctrl 3 — CRISPRa and SAMph lysates in alternating lanes, labelled as in Figure S1F. Same order on all Figure S1F slides.

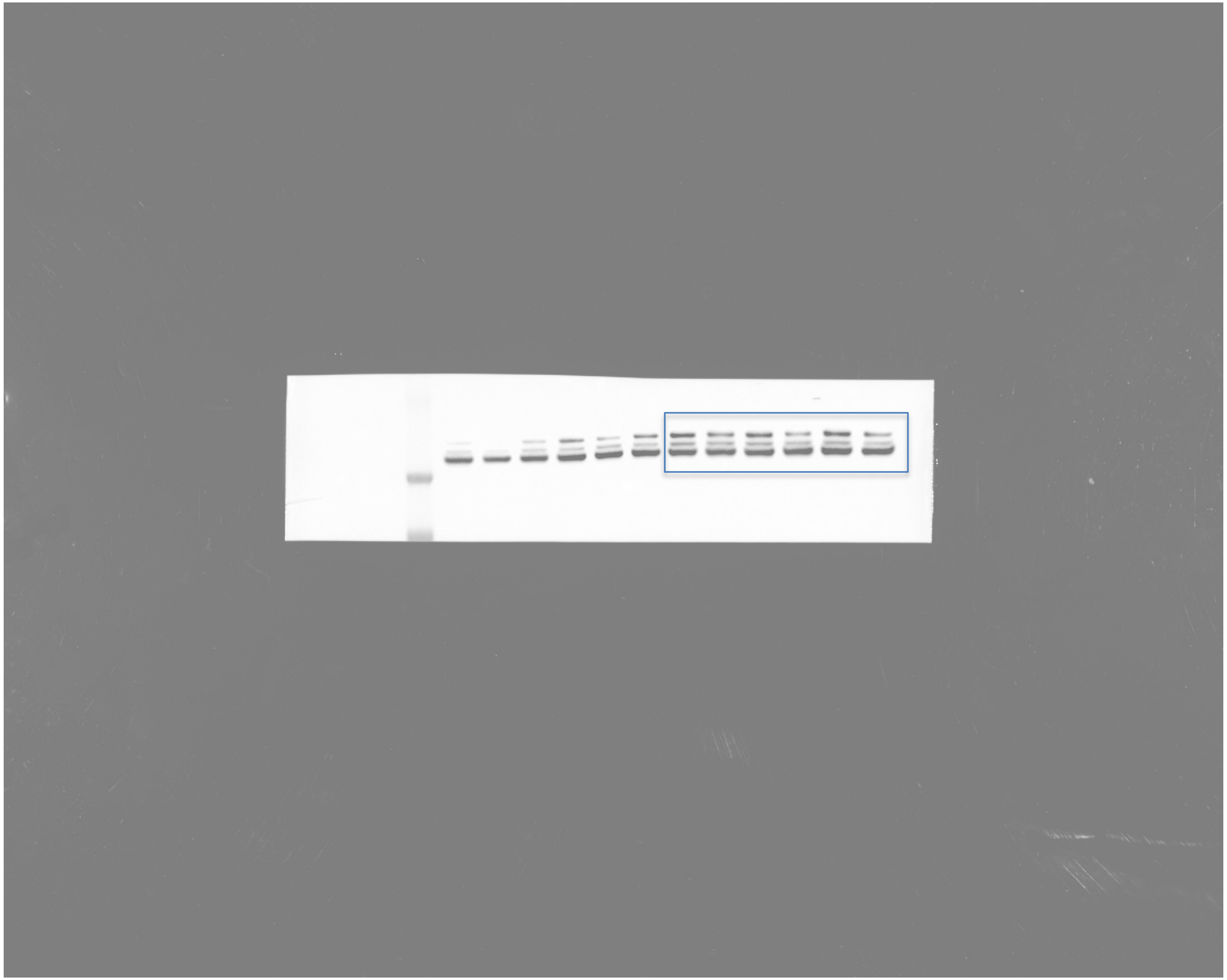

**Data S2. Uncropped immunoblots**

### Figure S1F — Ponceau S, KRT80 membrane, whole membrane (uncropped)

Lanes, left to right: GoF 1, Ctrl 1, GoF 2, Ctrl 2, GoF 3, Ctrl 3 — CRISPRa and SAMph lysates in alternating lanes, labelled as in Figure S1F. Same order on all Figure S1F slides.

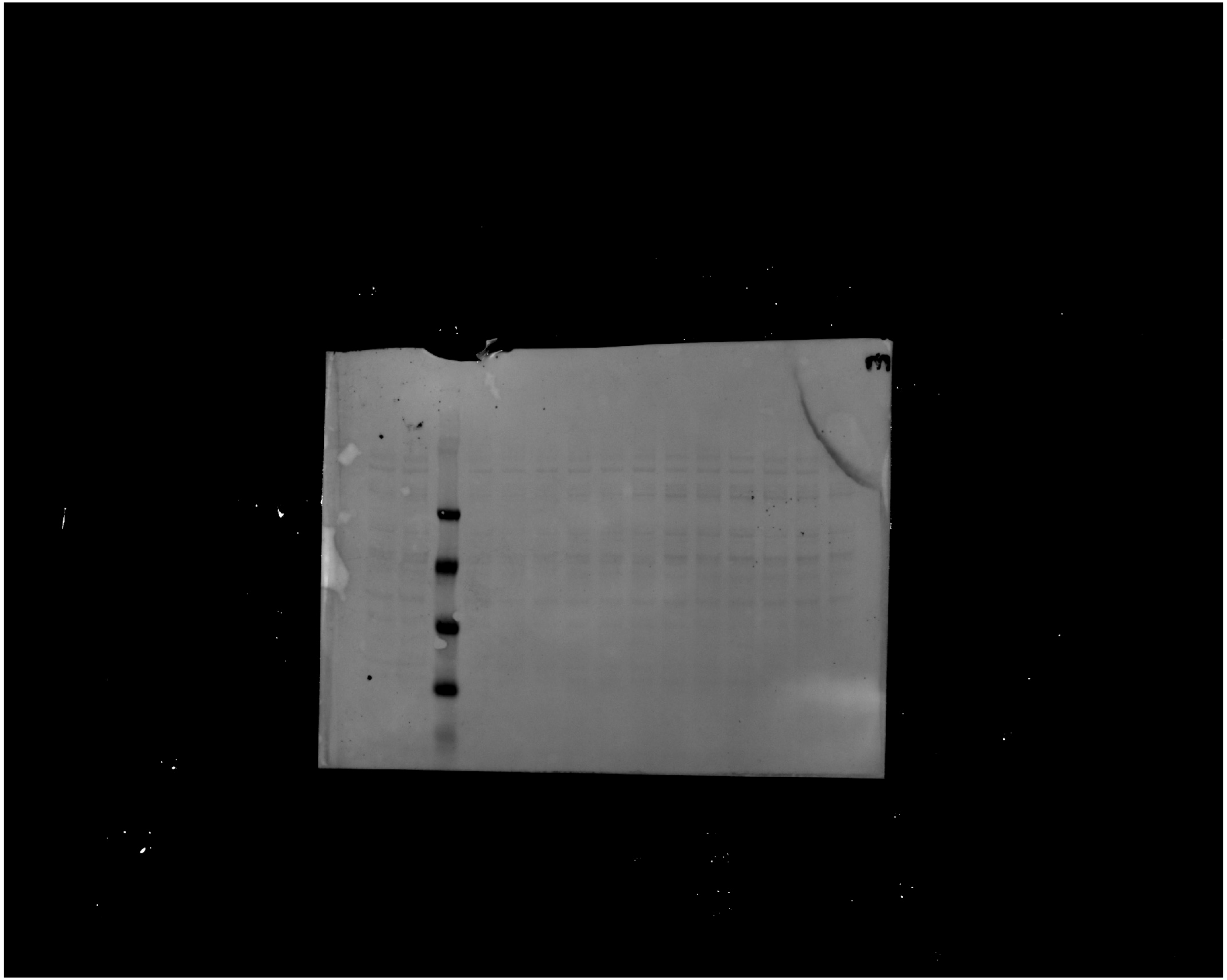

**Data S2. Uncropped immunoblots**

#### Figure S1F — KRT80, developed strip (merged)

Lanes, left to right: GoF 1, Ctrl 1, GoF 2, Ctrl 2, GoF 3, Ctrl 3 — CRISPRa and SAMph lysates in alternating lanes, labelled as in Figure S1F. Same order on all Figure S1F slides.

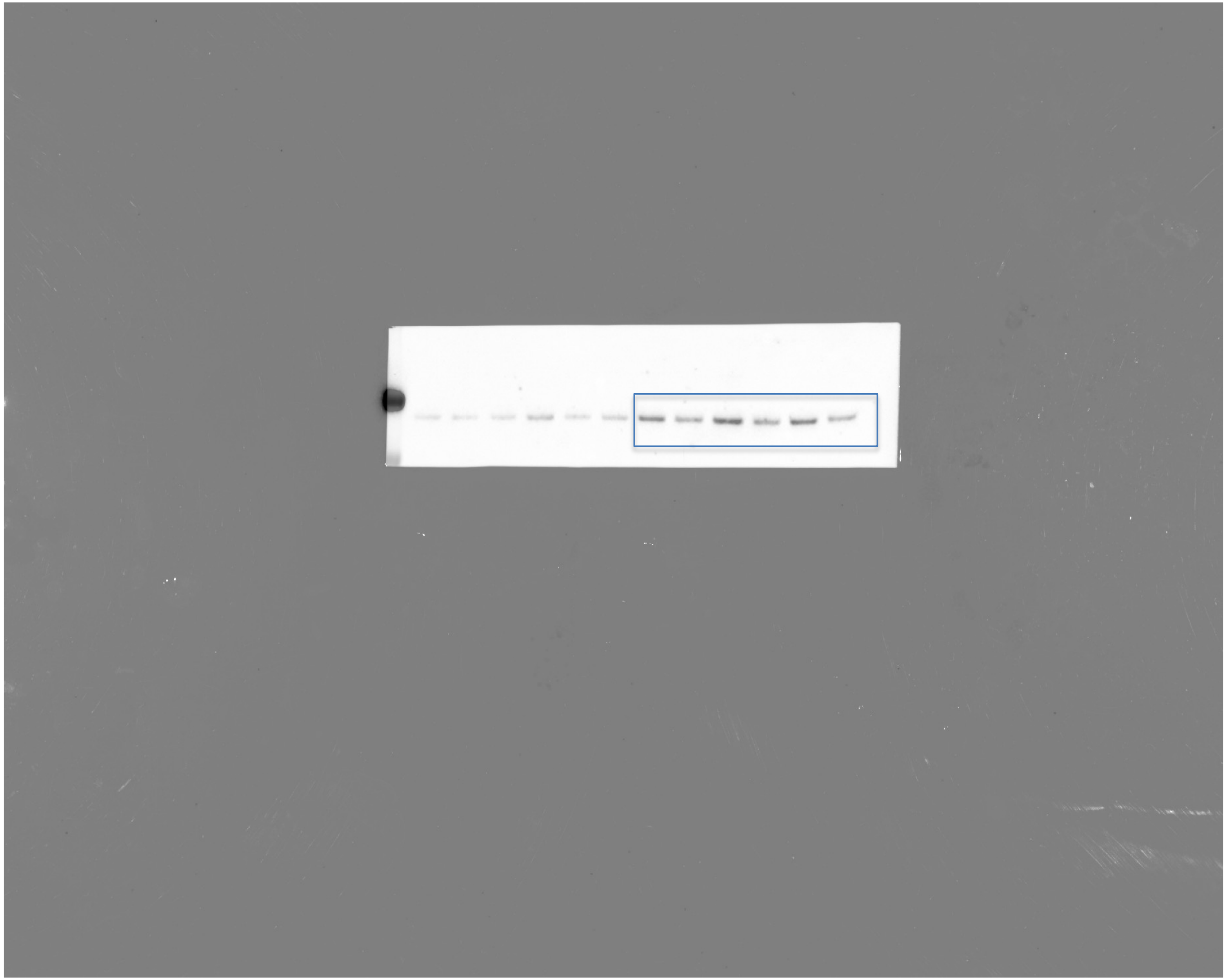

**Data S2. Uncropped immunoblots**

#### Figure 5E — CD133 input — chemiluminescence, whole membrane (uncropped)

2024-10-31 Chemiluminescence 26.574 s exposure 916x733 px 16-bit scan, linearly rescaled for display (61556-65005 of 0-65535); applied to the whole image.

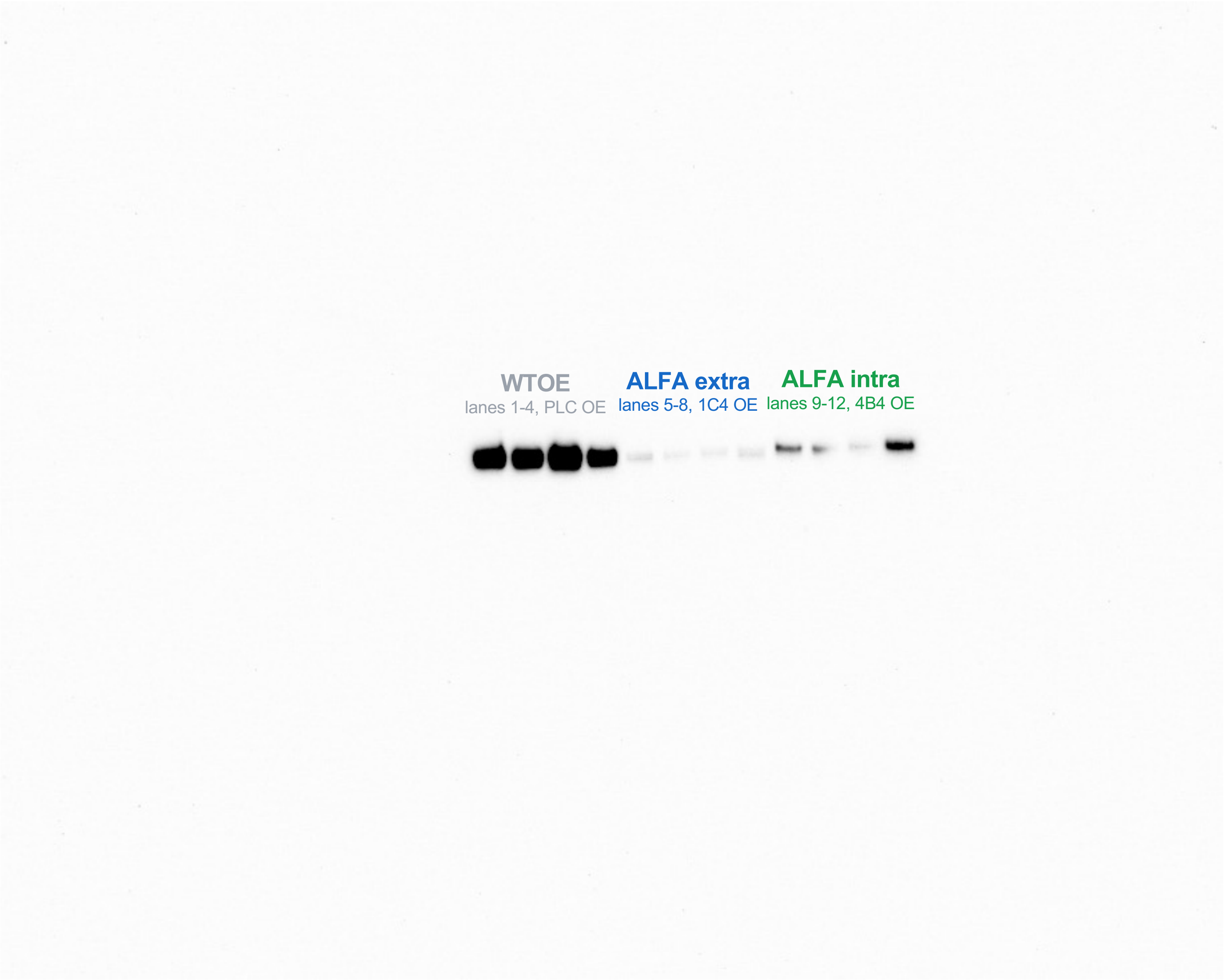

Data S2. Uncropped immunoblots

#### Figure 5E — CD133 IP — chemiluminescence, whole membrane (uncropped)

916x733 px  16-bit scan, linearly rescaled for display (48672-64985 of 0-65535); applied to the whole image.

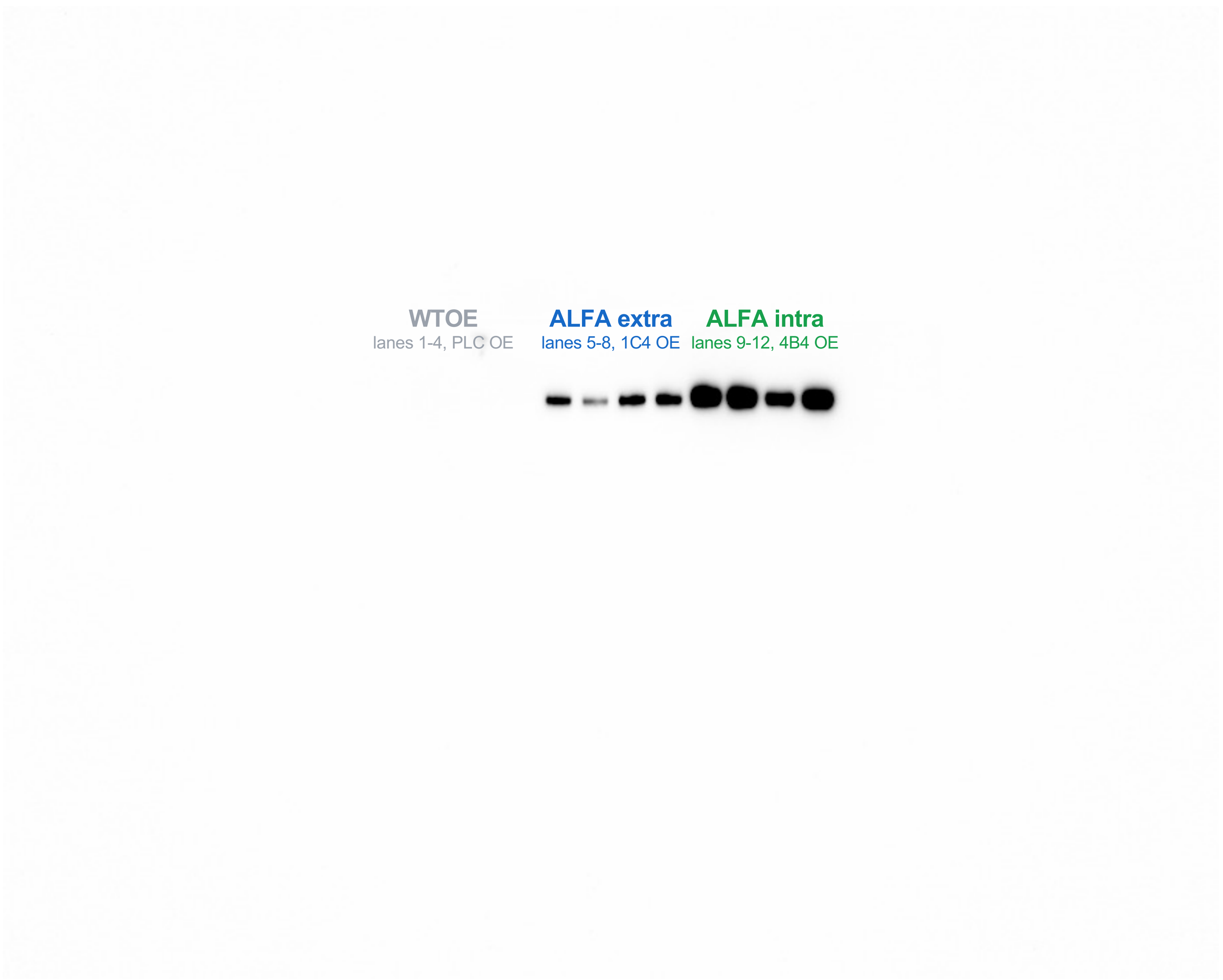

**Data S2. Uncropped immunoblots**

#### Figure 5E — TUBB6 input, whole membrane (uncropped)

2025-04-24  Chemiluminescence  5.95 s exposure  1375x1100 px  16-bit scan, linearly rescaled for display (35367-65007 of 0-65535); applied to the whole image.

**WTOE** lanes 1-4, PLC OE

**ALFA extra** lanes 5-8, 1C4 OE

**ALFA intra** lanes 9-12, 4B4 OE

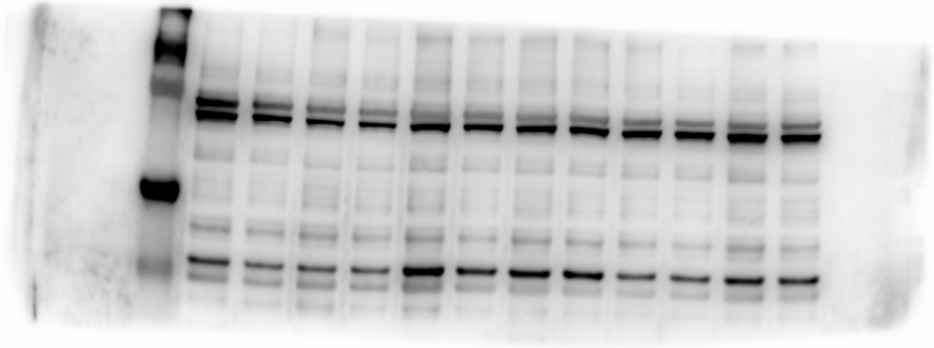

**Data S2. Uncropped immunoblots**

#### Figure 5E — TUBB6 IP, whole membrane (uncropped)

2025-04-24 Chemiluminescence 18.872 s exposure 1375x1100 px 16-bit scan, linearly rescaled for display (28965-64999 of 0-65535); applied to the whole image.

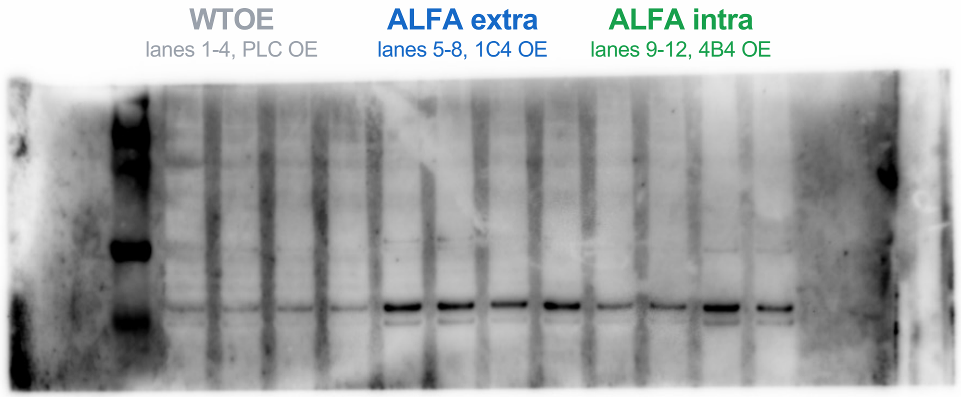

**Data S2. Uncropped immunoblots**

#### Figure 5E — DUSP9 input, whole membrane (uncropped)

916x733 px 16-bit scan, linearly rescaled for display (48046-65009 of 0-65535); applied to the whole image.

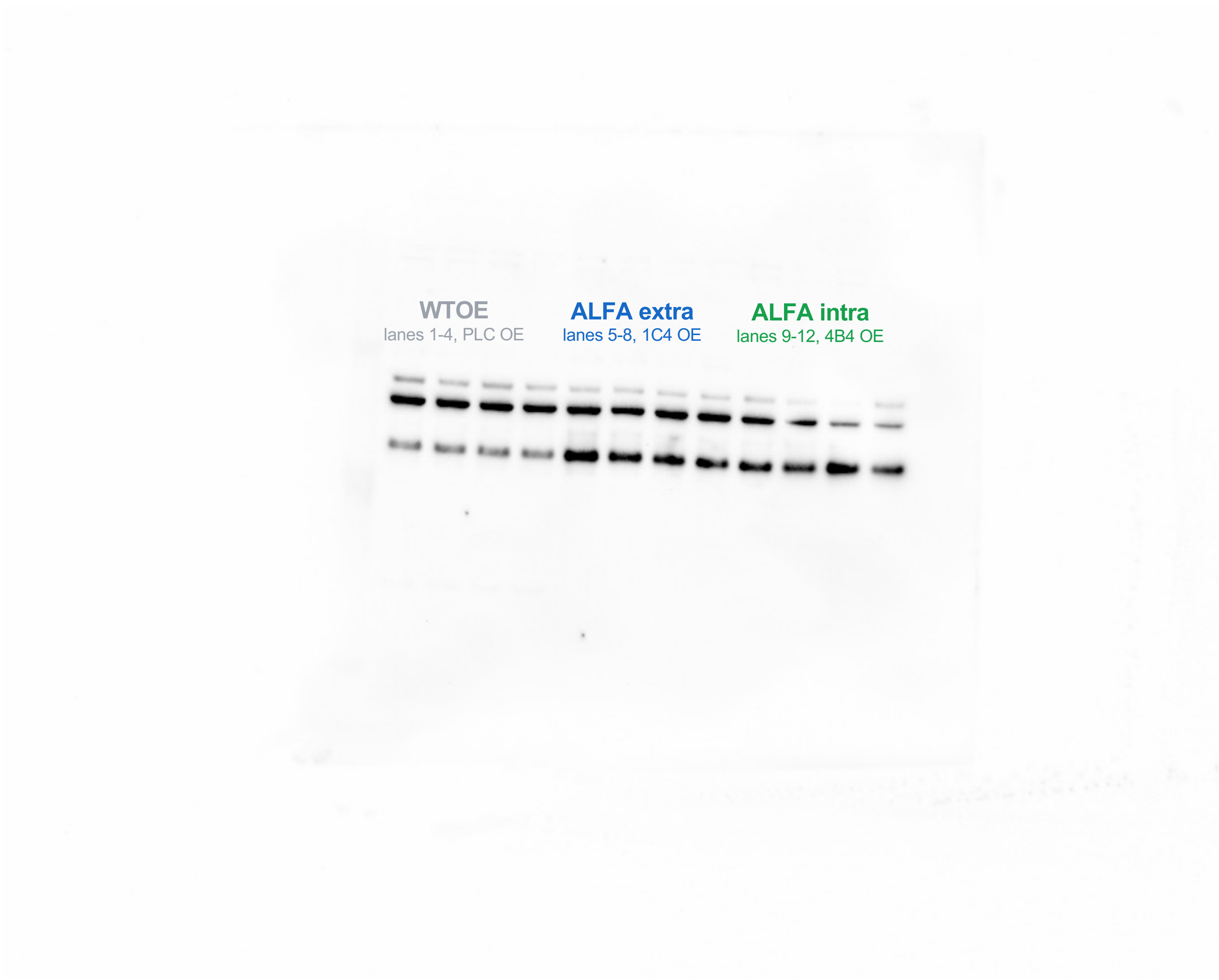

Data S2. Uncropped immunoblots

#### Figure 5E — DUSP9 IP, whole membrane (uncropped)

2025-04-23 Chemiluminescence 185.066 s exposure 916x733 px 16-bit scan, linearly rescaled for display (52197-64994 of 0-65535); applied to the whole image.

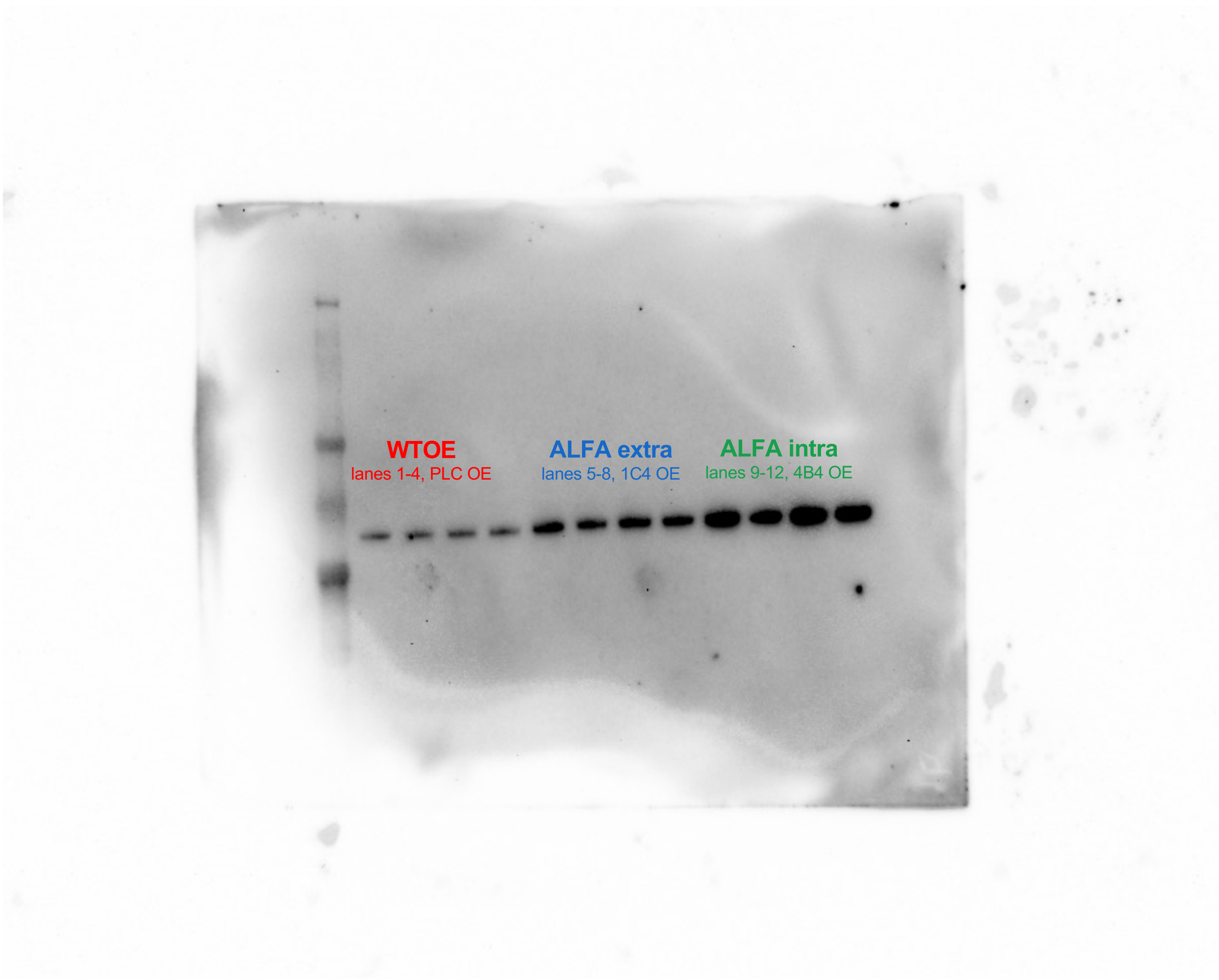

Data S2. Uncropped immunoblots

### Figure 5E — ATP6V1A input, whole membrane (uncropped)

2025-04-24  Chemiluminescence  366.958 s exposure  1375x1100 px  16-bit scan, linearly rescaled for display (37622-65003 of 0-65535); applied to the whole image.

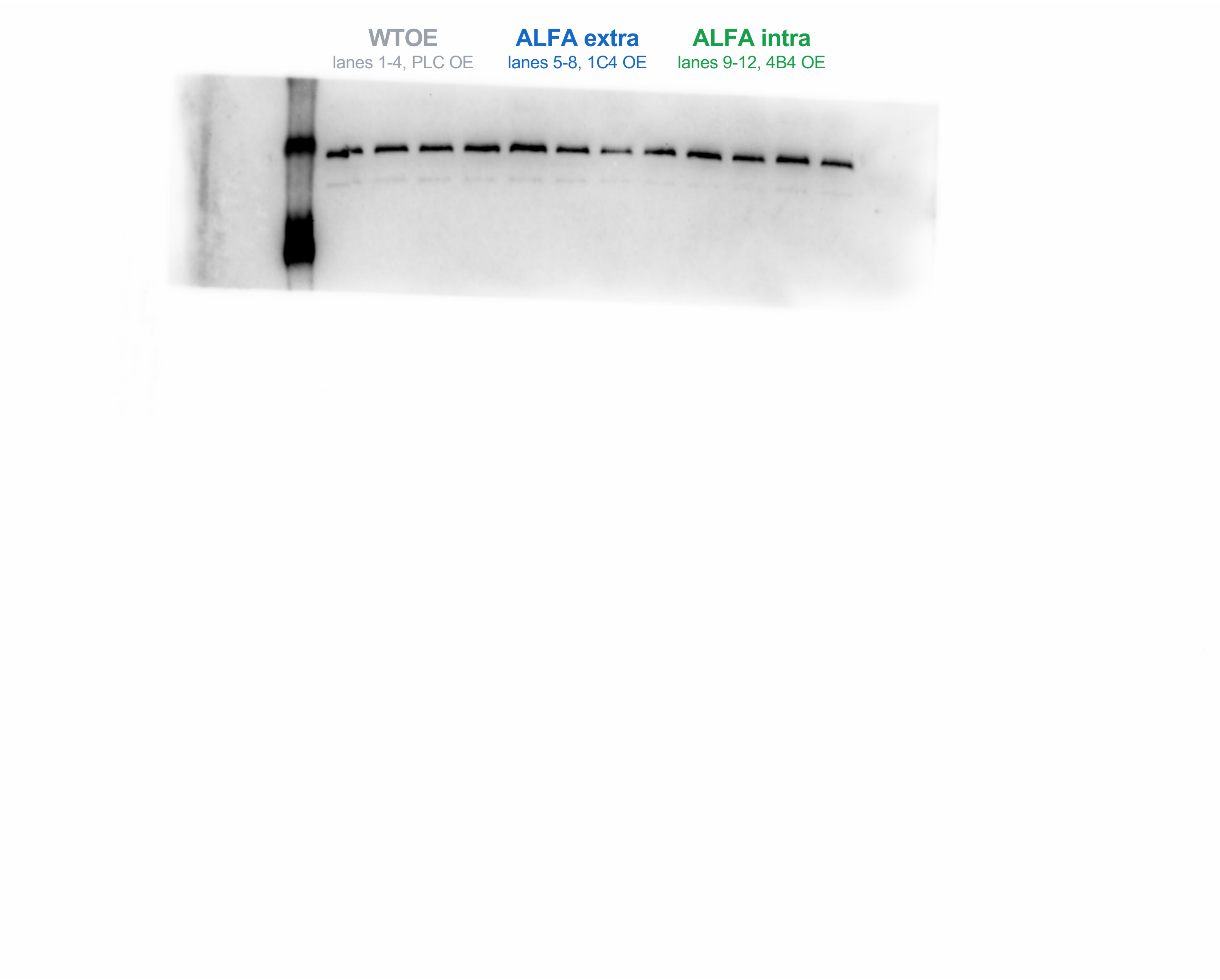

**Data S2. Uncropped immunoblots**

#### Figure 5E — ATP6V1A IP, whole membrane (uncropped)

2025-04-24 Chemiluminescence 550.778 s exposure 1375x1100 px 16-bit scan, linearly rescaled for display (39535-64990 of 0-65535); applied to the whole image.

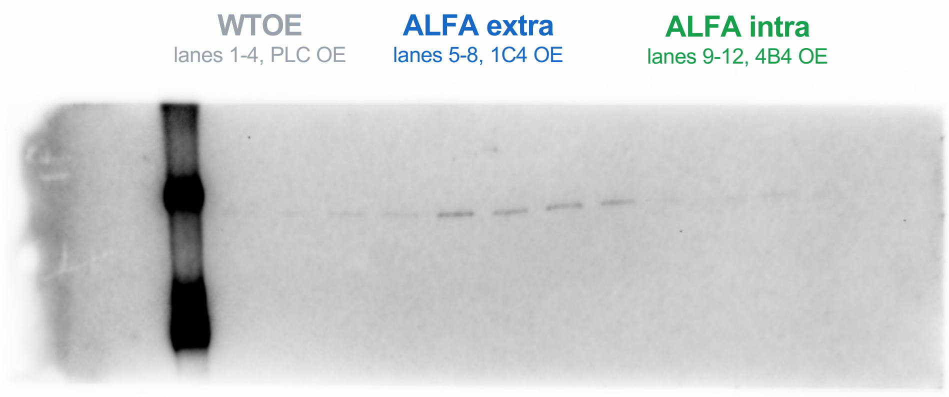

**Data S2. Uncropped immunoblots**

#### Figure 5E — UMPS input, whole membrane (uncropped)

1375x1100 px  16-bit scan, linearly rescaled for display (48170-65003 of 0-65535); applied to the whole image.

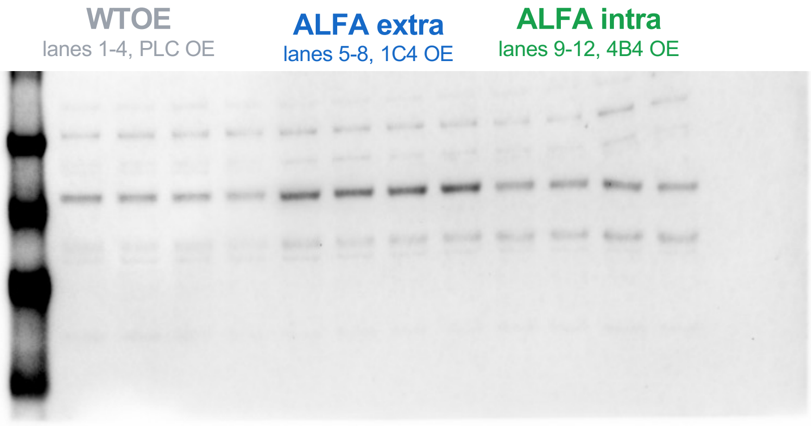

**Data S2. Uncropped immunoblots**

#### Figure 5E — UMPS IP, whole membrane (uncropped)

1375x1100 px  16-bit scan, linearly rescaled for display (44552-65017 of 0-65535); applied to the whole image.

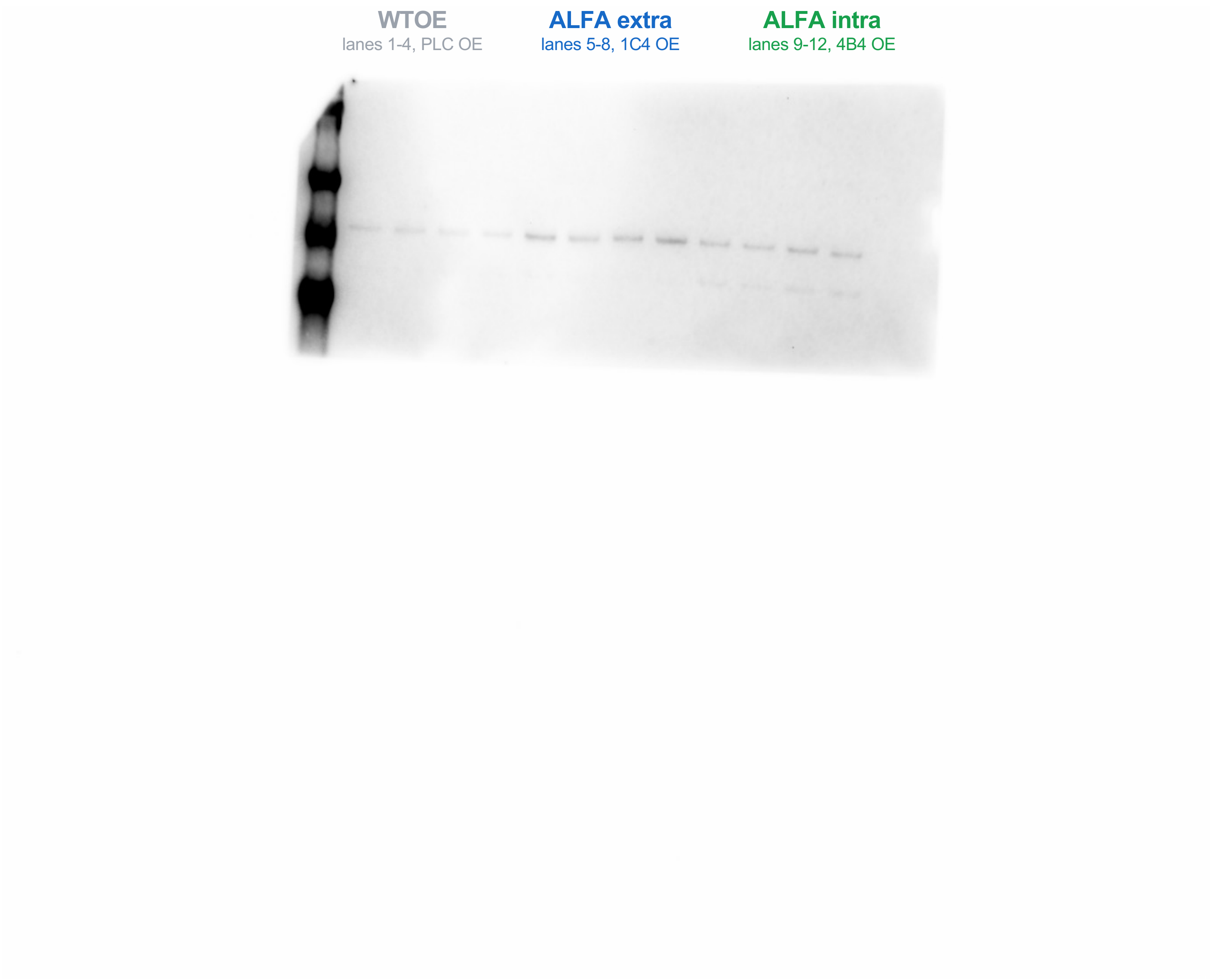
